# Modeling minimum viable population size with multiple genetic problems of small populations

**DOI:** 10.1101/2021.08.02.454753

**Authors:** Peter Nabutanyi, Meike J. Wittmann

## Abstract

An important goal for conservation is to define minimum viable population (MVP) sizes for long-term persistence. Although many MVP size estimates focus on ecological processes, with increasing evidence for the role of genetic problems in population extinction, conservation practitioners have also increasingly started to incorporate inbreeding depression (ID). However, small populations also face other genetic problems such as mutation accumulation (MA) and loss of genetic diversity through genetic drift that are usually factored into population viability assessments only via verbal arguments. Comprehensive quantitative theory on interacting genetic problems is missing. Here we develop eco-evolutionary quantitative models that track both population size and levels of genetic diversity. Our models assume a biallelic multilocus genome whose loci can be under either a single or interacting genetic forces. In addition to mutation-selection-drift balance (for loci facing ID and MA), we include three forms of balancing selection (for loci where variation is lost through genetic drift). We define MVP size as the lowest population size that avoids an eco-evolutionary extinction vortex after a time sufficient for an equilibrium allele frequency distribution to establish. Our results show that MVP size decreases rapidly with increasing mutation rates for populations whose genomes are only under balancing selection, while for genomes under mutation-selection-drift balance, the MVP size increases rapidly. MVP sizes also increase rapidly with increasing number of loci under the same or different selection mechanisms until a point is reached at which even arbitrarily large populations cannot survive anymore. In the case of fixed number of loci under selection, interaction of genetic problems did not necessarily increase MVP sizes. To further enhance our understanding about interaction of genetic problems, there is need for more empirical studies to reveal how different genetic processes interact in the genome.

## Introduction

In conservation management, the desire to protect threatened populations or species given limited available resources gave birth to the concept of a minimum viable population (MVP) size (Shaffer, 1981). Minimum viable population size is commonly defined as the lowest size that leads to population persistence for a specified duration with a set persistence probability, e.g., persistence for at least 1000 years with 99% probability or persistence for at least 50 generations with a probability of 90% (Shaffer, 1981; Traill et al., 2007). Alternatively, MVP size can be defined as the lowest size above which a substantial increase in extinction time occurs (Reed & Bryant, 2000) or in terms of fitness thresholds instead of persistence probability (Reed, 2005). So far, the choice of time and probability thresholds is arbitrary, varies among researchers, and across populations (Traill et al., 2007). This lack of consensus can make it hard to quantify relative extinction risk among different populations.

The viability of a small population and thus its MVP size depends on various ecological and genetic processes. Relevant ecological processes include demographic and environmental stochasticity, as well as Allee effects, e.g. through mate or pollen limitation at low density. In this study, however, we focus on the genetic processes shaping population viability and MVP sizes. In small populations, genetic problems such as inbreeding depression (ID), mutation accumulation (MA), and loss of genetic variation are inevitable (Lynch et al., 1995; Higgins & Lynch, 2001; Estes et al., 2004; Frankham, 2005; Coron et al., 2013). For example, increased homozygosity of deleterious alleles leads to a decrease in various fitness components (Spielman et al., 2004a; Bensch et al., 2006; Blomqvist et al., 2010; Palomares et al., 2012). Generally, genetic diversity is known to positively correlate with population size and influences population fitness (Reed & Frankham, 2003; Reed, 2005; Leimu et al., 2006; Hensen & Wesche, 2006). For instance, over 70% of threatened species have lower heterozygosity than their taxonomically related non-threatened taxa (Spielman et al., 2004b). However, diversity at neutral marker loci and high population size are not always a good predictor of population fitness (Yates et al., 2019; Teixeira & Huber, 2021). Other factors such as the genetic architecture of the population may have to be invoked for a better prediction of population fitness (Grossen et al., 2020; Kyriazis et al., 2021; Kardos & Luikart, 2021). But all in all, even if it is not always clear what aspects of diversity are most important and how they should be measured, there is increasing evidence for the importance of genetic diversity in population extinction (Frankham, 2005).

To date, inbreeding depression is the only genetic problem of small population size that is routinely included in MVP studies, e.g. those that employ the software package *VORTEX* (Lacy, 1993). Thus, other genetic problems such as mutation accumulation and loss of genetic diversity through drift are ignored although they are known to be important contributors to extinction risk (Frankham, 2005). In nature, these problems may occur simultaneously in a small population. However, to the best of our knowledge, no studies exist that quantify the expected interaction of the different genetic problems. At best, the interaction is verbally projected to increase extinction risk.

Another key factor for long-term population persistence is how genetic diversity is maintained. It is now clear that both mutation-selection-drift equilibrium and balancing selection mechanisms maintain genetic variation in populations (e.g., Hughes & Nei, 1988; Sharp & Agrawal, 2018), but balancing selection is still ignored in MVP studies. Balancing selection comes in various forms such as heterozygote advantage (HA), negative frequency-dependent selection (FD), and fluctuating selection with dominance reversals (FS), among others. The different forms of balancing selection can act in isolation or in some combination (e.g., Glémin et al., 2005; Hedrick, 2012; Bergland et al., 2014; Chen et al., 2015; Chapman et al., 2019; Mérot et al., 2020).

Our study has thus three goals: 1. to develop methods to account for genetic problems other than inbreeding depression in MVP studies, including loss of genetic variation at loci under balancing selection, 2. to quantify the effect of the interaction of two or more genetic problems on extinction risk, and 3. to find ways to reduce the arbitrariness of time and persistence probability thresholds in MVP analyses. To this end, we developed eco-evolutionary models using three approaches, i.e., individual-based model (IBM), diffusion approximation, and equilibrium-based Markov chain approaches. We defined MVP size as the minimum size below which an eco-evolutionary extinction vortex occurs (Nabutanyi & Wittmann, 2021). An eco-evolutionary vortex occurs when the percapita population growth rate turns negative. For the stochastic model, we used 95% persistence probability, but lower or higher probabilities did not substantially affect our MVP size estimates. To reduce the arbitrariness of time thresholds used in most definitions of MVP size, we either estimated MVP size based on time to attain genetic equilibria or even used a long-term stable equilibrium perspective.

### General Model

Our models considered a diploid hermaphroditic population initially at carrying capacity *K* with discrete-time population growth and non-overlapping generations. We assumed a genome with multiple unlinked bi-allelic loci. The genome was acted on by either one selection mechanism at all loci or multiple mechanisms simultaneously, each at a set of loci. We developed three different approaches to quantifying minimum viable population size (Fig. 1). The time-iterating approaches (one stochastic, one deterministic) tracked absolute fitness *W* and population size *N* until either the population went extinct or a set maximum number of generations (in this study, 4000 generations) was reached. The MVP size was the lowest population size needed for at least 1 individual to survive at the end of the simulation. For the stochastic IBM, this condition had to be satisfied in at least 95% of the replicates. We also used a time-independent approach that assumed a population at equilibrium with a stationary distribution of allele frequencies and constant population size. Here we defined MVP size as the minimum size required for absolute fitness *W* at equilibrium to be at least 1.

**Figure 1:**
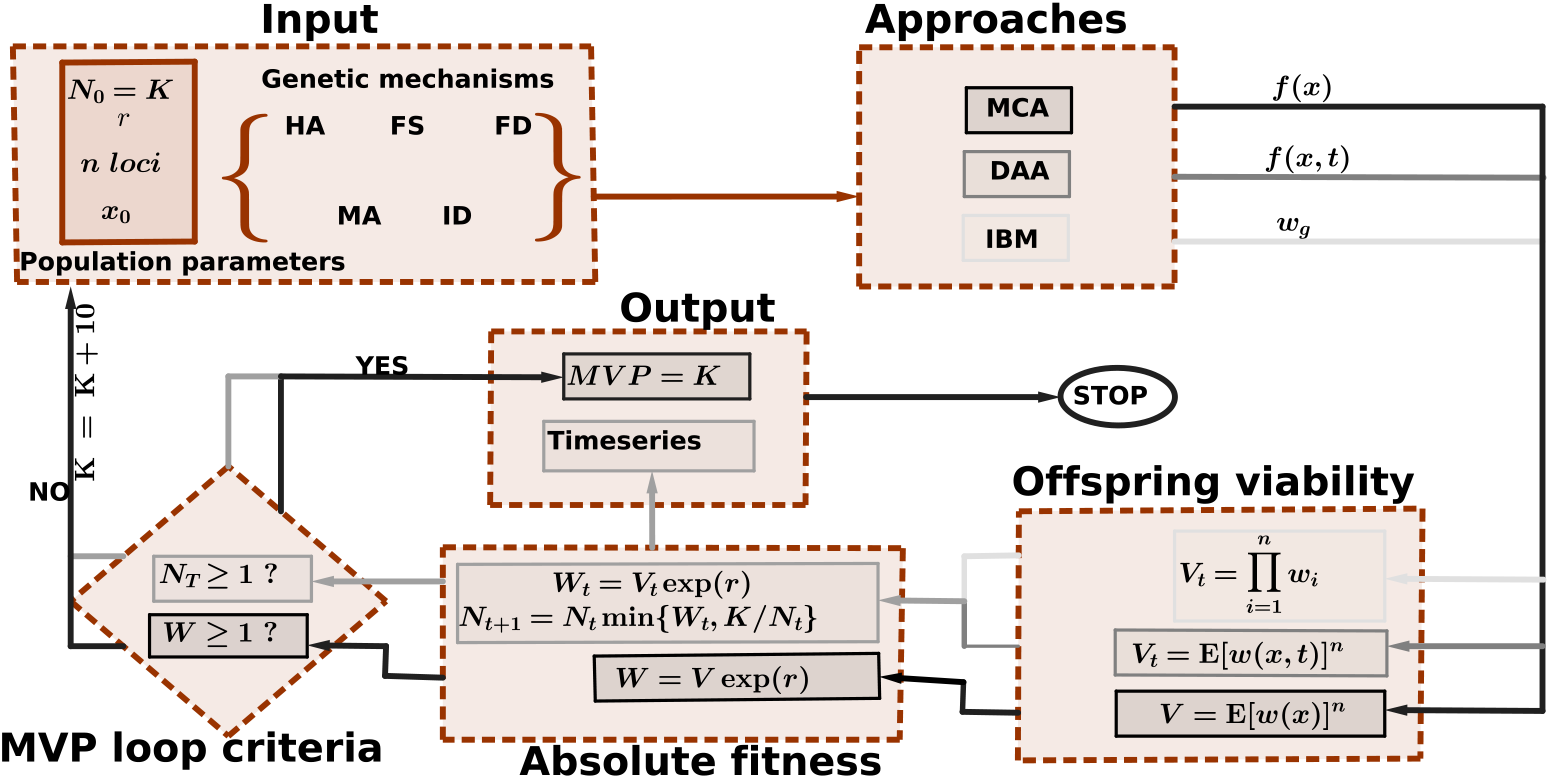
Overview of the three approaches to quantifying minimum viable population (MVP) sizes. For each population, we start with the required population parameters such as carrying capacity *K*, which also acts as the initial population size *N*_0_, intrinsic growth rate *r*, the number of loci on the genome *n*, and initial allele frequency *x*_0_. We then chose which selection mechanism(s) acts on the genome and simulated the population via either time-iterating approaches (i.e., diffusion approximation approach (DAA) or individual-based model (IBM)) or the equilibrium-based Markov chain approach (MCA). In the IBM, we tracked genotypes (and fitness) of each individual at each locus, and in DAA, we tracked allele frequency distribution *f*(*x, t*) from which offspring viability *V_t_*, absolute fitness *W_t_*, and population size *N_t_* were then determined. The iterations stopped when either the population went extinct or the maximum pre-set number of generations *T* was reached (whichever came first). In the MCA, we calculated the transition matrix for the number of allele copies at a constant population size from which the equilibrium allele-frequency distribution *f*(*x*), viability *V*, and absolute fitness *W*, were sequentially determined. Starting with *K* = 10, we incremented *K* by 10 in the MVP loop until either the MVP criteria was satisfied or *K* > 150000 (whichever came first).

### Selection Mechanisms

We considered five selection mechanisms, i.e., fluctuating selection with dominance reversals (FS), negative frequency-dependent selection (FD), heterozygote advantage (HA), inbreeding depression (ID), and mutation accumulation (MA). For HA, ID, and MA, the relative fitnesses of the three genotypes *AA, Aa*, and *aa* at a given locus had the general form *w_AA_* = 1 – *s_A_*, *w_Aa_* = 1 – *h* · *s_a_*, and *w_aa_* = 1 – *s_a_*, with the three mechanisms differing in the magnitude of selection (*s_i_*) and dominance (*h*) coefficients (Fig. 2a). In HA, 0 < *s_A_*, *s_a_* < 1 and *h* = 0 such that the heterozygotes had higher relative fitness than either homozygote. For both ID and MA, *s_A_* = 0,0 < *s_a_* ≤ 1 and 0 ≤ *h* ≤ 1. Thus, allele *A* was considered the wild-type allele whose homozygote had a maximum relative fitness while *a* was the deleterious allele. The difference between our example scenarios for ID and MA arose in the magnitude of *s_a_* and *h*, i.e., small *h* and potentially large *s_a_* for ID while larger *h* and smaller *s_a_* for MA (Fig. 2).

**Figure 2:**
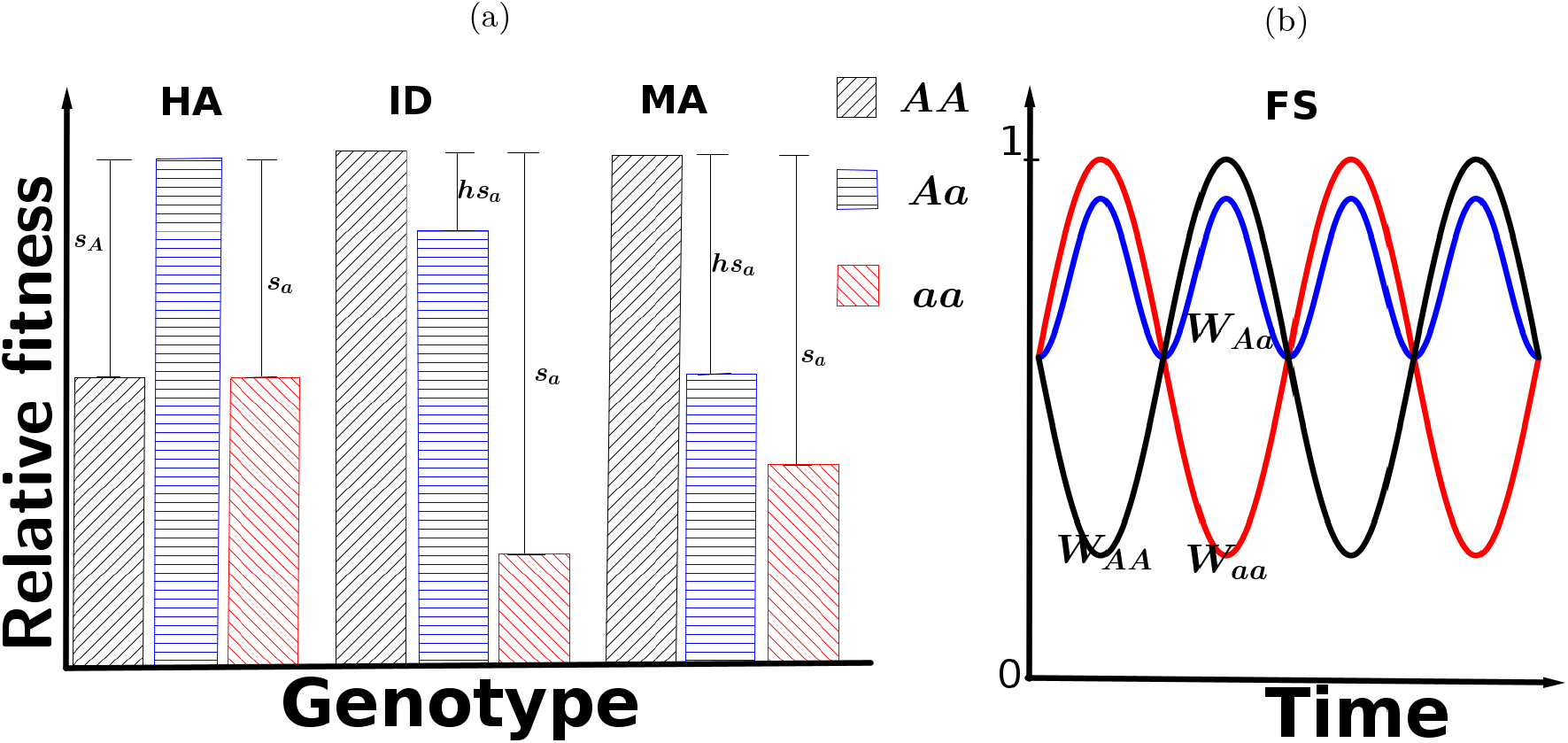
Relative fitness of the genotypes under different selection mechanisms. (a) For heterozygote advantage (HA), inbreeding depression (ID), and mutation accumulation (MA), the relative fitness is constant through generations. (b) For fluctuating selection with dominance reversals (FS), the relative fitness temporally fluctuates cyclically. We implemented this mechanism such that the heterozygote genotype is not the fittest in every generation but in any complete cycle of *κ* generations (we arbitrary used *κ* = 50), it emerged with a higher relative geometric mean fitness than either homozygote.

For FS and FD, the relative fitnesses of genotypes fluctuated over time. In FS, the fluctuations depended on the temporal changes in selection coefficients (Fig. 2b and Appendix S1). With FD, the fluctuations depended on allele frequency whereby a genotype with a rarer allele was more fit. Thus, we set *w_AA,t_* = 1 – *s_A_x_t_, w_aa,t_* = 1 – *s_a_*(1 – *x_t_*) and 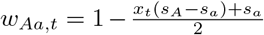, where *s_i_* was the maximum selection coefficient against the respective homozygote and *x_t_* is the frequency of allele *A*.

Three of the genetic mechanisms (HA, FS, and FD) give rise to balancing selection such that in an infinite population, both alleles can be maintained even without mutation. For these mechanisms, the mutation rates were assumed to be symmetric, i.e., *μ_A_* = *μ_a_*, where *μ_A_* is the mutation rate for allele *A* to allele *a* and vice versa for *μ_a_*. On the other hand, ID and MA do not yield balancing selection and genetic diversity at loci under these mechanisms is due to the mutation-selection-drift equilibrium. Here we assumed that back mutations are very rare and set 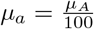 for both ID and MA.

### Individual-Based Model (IBM) Simulations

The initial population size was drawn from a Poisson distribution with mean *K*. Each initial individual was randomly assigned two gene copies at each of the *n* loci. The initial *A* allele frequency was 0.5 for all balancing selection mechanisms and 0.95 for ID and MA.

In each generation, each parent produced a random number of offspring drawn from a Poisson distribution with mean *E* = *e^r^*, where *r* is the intrinsic per-capita growth rate. The focal parent randomly chose a mating partner from all the available individuals including itself as a second parent for each offspring independently. For each locus, each partner independently and randomly passed on one of the available alleles to the respective offspring locus. Finally, the offspring alleles mutated with probability *μ_A_* or *μ_a_*.

Assuming multiplicative fitness across loci, each offspring was viable, i.e. survived, with probability, 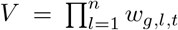, where *w_g,l,t_* denotes the fitness of genotype *g* ∈ {*AA, Aa, aa*} at locus *l* and generation *t*. After all individuals had reproduced, the surviving offspring replaced the parent generation. If there were more than *K* surviving offspring, only *K* offspring were randomly selected to replace the parents.

Each simulation run was iterated until either the population went extinct or generation 4000 was reached. This time was long enough for stationary allele frequency distributions to be attained under most selection scenarios except at very low mutation rates in ID and MA (see Results and Discussion sections).

### Deterministic Approximations to the Individual-Based Model

There were two main steps in the deterministic approximations, namely, approximating the distribution *f* of allele frequency *x* and determining the absolute fitness *W* using *f*. For the first step, *f* was determined by either the diffusion approximation approach (DAA) or the Markov chain approach (MCA).

The diffusion approximation involved numerically solving the diffusion equation

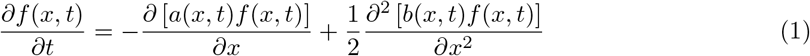

for *f*(*x, t*) in each generation subject to some initial and boundary conditions (Zhao et al., 2013; Xu et al., 2019; Nabutanyi & Wittmann, 2021), where *a*(*x,t*) and *b*(*x,t*) are the mean and variance of the change in allele frequency during an infinitesimal time interval, derived for each selection mechanism (Appendix S2.1).

For the MCA, we assumed constant population size *N* and computed the transition probabilities of allele A from *i* copies at time *t* to *j* copies at time *t* +1 (*i, j* ∈ {0,1, 2, ⋯, 2*N*}). Assuming random sampling, the number of copies of the A allele in the next generation follows a binomial distribution with parameters 2*N_t_* and *x*″ (the allele frequency after the action of selection and mutation). Due to numerical problems at large *N* (*N* > 500) with the binomial distribution, we approximated the transition probabilities using the Poisson distribution (when one of the alleles is rare) or the normal distribution (Appendix S2.2). Given the transition probability matrix, we then obtained the allele frequency distribution *f*(*x*) as the eigenvector corresponding to the largest eigenvalue of the transition matrix. Appendix S2.2.1 also shows how MCA can be made time-dependent.

In the second step, we began by estimating the mean offspring viability *V* from *f*(*x*). We let 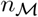 be the number of loci under genetic mechanism 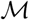. Then, assuming multiplicative fitness across loci and independence between loci, the fitness from 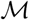, is given by

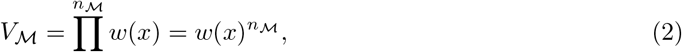

where *w*(*x*) is the fitness at allele frequency *x* at a specific locus given by

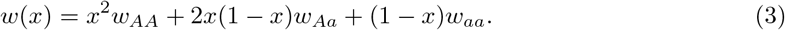

Because the exact allele frequency *x* at a given locus is not explicitly known, we calculated 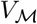 using the expected value of *w*(*x*) under the allele-frequency distribution:

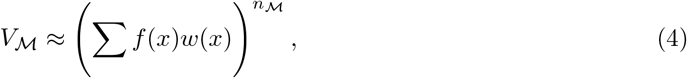

where the sum is over all allele-frequency bins. Also, assuming independence between selection mechanisms, then, the offspring survival fitness 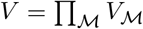.

The second step was completed by computing the absolute fitness, *W* = *VE*. Note that for DAA, the distribution *f* is calculated for every generation and so is 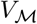 and *W*. Also, the population size *N*_*t*+1_ was determined from the previous generation *t* as

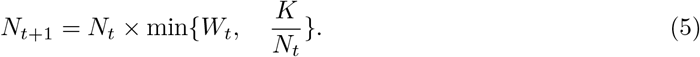

## Results

In this section, we focus on results from HA, FS, ID, and MA. The results from FD are to a large extent similar to those from HA (see supplement Appendices S3 to S7).

For both isolated and interacting selection mechanisms, the absolute fitness initially decreased while the population remained at the carrying capacity *K* (Fig. 3, Appendix S3.1). For relatively high *K*, absolute fitness decreased to an equilibrium value of at least 1, while populations remained at *K*. However, at lower *K*, absolute fitness decreased below a minimum threshold of 1 (gray line on Fig. 3 row 2) while the populations remained at *K*, before rapidly decreasing to extinction at the time when absolute fitness just fell below 1. For scenarios with FS, there were expected periodic fluctuations in both population size and absolute fitness, and the absolute fitness fell below 1 in some generations without causing population extinction (Fig. 3 column 4). The lowest carrying capacity that avoided population extinction was taken as the MVP size (indicated in red, Fig. 3 row 1). The MVP size estimate from scenarios that involved ID (and MA) increased with increasing simulation time (Appendix S3.1.1)

**Figure 3:**
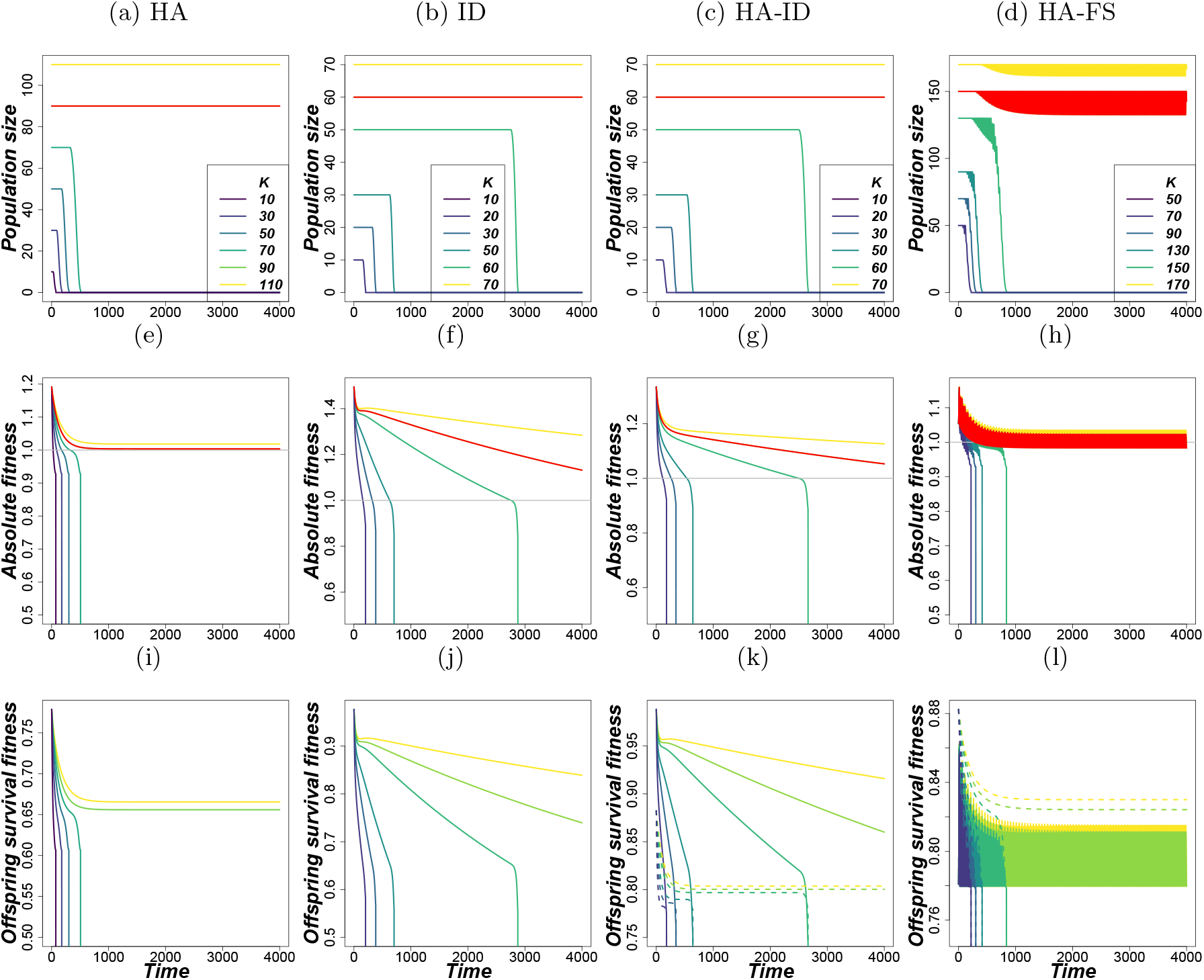
Example trajectories under various carrying capacities for; Column 1: HA, Column 2: ID, Column 3: HA-ID and Column 4: HA-FS while for Row 1: population size, Row 2: absolute fitness and Row 3: offspring viability fitness for each constituent mechanism. The trajectories were obtained using the diffusion approximation approach (see Appendix S3.2 for some IBM trajectories). The gray horizontal line is the threshold absolute fitness of 1. For balancing selection, *s_a_* = *s_A_* = 0.005, *μ_A_* = *μ_a_* = 0.0005 and for ID *s_A_* = 0, *s_a_* = 0.05, *h* = 0.01, *μ_A_* = 0.0005, 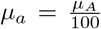,. Other shared parameters are *n* = 100, *K* = *N*_0_, *r* = ln(1.53). For scenarios with two selection mechanisms (columns 3 and 4), 50 loci were under each mechanism. For population trajectories from other scenarios, see Appendix S3.

In cases involving more than one selection mechanism, the individual mechanisms reduced maximum offspring viability (which is 1) by different amounts that varied through time (Fig. 3, row 3). The rate at which locus fitness decreased (and attained equilibrium) also differed between constituent mechanisms.

In Fig. 4, we simulated populations with different number of loci under selection. We first varied the number of loci under selection between 1 and 100 for each mechanism separately. Thus, genomes with at most 100 loci under selection were acted on by one selection mechanism. The additional loci were either under the same mechanism or one of the other three selection mechanisms. The MVP size initially remained constant at 10 (our lowest tested population size) as the number of loci increased until a certain number was reached where a rapid increase was registered. Even under the same selection mechanism, the rate at which log MVP size increased decreased after a certain number of loci. As more loci were added, log MVP size again increased rapidly, producing an inflection point. Note though that on a linear scale, MVP size increases more and more rapidly with increasing number of loci and there is no inflection point, except for MA which according to the MCA mechanism still exhibits an inflection point (Appendix S4.3). At a certain critical number of loci, MVP sizes exhibited an asymptotic increase (to infinity), indicating that no finite population size was sufficient to keep the population at equilibrium. Note that we tested *K* values from 10 to 150000 (using DAA) and numbers of loci from 1 to 1000 for all the scenarios. Unlike for ID and MA, the critical number of loci required for an asymptotic increase in MVP size was independent of mutation rates for balancing selection mechanisms (Appendices S4 and S4.3). Also, MCA produces higher MVP estimates than DAA in some cases of ID and MA scenarios, but the divergence disappears as more loci are added.

**Figure 4:**
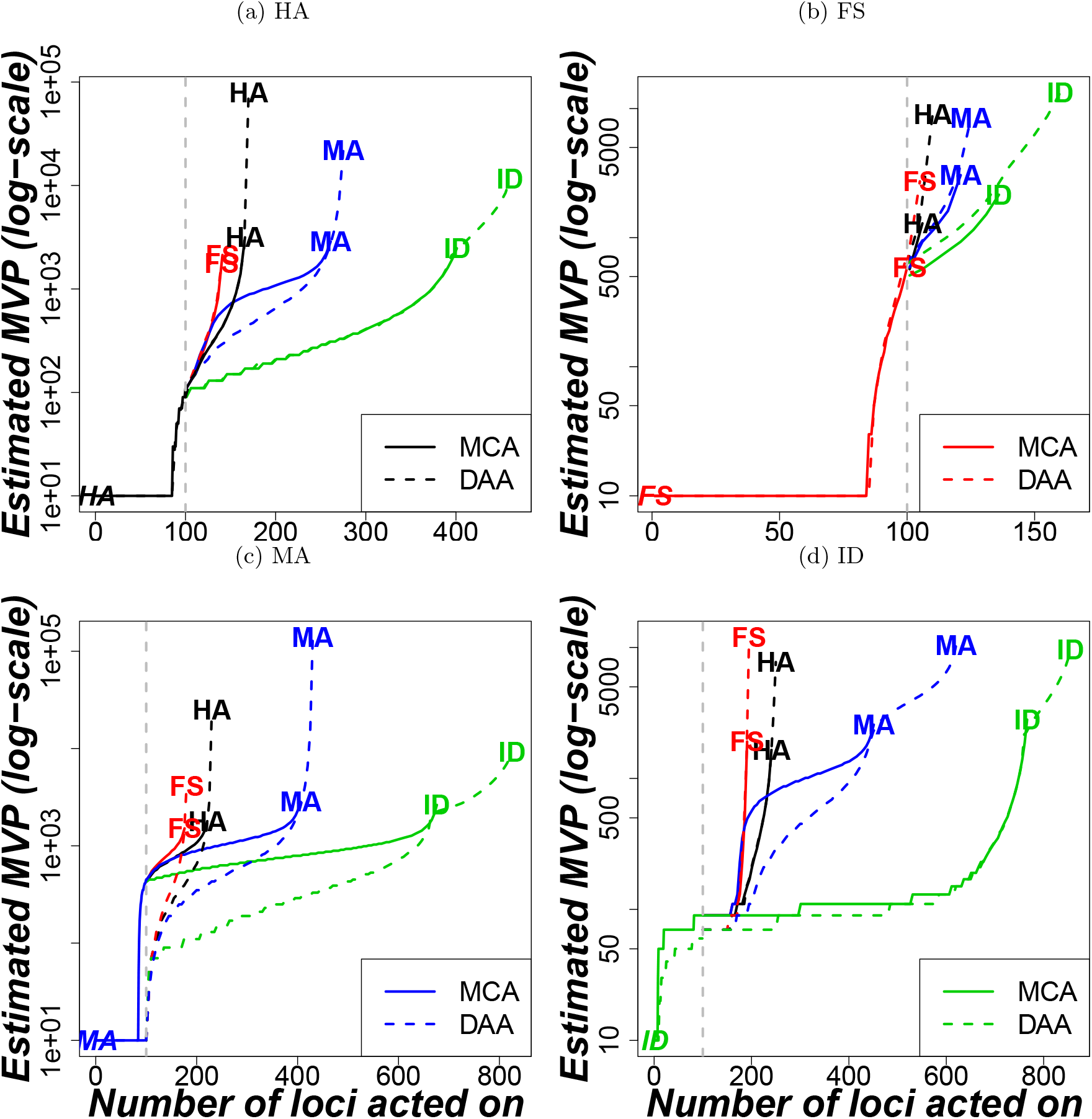
Effect of increasing number of loci under selection on MVP size. The genome from populations with at most 100 loci (below the vertical gray line) was acted on by the same selection mechanism (indicated at the start of the line) (a) HA, (b) FS, (c) ID and (d) MA. Those genomes with above 100 loci, the mechanism indicated at the end of the line acted on the remaining loci, e.g., the line with HA—ID label implies the first 100 loci were acted on by HA while ID acted on the remaining loci. See Appendix S4.3 for results from more scenarios. Parameters for balancing selection were *s_a_* = *s_A_* = 0.005, *μ_A_* = *μ_a_* = 0.0005, for ID and MA, *s_A_* = 0, 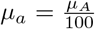 while *h* = 0.01, *s_a_* = 0.05 for ID and *h* = 0.5, *s_a_* = 0.005 for MA. Shared parameters were *K* = *N*_0_, *r* = ln(1.53).

The effect of mutation rate on MVPs varied strongly among genetic mechanisms (Fig. 5). For genomes that were acted on by only balancing selection mechanisms (HA, FS) either separately or simultaneously, the MVP size estimates generally decreased rapidly with mutation rates (Fig. 5). By contrast, MVP estimates from genomes acted on by either ID, MA, or both increased rapidly.

**Figure 5:**
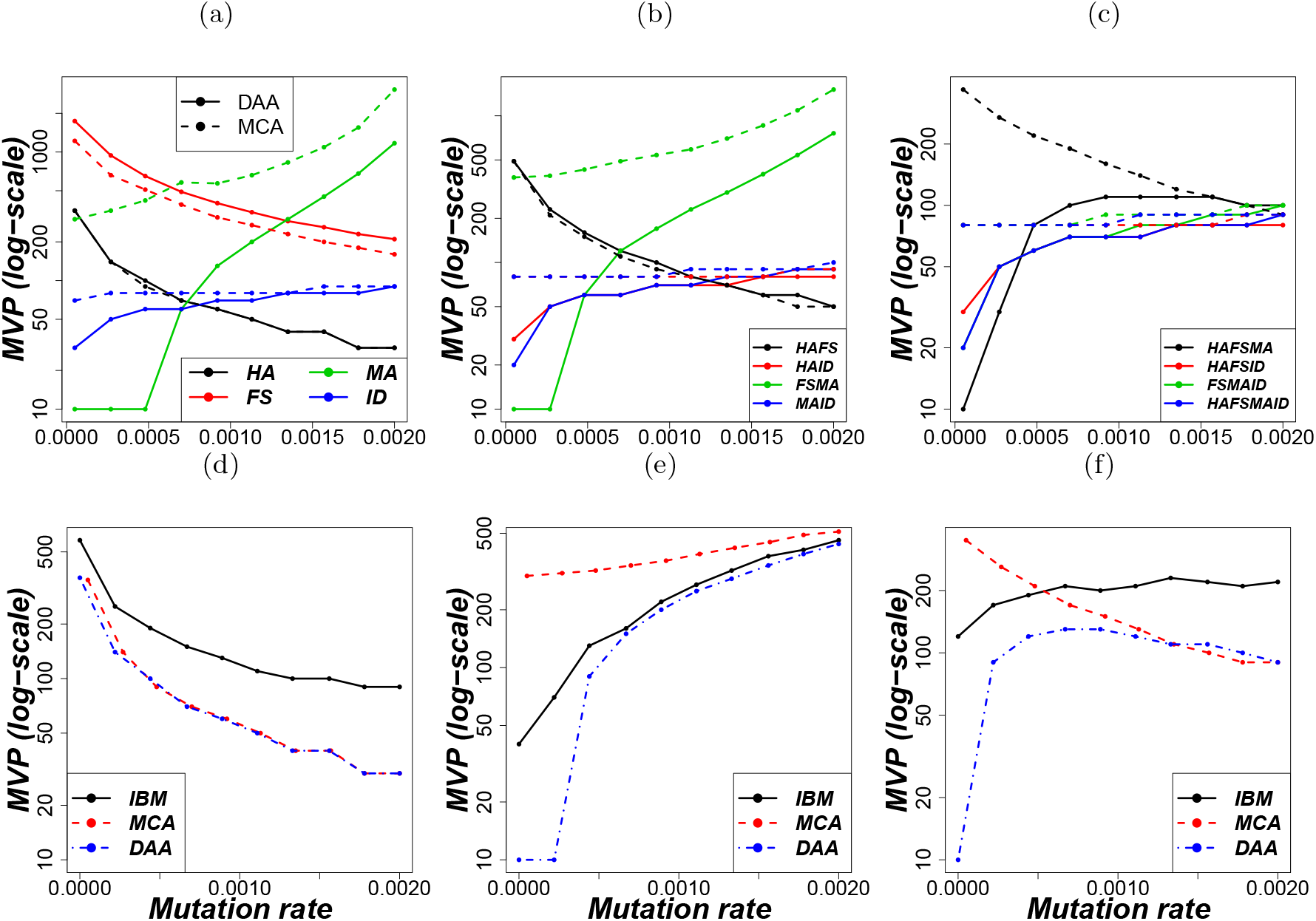
Comparison of MVP size estimates across approaches with varying forward mutation rate, *μ_A_*. Row 1: Results obtained from the Markov chain approach (MCA; dashed lines) and the diffusion approximation (DAA; solid lines). Genomes under (a) isolated mechanisms, (b) paired mechanisms and (c) three and four mechanisms. Row 2: Comparison of MVP size estimates from the IBM, DAA and MCA. (d) HA only, (e) ID only and (f) HA-FS-FD-ID-MA. The estimates from IBM assumed 95% survival probability of 60 replicates. In both rows, parameters for balancing selection were *s_a_* = *s_A_* = 0.005, *μ_A_* = *μ_a_*, and for ID and MA *s_A_* = 0, 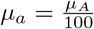. ID and MA differed in *h* and *s_a_* with row 1: *h* = 0.5, *s_a_* = 0.005 for MA and *h* = 0.01, *s_a_* = 0.05 for ID while row 2: *s_a_* = 0.005 for both ID and MA but *h* = 0.5 for MA and *h* = 0 for ID. Other shared parameters were *K* = *N*_0_, *r* = ln(1.53). The total number of loci *n* on the genome is 100 except for combinations with 3 mechanisms where it is 99 for purposes of equal distribution of loci between mechanisms.

With a fixed total number of loci under selection, in genomes with more than one mechanism, the joint MVP size was in most cases closer to the harmonic (or geometric) mean of the MVP estimates obtained when the constituent selection mechanisms acted in isolation than to the arithmetic mean (Appendix S5.3). However, there were also exceptions where the joint MVP size was close to or even larger than the arithmetic mean of single MVP sizes.

In most scenarios that contained only balancing selection, the estimates from DAA and MCA roughly coincided. However, as mutation rates decrease in ID and MA, there were substantial quantitative differences at lower mutation rates between MVP estimates from DAA and MCA (Fig. 5 row 1). The IBM estimates (Fig. 5 row 2) were (slightly) higher than estimates from DAA, and converged better to the MCA and DAA estimates at higher MVP sizes. Also, the IBM estimates were fairly insensitive to levels of survival probability thresholds (Appendix S5.1).

## Discussion

Small populations can be faced with multiple genetic problems simultaneously. In this study, we quantified the effect of such interactions on extinction risk by estimating MVP sizes. MVP size rapidly increased as more loci under selection were added. However, when the total number of loci in the genome was fixed, MVP size estimates were in most but not all of the scenarios we considered dominated by the mechanism that yielded the lowest estimate when acting alone on the genome.

### Number of loci under selection

As the number of loci under selection increased, cumulative selection pressure on the genome increased and as a result, offspring viability decreased, thus increasing MVP size. Under balancing selection mechanisms, the critical number of loci above which no arbitrary population size survived was independent of mutation rates and decreased with increasing selection coefficients (Appendix S4). This is because, in an infinite population under symmetric balancing selection and symmetric mutation rates, equilibrium fitness is independent of mutation rates. Moreover, heterozygosity increases with increasing mutation rate, which increases population fitness. Thus MVP estimates are lower at higher mutation rates. By contrast, the critical number of loci under mutation-selection-balance was independent of the selection coefficient but decreased with an increase in mutation rates. This is in agreement with classical studies showing that equilibrium fitness only depends on mutation rate (*μ_A_*) and not on selection coefficient (*s_a_*) if *μ_A_* << *s_a_* (Haldane, 1937; Agrawal & Whitlock, 2012; Felsenstein, 2015).

Most of the predominance of the mechanism that yielded the lowest MVP estimate in the fixed number of loci scenario can be explained by the relationship between viability per locus and population size. In Appendix S6, we assume that viability increases linearly with population size, and show that the joint MVP is approximately equal to the harmonic mean of the MVPs obtained from the constituent mechanisms when acting in isolation. We also illustrate scenarios where the joint MVP can be even higher than the harmonic, geometric, and arithmetic means as also observed in some cases in our results.

### DAA, MCA, and IBM

In many regions of parameter space, the three approaches largely agreed in their MVP size estimates both qualitatively and quantitatively, but there were also discrepancies and each approach has its advantages and disadvantages. The relatively low MVP estimates from the iterating approaches in ID and MA at low mutation rates can be attributed to lack of attainment of a stationary allele frequency distribution within the simulated period of time. For example, for mutation rate of *μ_A_* = 0.00005 and 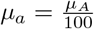, equilibrium is attained after at least 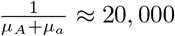 generations from mutation alone (see Felsenstein, 2015). The equilibrium-based MCA estimates did not depend on the length of the simulation period and the initial conditions. Therefore, MCA estimates can be considered as the ”true” MVP estimates if one is interested in the possibility of a long-term stable population. However, MCA may give overly optimistic predictions for population survival if the population considered starts far from genetic equilibrium (Appendix S7; see also Kardos & Luikart, 2021). For example, during founder events, population fragmentation, and bottlenecks, deleterious allele frequencies may be above equilibrium values such that populations may be unable to replace themselves in the early generations and go extinct before reaching a genetic equilibrium. The time-dependent approaches could account for such transient dynamics, but for this the initial genetic composition of the population needs to be known.

Unlike in the DAA and MCA approaches where allele frequencies evolve completely independently across loci, in the IBM evolution at one locus can potentially interfere with evolution at another locus, making the fitnesses of the genotypes at the focal locus more variable. IBMs also have the advantage that they take into account demographic stochasticity (which in some cases led to higher MVP size estimates than for the MCA and DAA approach). Therefore IBM gives a better representation of natural populations. However, IBMs are extremely time-consuming especially at higher *K* since many replicates are required to infer MVP sizes. Similarly, obtaining the transition matrix in the MCA is computationally impossible for higher *K*. On the other hand, the accuracy in DAA depends the discretization of allele frequency, *x* and can perform poorly for very small population sizes. Thus each of the approaches will be most useful in a different context: MCA to get long-term predictions for small populations, IBM for small populations when demographic stochasticity and inter-locus interactions should be accounted for, and DAA for large populations.

### Choice of the MVP Definition Thresholds

In the case of more than one threatened population, managers may have to prioritise allocation of available limited resources. Therefore, there is need to establish relative viability status of the populations e.g., by comparing required MVP sizes to current population sizes. Equilibrium-based MVP sizes could help make MVP sizes more comparable across populations, while still taking into account specific population parameters. In the IBM, our estimates were largely insensitive to survival thresholds (Appendix S5.1), possibly due to low sources of stochasticity in the model. We expect survival thresholds to be important in estimates that consider more ecological stochastic processes such as environmental stochasticity.

Furthermore, we chose an extinction threshold of 1 individual for the time-iterating approaches. However, larger thresholds could also be chosen for dioecious populations and populations that suffer from Allee effects due to mate-finding difficulties or other ecological problems at low population size. For the MCA with equilibrium assumptions, a quasi-extinction threshold of absolute fitness 1 was used. The absolute fitness of 1 acts as a boundary between population persistence (*W* > 1) and rapid population decline (*W* < 1) in an eco-evolutionary extinction vortex (Nabutanyi & Wittmann, 2021).

### Concluding Remarks and Recommendations

Since our models assumed an idealized randomly mating population, the estimates in this paper should be compared to the genetically derived effective population sizes in the literature such as the long-term 500 estimate in the 50/500 rule of thumb (Franklin, 1980; Frankham et al., 2014). Our estimates range from 10 to about 5000 before the critical number of loci is reached. Some studies have proposed that census MVP values would need to be up to 10 times higher than the effective MVP sizes (Frankham, 1995). This would translate our estimates to the range 100 to 50000, which generally lies in the range reported by most studies, (Harcourt, 2002; Reed et al., 2003; Traill et al., 2007; Wang et al., 2019).

In nature, both mutation-selection balance and balancing selection contribute to standing genetic variation (Sharp & Agrawal, 2018), potentially with the different genetic mechanisms playing a larger role at different life stages. For instance, variation in offspring viability traits may often be under mutation-selection balance while variation in traits linked to sexual reproduction may more often be under some form(s) of balancing selection (Sharp & Agrawal, 2018). Our observations were based on independence between loci. Epistasis and linkage disequilibrium may give different results. Therefore, to improve extinction risk assessment, a better understanding of the underlying genetic processes in a given population is needed. Also, balancing selection may be more common in natural populations than earlier proclaimed (Chapman et al., 2019; Mérot et al., 2020), and thus we suggest that it would make sense to incorporate balancing selection mechanisms into software tools designed to evaluate population persistence.

To better understand how interacting genetic problems affect population persistence, empirical studies are needed. There are many technical challenges in obtaining the necessary detailed genetic information, e.g., identifying which and how many loci are responsible for specific fitness traits, and which genetic mechanism(s) maintain variation at which loci. However, current studies are already heading in the right direction. For example, laboratory experiments with *Drosophila melonagaster* can now distinguish the contributions of different genetic mechanisms in maintaining variation at a given locus (Charlesworth, 2015; Sharp & Agrawal, 2018). In addition, modern sequencing techniques can allow quantitative trait loci to be mapped to different fitness components and genomic regions with traits that influence fitness components (such as survival and fecundity) can be identified from neutral quantitative trait loci (Ågren et al., 2013).

To conclude, our results revealed that if the number of loci under selection is fixed, a mixture of genetic processes is not necessarily worse than having the same genetic mechanism acting at all loci. However, if the total number of loci under selection increases, for example because genetic mechanisms that were previously neglected are now included, estimated extinction risk and MVP sizes can dramatically increase. These observations call for inclusion of all genetic processes in assessing the viability of populations. Populations normally also face other ecological problems such as environmental stochasticity and Allee effects, and their consideration is expected to further increase extinction risk and MVP size estimates (Tanaka, 2000; Wittmann et al., 2018).

## Supporting information

R codes used to simulate results

## Supporting Information

In the supplementary material, we briefly show the mathematical derivation for FS (Appendix S1). In Appendix S2, we provide more details about DAA and MCA. Appendices S3 through S7 provide some additional results that also involve FD. Also, we show how estimates of minimum viable population size depend on fitness per locus (Appendix S6) and how equilibrium fitness and critical number of loci can be estimated (Appendix S4).

## Appendices

### S1 Brief Description of Fluctuating Selection with Reversal of Dominance (FS)

Briefly, FS was derived as follows (see Nabutanyi & Wittmann (2021) for a detailed description). The relative fitnesses of the genotypes were 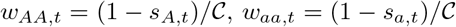 and 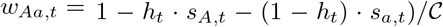, where *s_A,t_* = *s_A_* · sin(2π · *t/κ*) and *s_a,t_* = *s_a_* · sin(π + 2π · *t/κ*) are temporally fluctuating selection coefficients and *h_t_* = 0.5 – *c* · sin(2π · *t/κ*) is the temporally fluctuating dominance coefficient. Here, 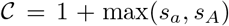 is the normalization constant such that the genotype fitness lies between 0 and 1, *s_A_* and *s_a_* determine the amplitude of selection and 0 ≤ *c* ≤ 0.5 determines the strength of dominance. We arbitrarily used *c* = 0.4 and *κ* = 50 throughout the paper, where *κ* is the number of generations in a complete cycle.

### S2 Diffusion Approximation and Markov Chain Approaches

#### S2.1 Diffusion Approximation Approach (DAA)

Here, *f*(*x,t*) at a given locus was approximated using the diffusion equation (e.g in Kimura et al. (1955))

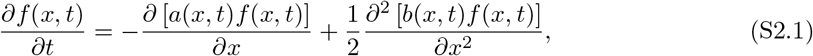

where *a*(*x, t*) and *b*(*x, t*) refer to the mean and variance of the change in allele frequency during an infinitesimal time interval. For all the mechanisms, we took

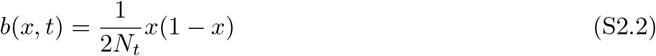

(e.g see Durrett (2008)) while *a*(*x,t*) depended on the mechanism. For example, assuming *s_a_* and *s_A_* are very small that higher order terms involving 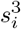 and above are negligible, for HA, ID and MA,

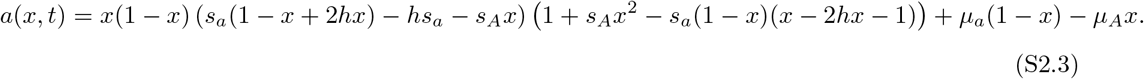

The three mechanisms differed in *h, s_a_* and *s_A_*. For FS (see Nabutanyi & Wittmann (2021) supplementary Section A.3),

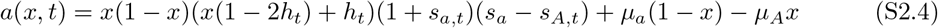

and for FD,

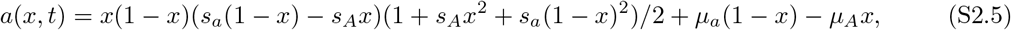

where, *h_t_*, *h*, *μ_a_*, *μ_A_*, *s_A_*, *s_a_*, *s_a,t_*, and *s_A, t_* are as defined under ”Selection Mechanisms” subsection.

We adopted the numerical scheme in Zhao et al. (2013); Xu et al. (2019) to solve Equation (1) subject to the initial condition *f*(*x*, 0) = *x*_0_ and the boundary condition that conserves probability at all times. The scheme basically discretizes both time *t* and allele frequency *x* over the interval 0 and 1 inclusive. The step-length for *x* is 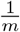, where *m* is an integer above 0 (used *m* = 20 for all the results) and step-length for time being *τ* > 0 (in the whole paper, we used *τ* = 1 so as to correspond to a full generation). The numerical scheme eventually gives a vector of probabilities *f*(*x_i_, t*) with *i* ∈ {0,1, ⋯, *m*} for each time point as the distribution *f*(*x, t*).

#### S2.2 Markov Chain Approach (MCA)

This approach involved finding the transition probabilities from *i* copies of a given allele at time *t* to *j* copies of the same allele at *t* + 1. Let *X* be a random variable representing the number of copies of allele *A* in a population of size *N_t_* at time *t*. In a diploid population with two alleles at a given locus, there are 2*N_t_* alleles in total, and *X* can take on {0,1, 2, ⋯, 2*N_t_*}. The allele copies in the next generation *t* + 1 were randomly sampled (with replacement) from those present at the current generation *t*. Therefore, *X* follows a binomial distribution with parameters 2*N_t_* and *x*″ (the allele frequency after selection and mutation acting). Due to computational failure of the binomial distribution for large *N_t_* > 500, we approximated the binomial distribution with one of two distributions, the Poisson distribution with parameter 2*x*″*N*_*t*+1_ (or 2(1 – *x*″)*N*_*t*+1_ when one of the alleles is rare (expected copy number < 35) (Fig. S2.1), and otherwise the normal distribution with parameters 2*x*″*N*_*t*+1_ and 2*x*″(1 – *x*″)*N*_*t*+1_ as mean and variance respectively. In summary, the probability of having *j* allele copies at time *t* +1 given that *i* copies are present at time *t* yields a transition matrix **P***_ij_*,

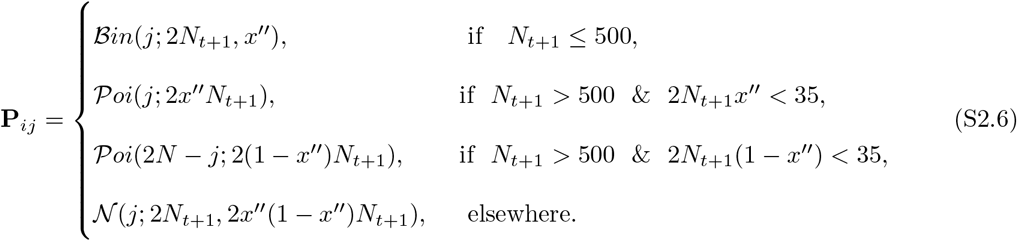

where 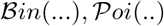 and 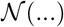 are the probability mass functions of the binomial, Poisson and normal distributions respectively. Here, *i* = {0, 1, ⋯, 2*N*_t_}, *j* = {0, 1, ⋯, 2*N*_*t*+1_} and *x*″ is the frequency of allele *A* at the end of a given generation *t* after possible selection and mutation, i.e., 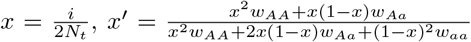 and *x*″ = (1 – *μ_A_*)*x*′ + *μ_a_*(1 – *x*′). We then normalise each row such that the transition probabilities always sum to 1.

Assuming a population at equilibrium with a constant size *N*, **P***_ij_* is a (2*N* +1) × (2*N* + 1) square matrix. The equilibrium distribution of allele copies was obtained by getting the eigenvector corresponding to the largest real part of the eigenvalues (which is always equal to 1) of matrix **P***_ij_*, which was then normalised to get *f*(*x,t*).

##### S2.2.1 Distribution of Allele Frequencies via Iterating through Generations for the Markov Chain Approach

Similar to the diffusion approximation, the MCA can also be made time-dependent by starting with some initial allele frequency distribution and evaluating the distribution in every generation (Section S2.2.1). Here, the transition matrix is not necessarily a square matrix and allows varying population sizes between generations.

Here, matrix **P**_*t*_ (subscript *t* due to change with time) is calculated every generation and is not necessarily a square matrix. **P** is generally a 2*N_t_* + 1 × 2*N*_*t*+1_ +1 matrix filled according to Equation (S2.6). At each generation *t* ≥ 0, the (row vector) distribution of allele copies *X* is determined as

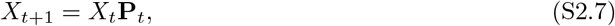

where we start with some initial allele frequency distribution *X*_0_. The distribution is used in a similar way as in the diffusion approximation to determine population size (Equation (5)). However, this approach is computationally more expensive than the DAA and equilibrium-based MCA.

**Figure S2.1:**
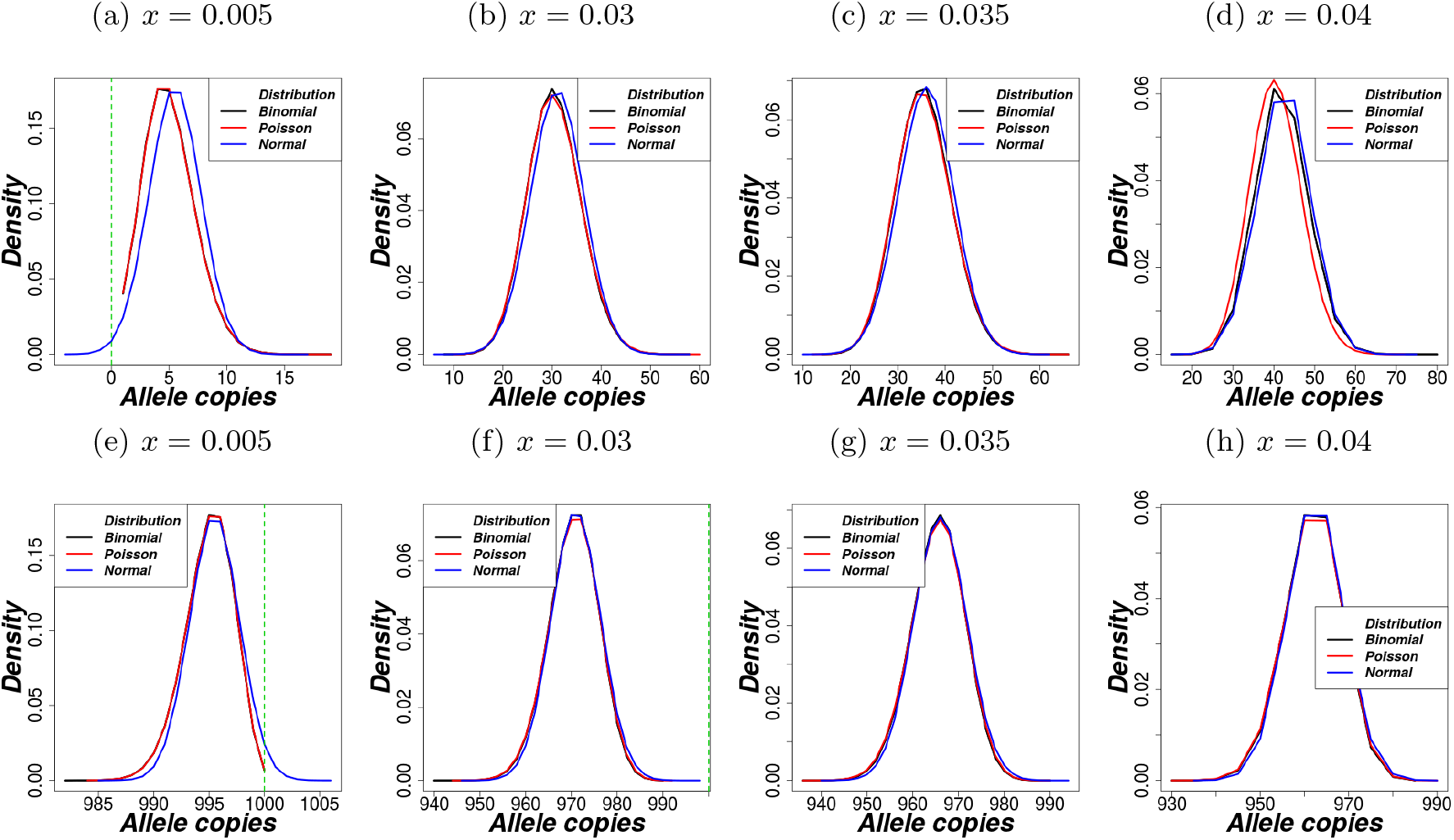
Densities for the distribution of allele frequencies with different values of probability, *x*. The upper panel is obtained from the binomial (*n* = 2*N*, *p* = *x*), Poisson (λ = 2*Nx*), and normal (*μ* = 2*Nx*, *σ*^2^ = 2*Nx*(1 – *x*)) distributions, while for lower panel, we subtract the random draws (using same parameters) from 2*N*; where *N* = 500 and number of draws = 500000. The green vertical dotted line is at either 0 or 2*N* to indicate cases where the normal distribution yield non-feasible observations (allele copies below 0 and above 2*N*). From these plots, we consider less than 35 allele copies to be rare enough to use the Poisson approximation (as this threshold yields a good fit to the binomial and avoids negative samples or samples above 2*N* at extreme allele frequencies.)

### S3 Population trajectories

#### S3.1 Population Trajectories from DAA

**Figure S3.1:**
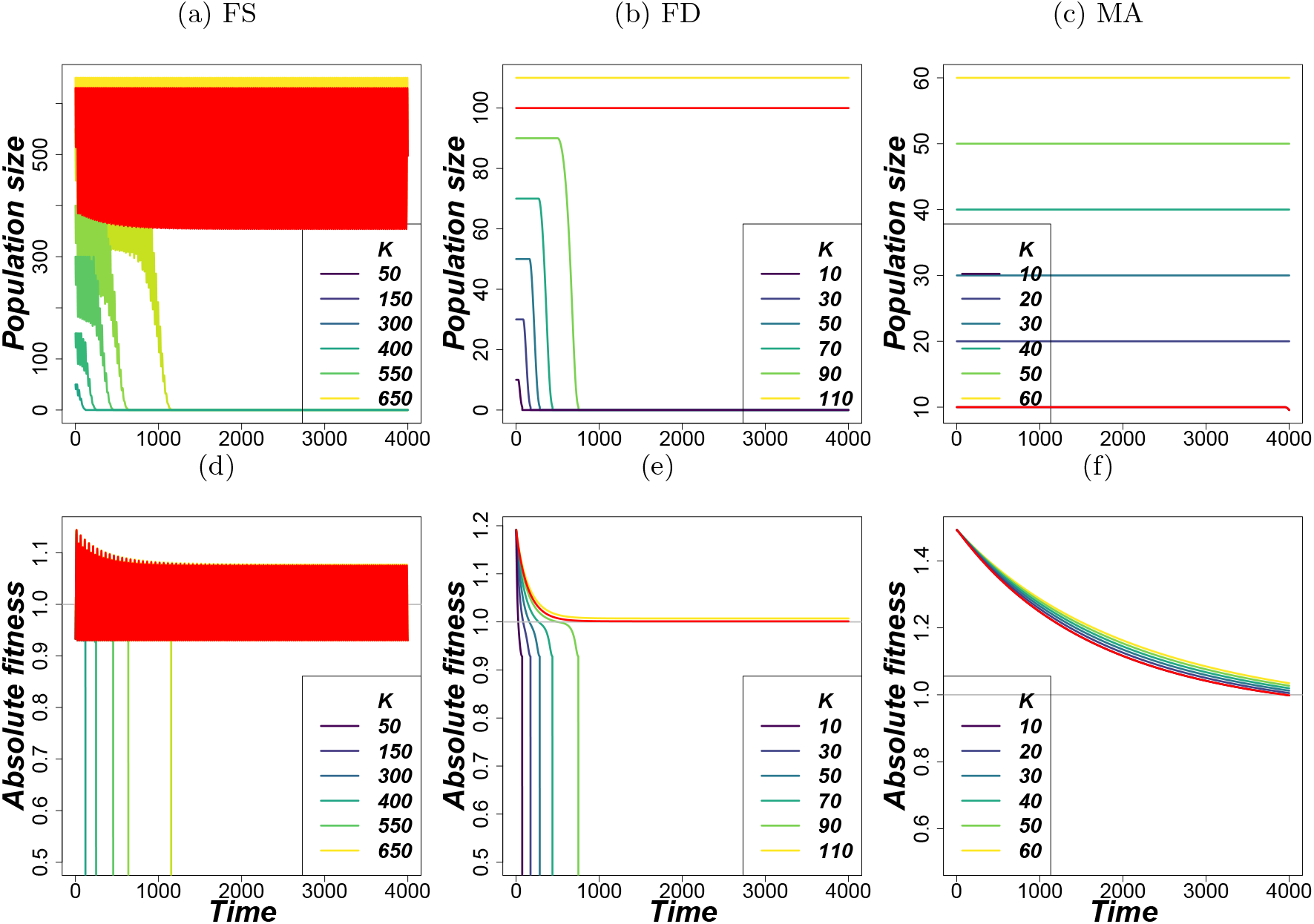
Isolated selection mechanisms: Trajectories for various carrying capacities from the diffusion approximation approach. Row 1: population size and Row 2: absolute fitness. The horizontal gray line in Row 2 corresponds to the threshold absolute fitness of 1. Parameters for balancing selection were *s_a_* = *s_A_* = 0.005, *μ_A_* = *μ_a_* = 0.0005, *κ* = 50 and for MA *s_A_* = 0, *s_a_* = 0.005, *h* = 0.5, *μ_A_* = 0.0005, 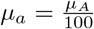. Other shared parameters were *n* = 100, *K* = *N*_0_, *r* = ln(1.53).

**Figure S3.2:**
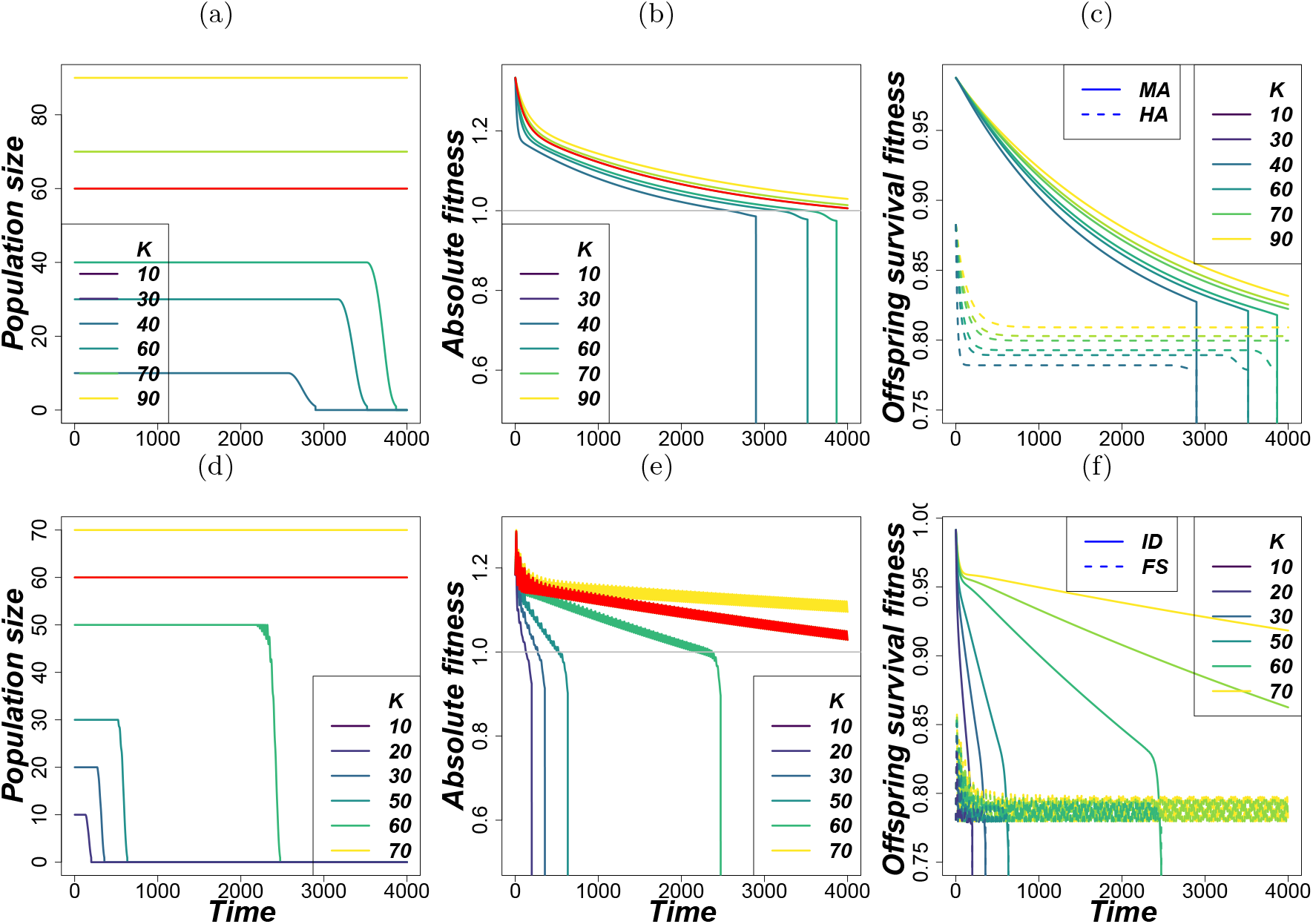
Interacting mechanisms: Trajectories for different carrying capacities from the diffusion approximation for Row 1: HA-MA and Row 2: FS-ID. Column 1: population size, Column 2: absolute fitness, Column 3: contribution of constituent selection mechanisms to offspring viability. The gray horizontal line in Column 2 is the threshold absolute fitness of 1. Under balancing selection: *s_a_* = *s_A_* = 0.005, *μ_A_* = *μ_a_* = 0.0005, *κ* = 50 and for ID and MA *s_A_* = 0, *μ_A_* = 0.0005, 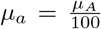 with *h* = 0.5, *s_a_* = 0.005 for MA and *h* = 0.01, *s_a_* = 0.05 for ID. Shared parameters: *n* = 100, *K* = *N*_0_, *r* = ln(1.53). The number of loci were equally shared between constituent mechanisms.

##### S3.1.1 Longer-time trajectories

**Figure S3.3:**
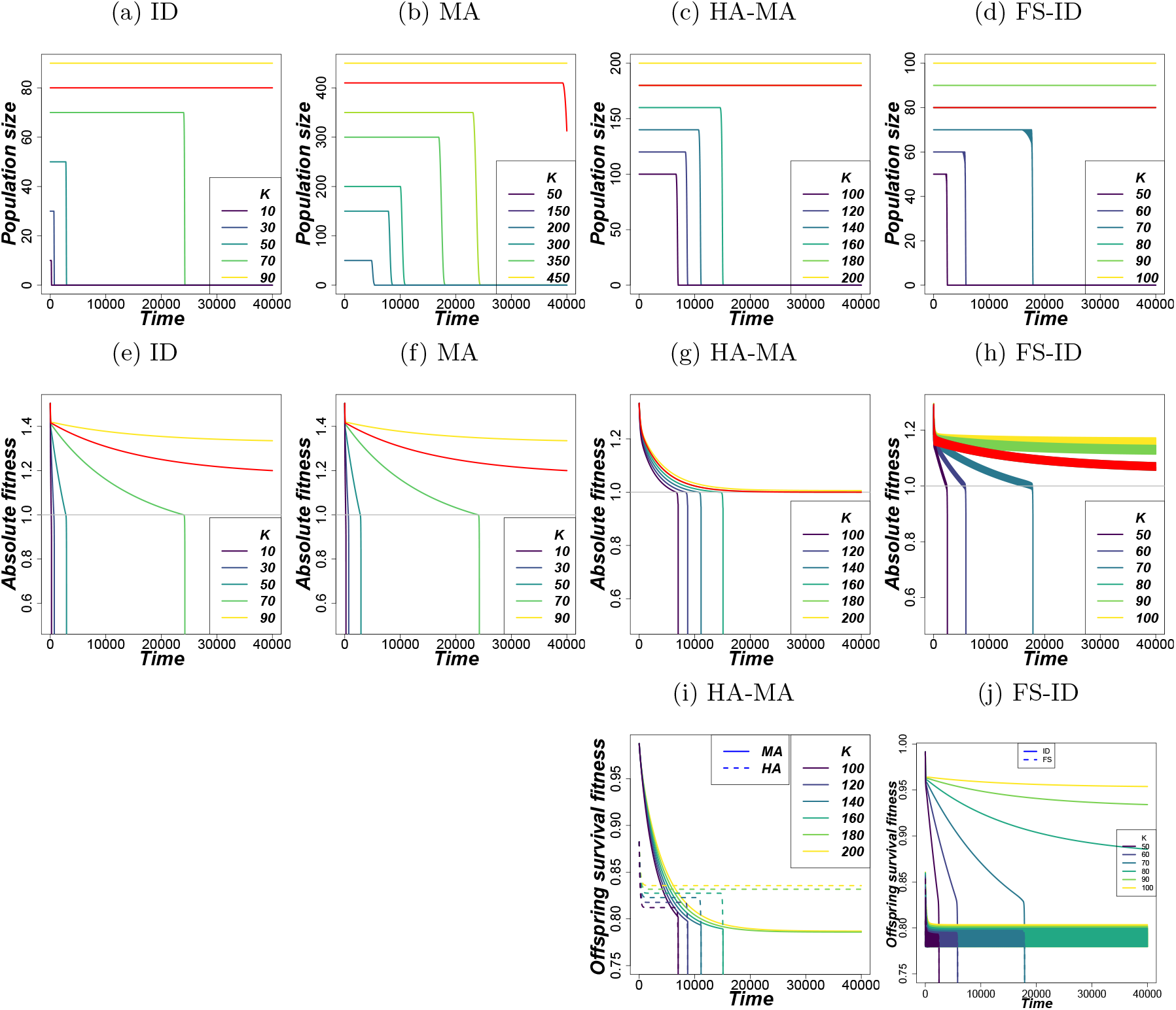
Longer-time population and fitness trajectories (run for 40000 generations) for various carrying capacities using the diffusion approximation approach. The gray horizontal line is the threshold absolute fitness of 1. The red line shows the MVP size estimate. In cases of more than one selection mechanism, the number of loci under each mechanism was equal. Parameters for balancing selection were *s_a_* = *s_A_* = 0.005, *μ_A_* = *μ_a_* = 0.0005, *κ* = 50 and for ID and MA *s_A_* = 0, *μ_A_* = 0.0005, 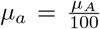 with *h* = 0.5, *s_a_* = 0.005 for MA and *h* = 0.01, *s_a_* = 0.05 for ID. Other shared parameters were *n* = 100, *K* = *N*_0_, *r* = ln(1.53).

#### S3.2 Population Trajectories from the IBM

**Figure S3.4:**
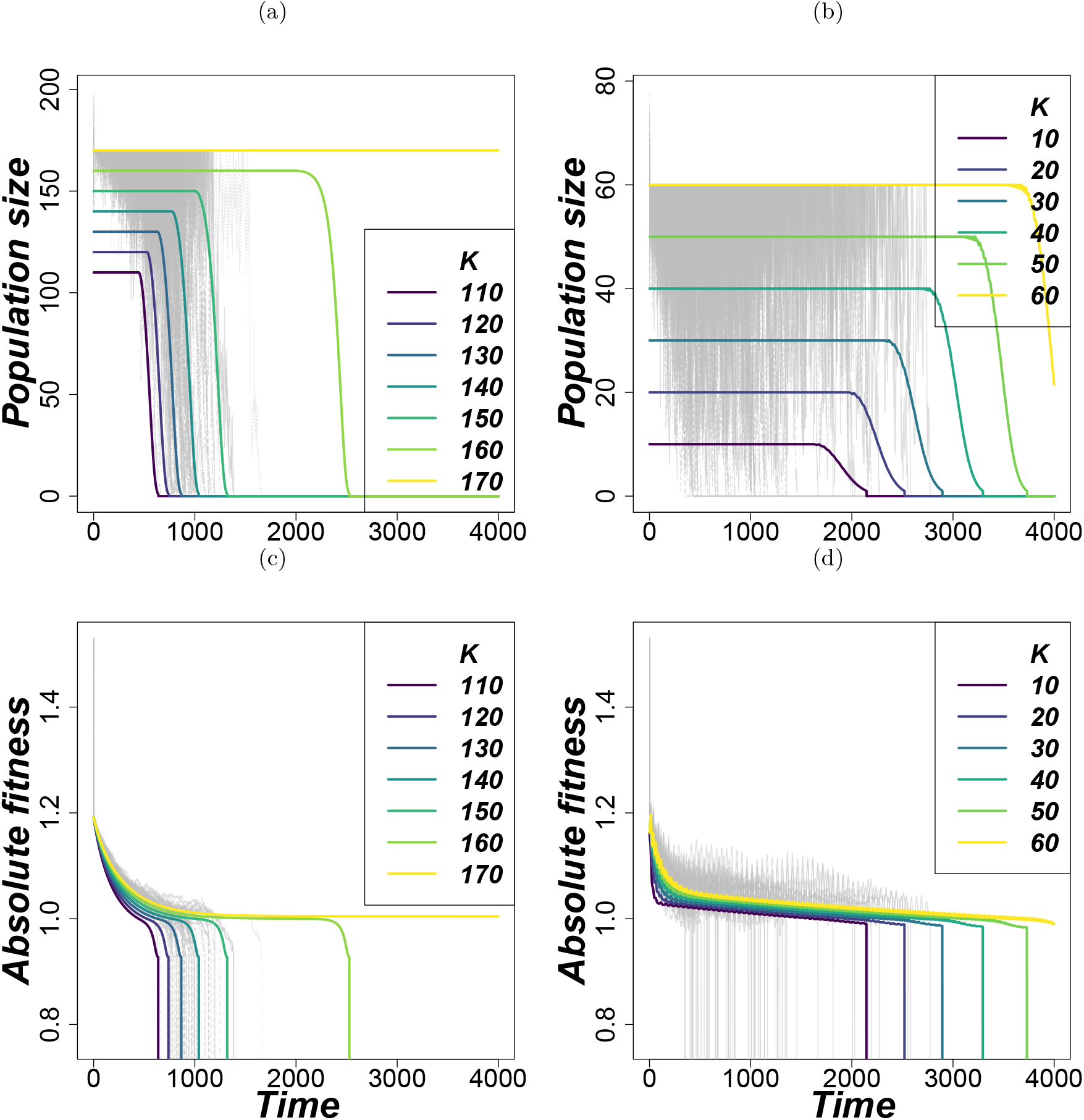
Individual-based simulations of 60 trajectories (in gray) for HA at *K* = 170 (column 1) and HA-FS-FD-ID-MA at *K* = 60 (column 2). The thick lines are for various carrying capacities using the diffusion approximation approach. Parameters for balancing selection are *s_a_* = *s_A_* = 0.005, *μ_A_* = *μ_a_* = 0.00022, *κ* = 50 and for ID and MA *s_A_* = 0, *s_a_* = 0.005, *μ_A_* = 0.00022, 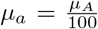 with *h* = 0.5 for MA and *h* = 0.0 for ID. Other shared parameters were *n* = 100, *K* = *N*_0_, *r* = ln(1.53). In column 2, the number of loci under each mechanism was equal (i.e., 20 loci each).

### S4 Critical Number of Loci and Equilibrium Fitness

#### S4.1 Equilibrium Fitness

We start with single-locus dynamics where the mean fitness is obtained from the deterministic equilibrium allele frequency *x* (which would be reached in a population of infinite size) as

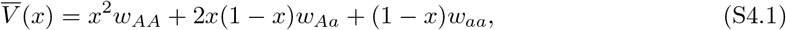

where *w_ij_*, *i,j* ∈ {*A, a*} is the equilibrium genotype fitness (geometric mean fitness in a complete cycle for FS) and 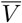 is the mean viability fitness for a given mechanism. Assuming equilibrium conditions and for symmetric selection, the equilibrium mean fitness for HA is given by

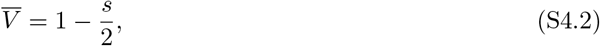

where *s* is the selection coefficient against homozygotes.

For a purely recessive deleterious allele, the fitness is given by (assuming Hardy-Weinberg proportions in an infinite population; Felsenstein (2015))

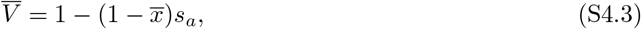

where *s_a_* the selection coefficients against the homozygote of the recessive allele *a* and 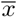 is the frequency of wild-type allele *A* approximated by

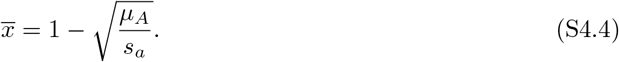

However, when the deleterious allele is not completely recessive,

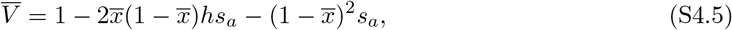

where *h* > 0 is the dominance coefficient and this time 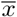 is approximated by

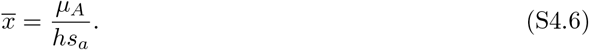

Note that when *s_a_* < *μ_A_* in Equation S4.4 and *hs_a_* < *μ_A_* in Equation S4.6 gives 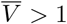 which does not make sense. So, the approximations fail for these cases because the equilibrium allele frequency of the wild-type allele will be below 0. Ideally, in such cases, selection against deleterious allele is so weak and yet their inflow via mutation is so high and thus it will result into fixation of the deleterious allele.

In Fig. S4.1, we show effect of increasing selection coefficient and mutation rates on MVP estimates for loci under mutation-selection-drift equilibrium. At a given mutation rate, as *s_a_* (or *hs_a_*) increases, the MVP decreases rapidly. Also, for a given selection coefficient, increase in mutation increases MVP size estimates. These observations are expected as shown from Equations (S4.4) and (S4.6). The increase in *s_a_* reduces the frequency of the deleterious allele. Thus strong purging of deleterious alleles eventually lead to a higher equilibrium fitness. On the other hand, increase in *μ_A_* increases frequency of the deleterious mutation at equilibrium leading to low equilibrium fitness.

**Figure S4.1:**
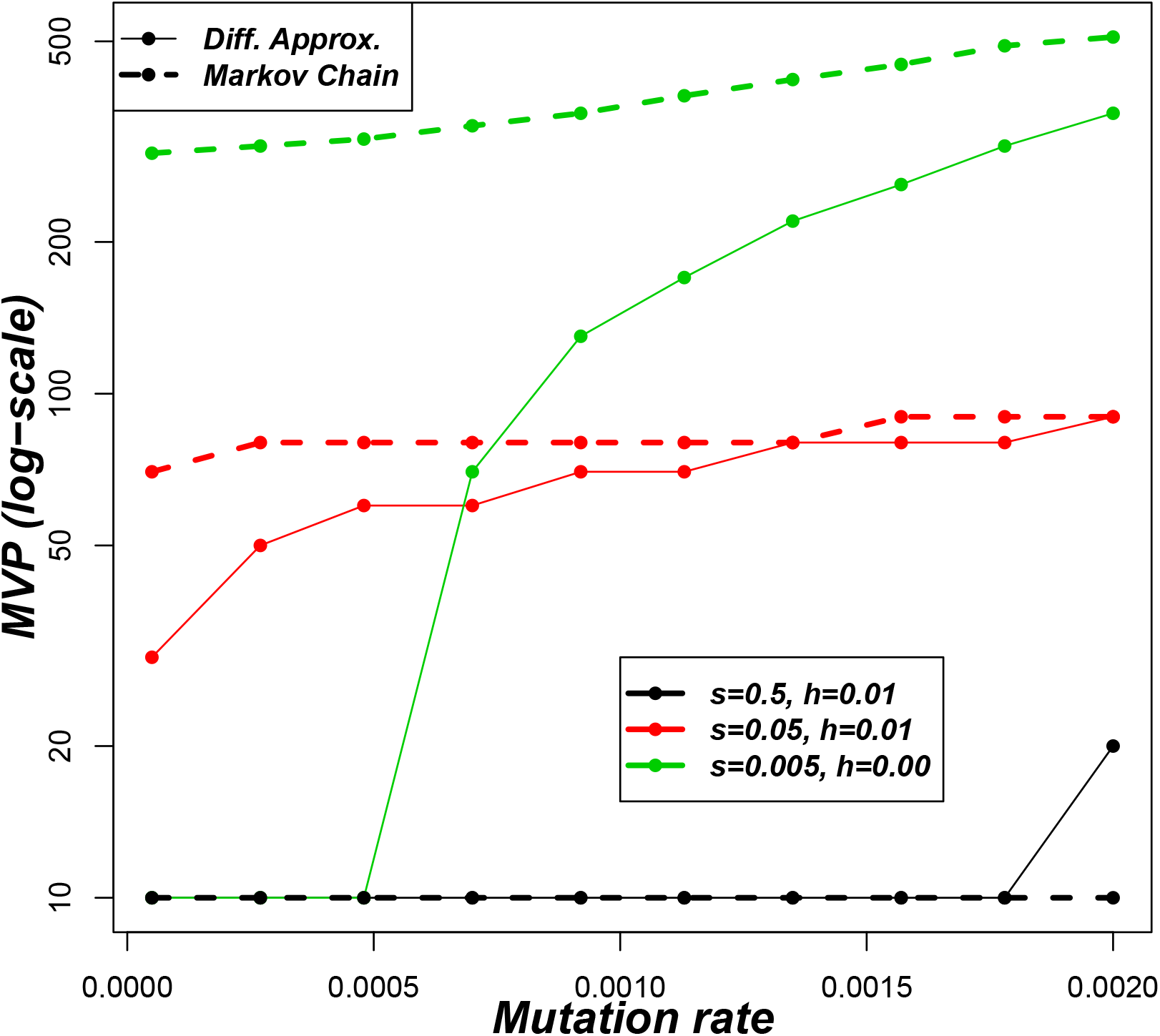
MVP estimates for populations under different selection pressure from inbreeding depression with fixed number of loci *n* = 100. On the x-axis, we have the forward mutation rate *μ_A_*. Other parameters are 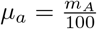, *K* = *N*_0_, *r* = ln(1.53).

#### S4.2 Critical number of loci

It is possible to have the critical number of loci *n_c_*, above which no arbitrary size of a population can survive. Here, *n_c_* is obtained based on the assumption that absolute fitness of 1 is the threshold below which populations go extinct i.e., 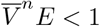, *n* is the number of loci under selection,

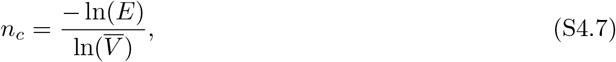

while for two mechanisms, 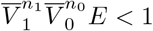, *n_i_* being number of loci under respective selection mechanism.

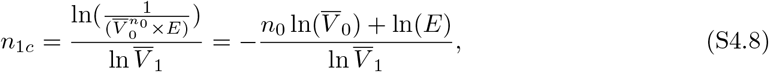

where *n*_0_ is the number of loci originally occupied by a genetic mechanism whose equilibrium fitness per locus is *V*_0_ and *n*_1*c*_ is the critical number of loci added from the new genetic mechanism whose mean fitness per locus is *V*_1_, both evaluated at the respective equilibrium frequencies 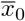 and 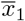 while *E* is the expected number of offspring per individual. So, *n*_1*c*_ gives the maximum additional loci that can be acted on to have a finite population size surviving. We illustrate how *n_c_* varies with selection coefficient, fitness and mutation rate in Figure S4.2

**Figure S4.2:**
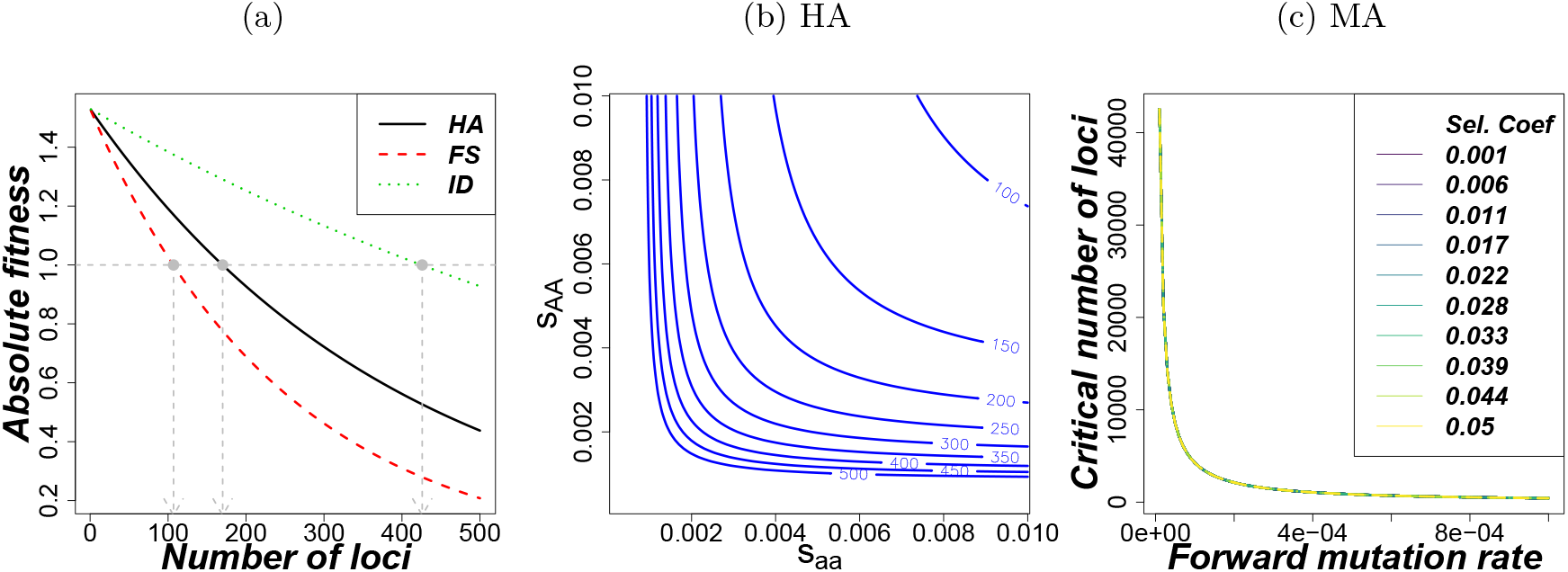
(a) How absolute fitness changes with increasing number of loci under selection (assuming infinite population size). Parameters used *s_a_* = *s_A_* = 0.005 for HA and FS, *s_a_* = 0.05, *s_A_* = 0, *μ* = 0.001, *h* = 0 for ID (b) Shows how critical number of loci (that results into an asymptotic increase in MVPs) varies with selection coefficients in HA (from Equation (S4.7)). (c) Shows how the critical number of loci under selection varies with mutation rate *μ_A_* for different selection coefficients for MA. Parameters used 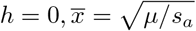 for the deleterious allele.

#### S4.3 Effect of Number of Loci on MVP estimates from the MCA

**Figure S4.3:**
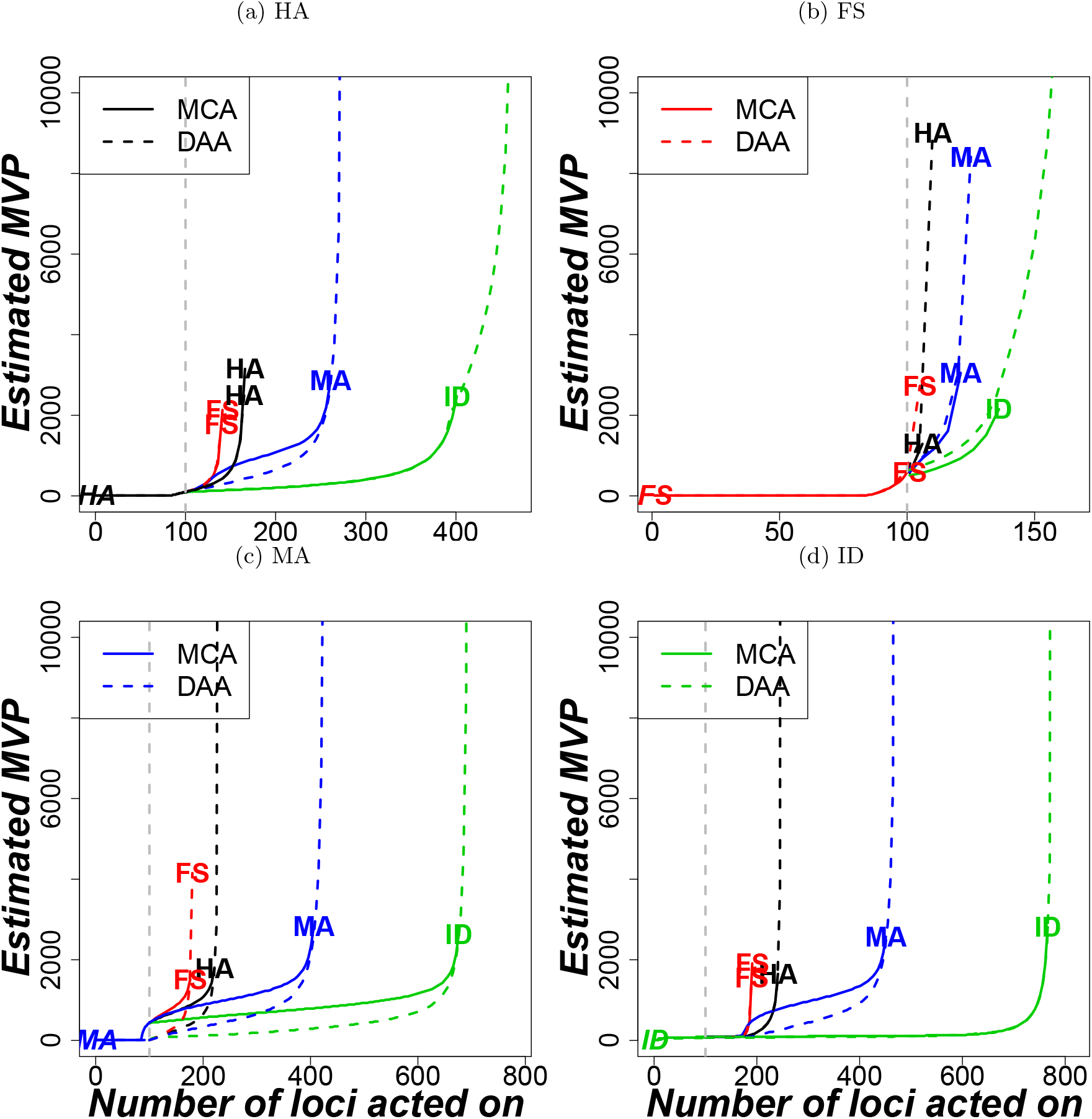
Effect of increasing number of loci under selection on MVP size. Same data as in 4 but the y-axis not log-scaled but limited to a maximum of 10000. The genome from populations with at most 100 loci (below the vertical gray line) was acted on by the same selection mechanism (indicated at the start of the line) (a) HA, (b) FS, (c) ID and (d) MA. Those genomes with above 100 loci, the mechanism indicated at the end of the line acted on the remaining loci, e.g., the line with HA—ID label implies the first 100 loci were acted on by HA while ID acted on the remaining loci. Parameters for balancing selection were *s_a_* = *s_A_* = 0.005, *μ_A_* = *μ_a_* = 0.0005, for ID and MA, *s_A_* = 0, 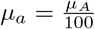 while *h* = 0.01, *s_a_* = 0.05 for ID and *h* = 0.5, *s_a_* = 0.005 for MA. Shared parameters were *K* = *N*_0_, *r* = ln(1.53).

**Figure S4.4:**
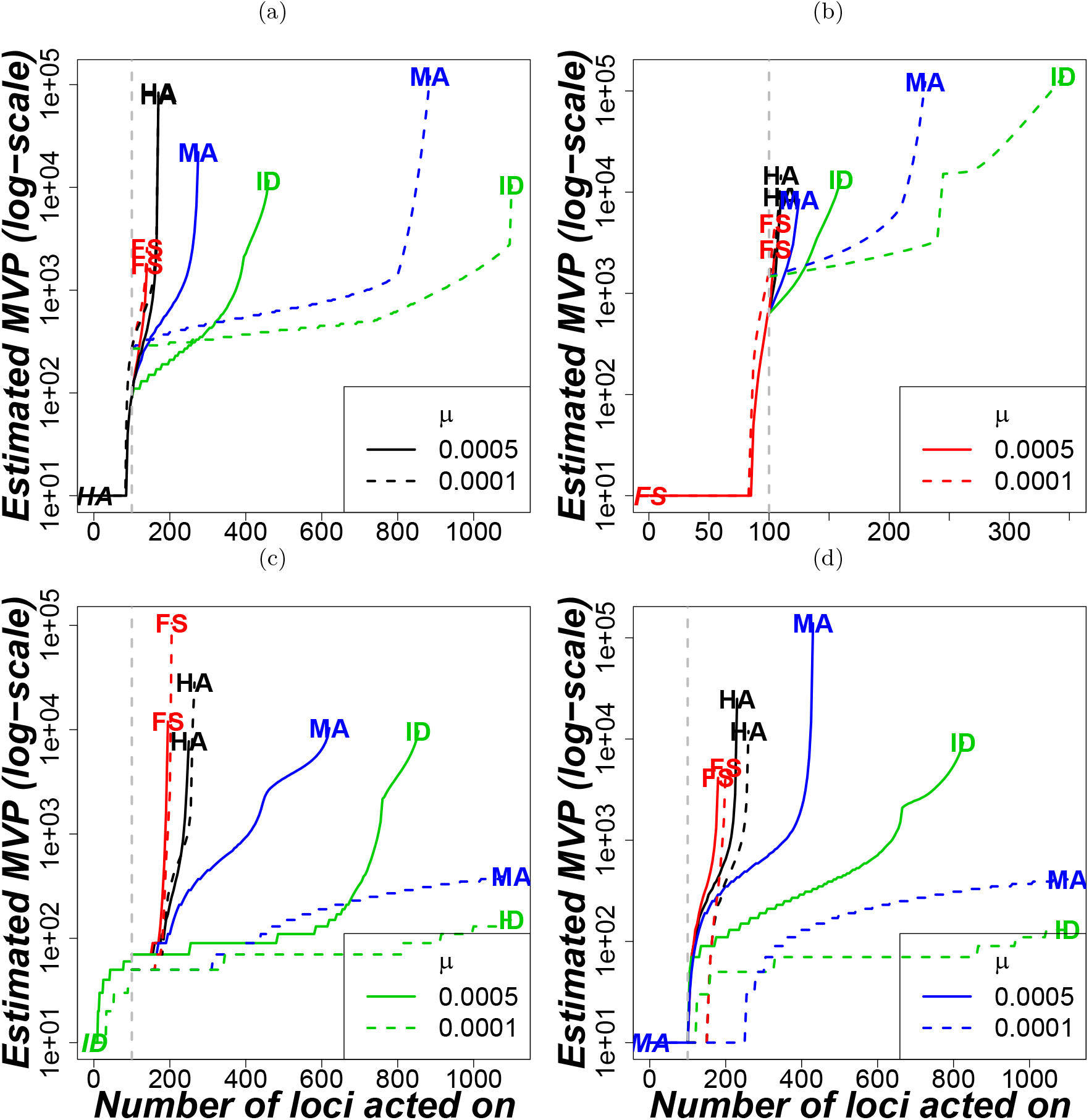
Effect of mutation rates on MVP sizes and critical number of polymorphic loci above which no arbitrary finite population survives. The results used the diffusion approximation approach (DAA). Mutation rates only affects non-balancing selection mechanisms. Genomes with less than 100 loci (separated by the vertical gray line) are acted on by the same selection mechanism (a) HA, (b) FS, (c) ID and (d) MA. Those with above 100 loci, the mechanism indicated at the end of the line acts on the remaining loci, e.g., the line with HA—ID label implies the first 100 loci were acted on by HA while ID acts on the possible available loci. Here, *μ* = 0. 0001 for unbroken lines and *μ* = 0.0005 for broken lines. Other parameters for balancing selection were *s_a_* = *s_A_* = 0.005, *μ_A_* = *μ_a_*, *κ* = 50, for ID and MA, *s_A_* = 0, 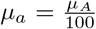 while *h* = 0.01, *s_a_* = 0.05 for ID and *h* = 0.5, *s_a_* = 0.005 for MA. Shared parameters were *K* = *N*_0_, *r* = ln(1.53).

**Figure S4.5:**
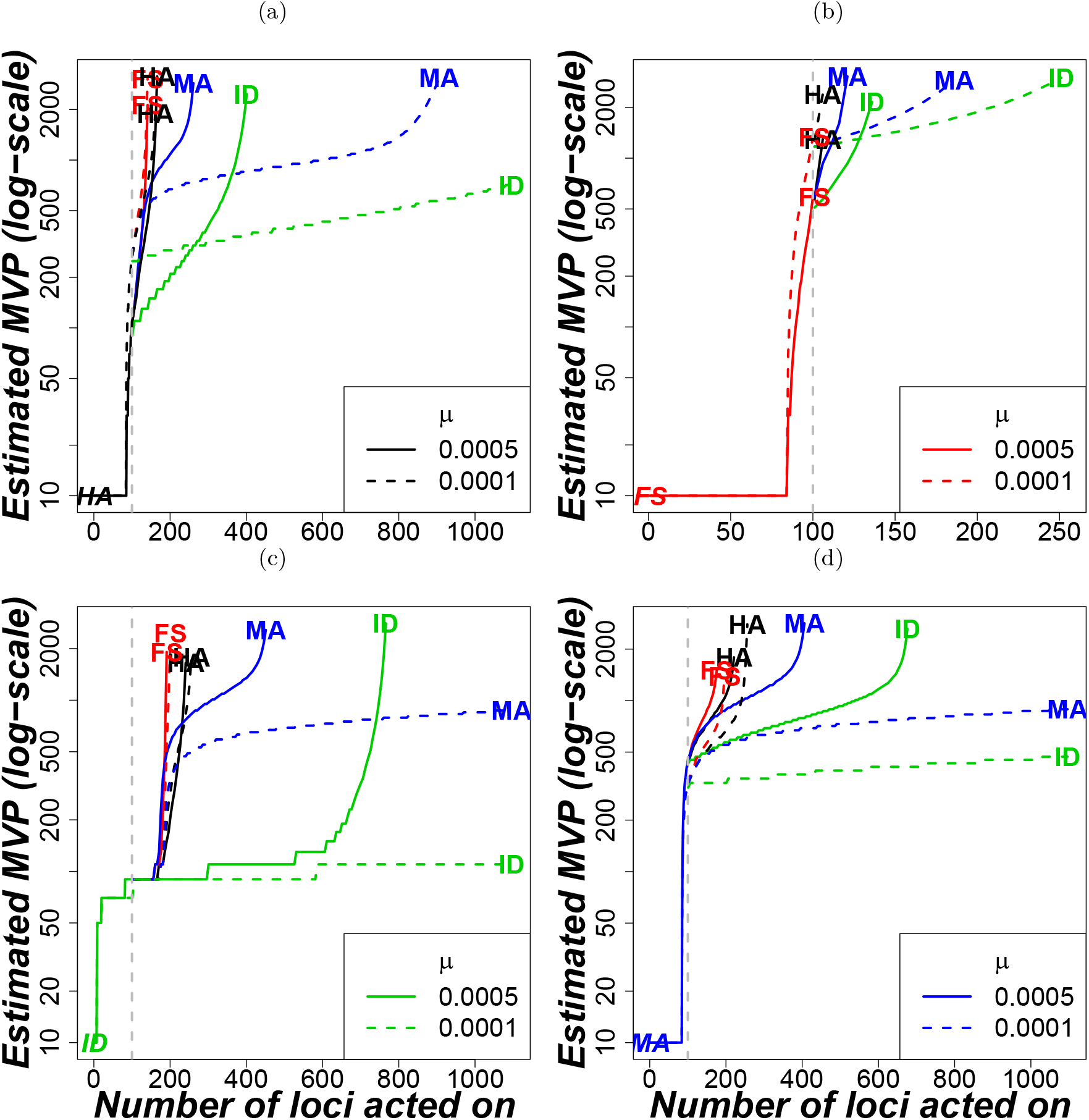
Effect of increasing number of loci under selection on MVP size. Everything is as in Fig. S4.4 but this is shown for the Markov chain mechanism.

### S5 MVP Estimates from Fixed Number of Loci

#### S5.1 MVP Estimates from IBM with Fixed Number of Loci

**Figure S5.1:**
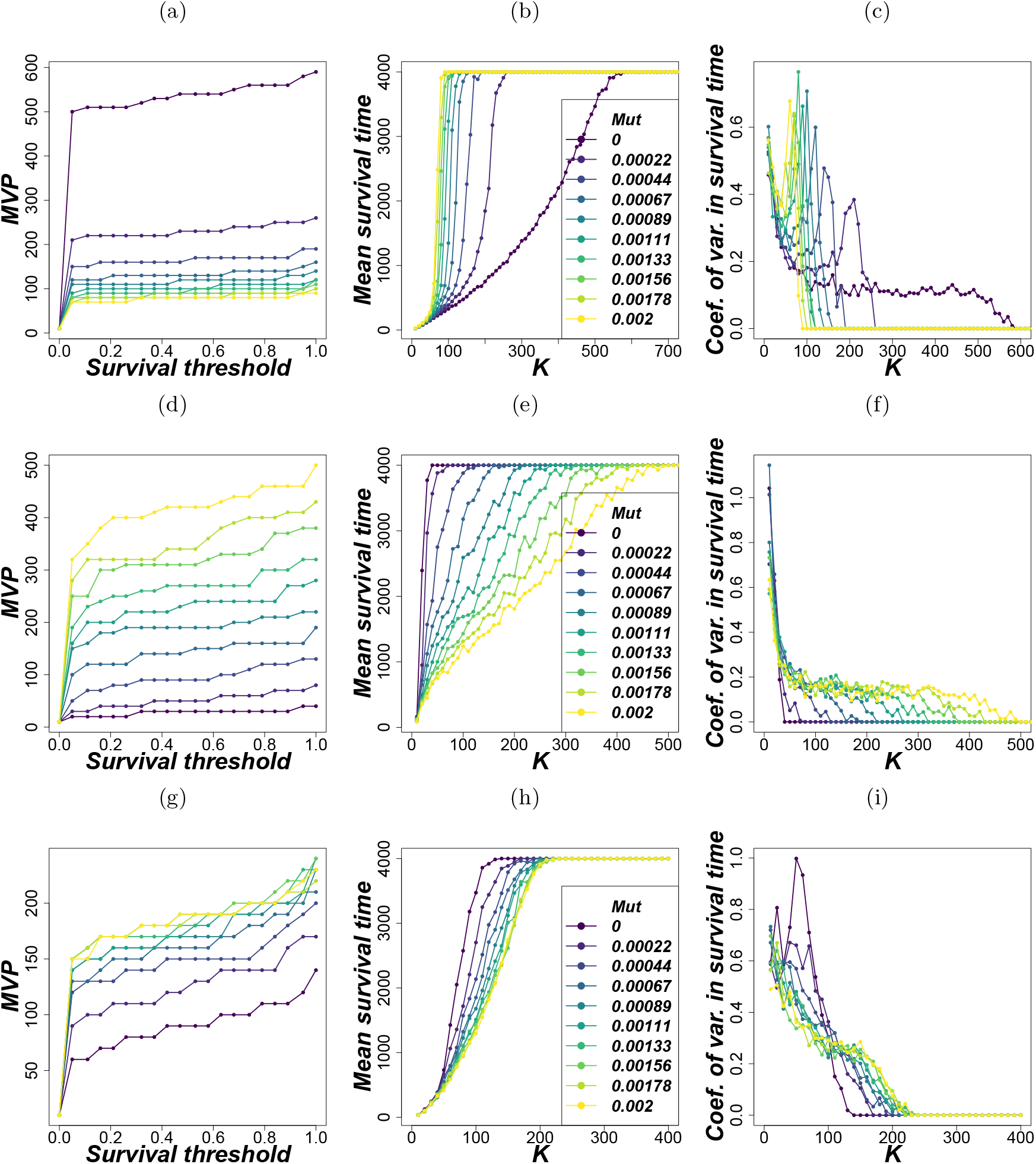
MVP estimates from the IBM. Row 1: HA only, Row 2: ID only, and Row 3: HA-FS-FD-ID-MA. 1st column: MVP estimates for different mutation rates measured at different levels of survival probability. 2nd column: Mean survival time of populations at different carrying capacities estimated under different mutation rates. 3rd column: Coefficient of variation in survival times of populations at different carrying capacities estimated under different mutation rates. Parameters for balancing selection; *s_a_* = *s_A_* = 0.005, *μ_A_* = *μ_a_*, *κ* = 50, for ID and MA *s_A_* = 0, *s_a_* = 0.005, 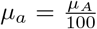 while *h* = 0.5 for MA and *h* = 0 for ID. Other shared parameters were *K* = *N*_0_, *r* = ln(1.53) and a total of 100 loci were equally shared by the constituent mechanisms.

Figure S5.1 shows MVP size estimates from the IBM. The MVP size estimates from the IBM were qualitatively similar to those estimated by DAA and MCA (Fig. 5 in the main text). Quantitatively, estimates from the IBM were (slightly) higher than corresponding estimates (Fig. 5 row 2). The MVP estimates were fairly insensitive to levels of survival probability threshold (Fig. S5.1 column 1). As expected, the mean time to extinction increased rapidly with increasing carrying capacity *K* (Fig. S5.1 column 2) Also, there was more uncertainty in the extinction times in populations with lower carrying capacities than their counterparts with large capacities (Fig. S5.1 column 3).

#### S5.2 MVP Estimates with Fixed Number of Loci Including those from Negative Frequency-Dependent Selection (FD)

**Figure S5.2:**
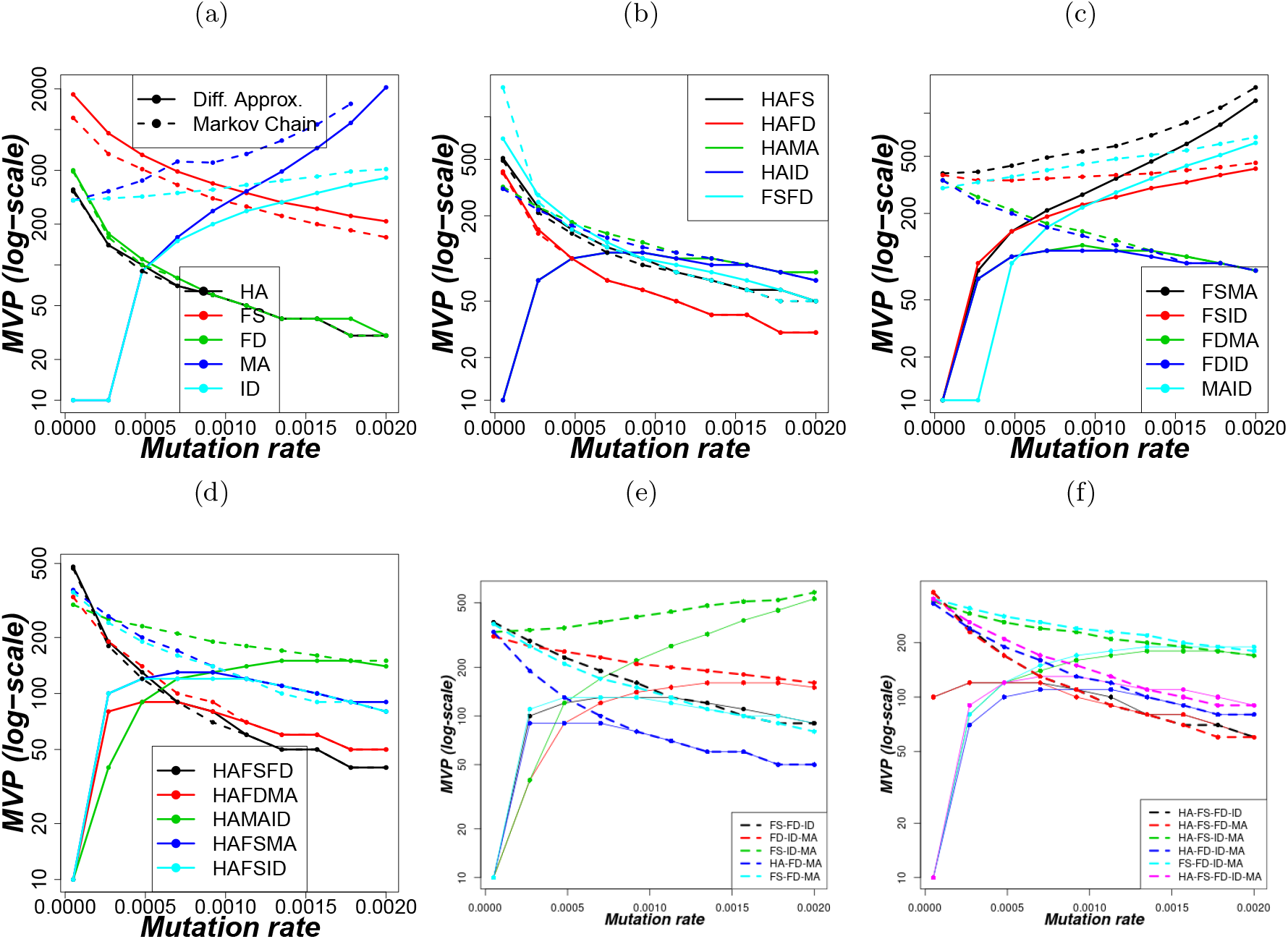
MVP estimates from the Markov chain approach (with broken thick lines) and the diffusion approximation (with unbroken thin lines). Parameters *s_a_* = *s_A_* = 0.005, *K* = *N*_0_, *r* = ln(1.53), *μ_A_* = *μ_a_*, *κ* = 50 except for ID and MA where *s_A_* = 0, 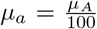 and also for ID, *h* = 0.5. The total number of loci on the genome is *n* = 100 except for genomes with 3 selection mechanisms where *n* = 99 for purposes of equal distribution of loci.

#### S5.3 Arithmetic, Geometric and Harmonic Mean of the MVPs from Constituent Selection Mechanisms

**Figure S5.3:**
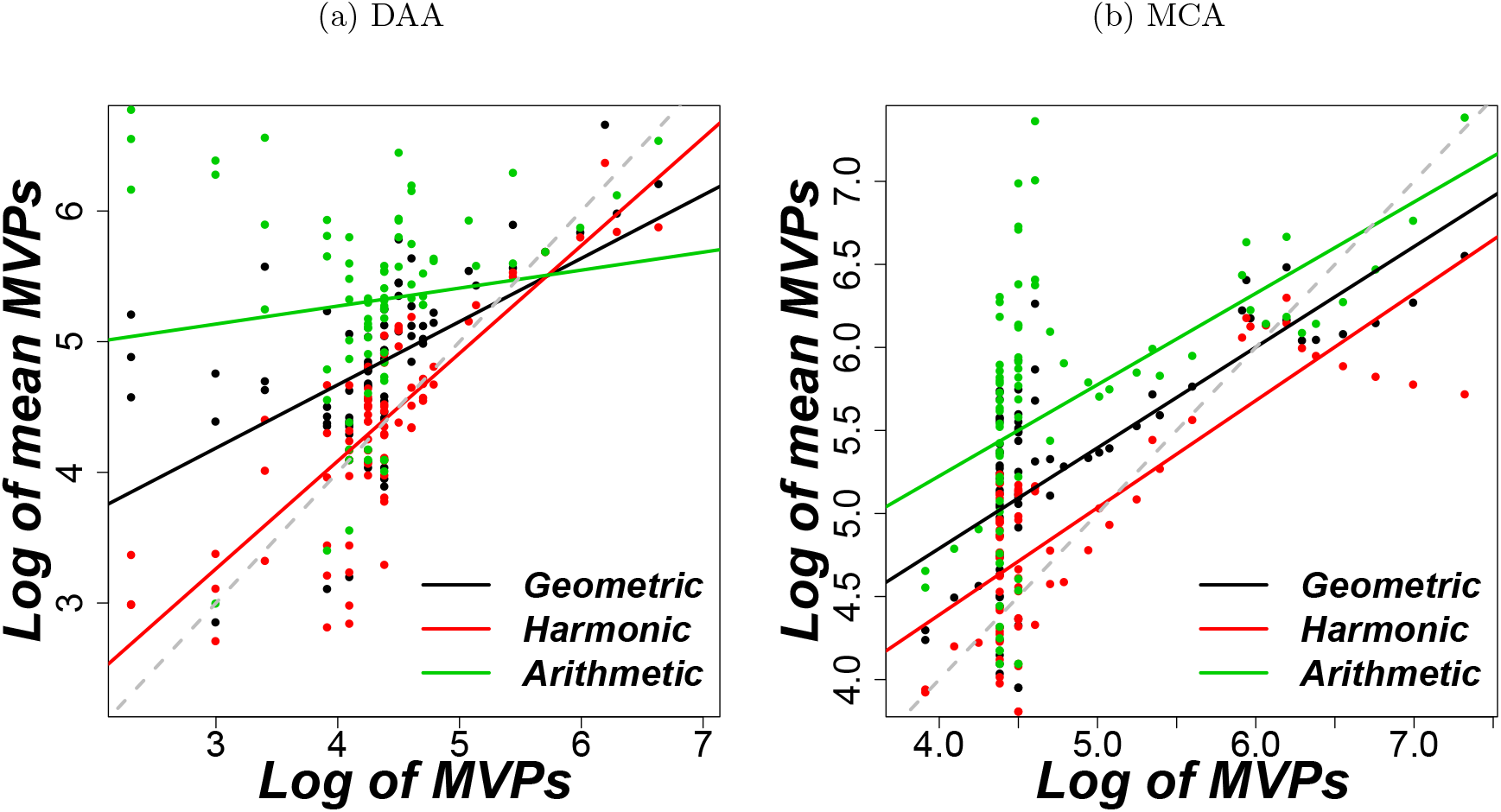
Comparison of the estimated MVPs (on x axis) to the geometric, harmonic and arithmetic means (on y axis) of the constituent isolated selection mechanisms for a given scenario using (a) the diffusion approximation and (b) the Markov chain approaches. The different colours show different calculated means (black, red, and green points for geometric, harmonic, and arithmetic means respectively) while the solid lines are the regression lines for each calculated mean. Parameters were as in Fig. S5.2. The points are from HAFS, HAID, FSMA, MAID, HAFSMA, HAFSID, FSMAID, and HAFSMAID.

In Fig. S5.3, we compare the joint MVP to the different means of the MVPs estimated from the isolated selection mechanisms. Generally, the observed MVPs under a mixture of mechanisms are closer to the harmonic and geometric means than the arithmetic mean of the MVPs calculated for the mechanisms in isolation with the same total number of loci. To a large extent, the joint MVPs are (slightly) above the calculated harmonic means and (slightly) below the geometric means. We briefly illustrate why joint MVP is closer to the harmonic mean in Section S6.

### S6 Relationship between MVP estimates and fitness across loci

**Figure S6.1:**
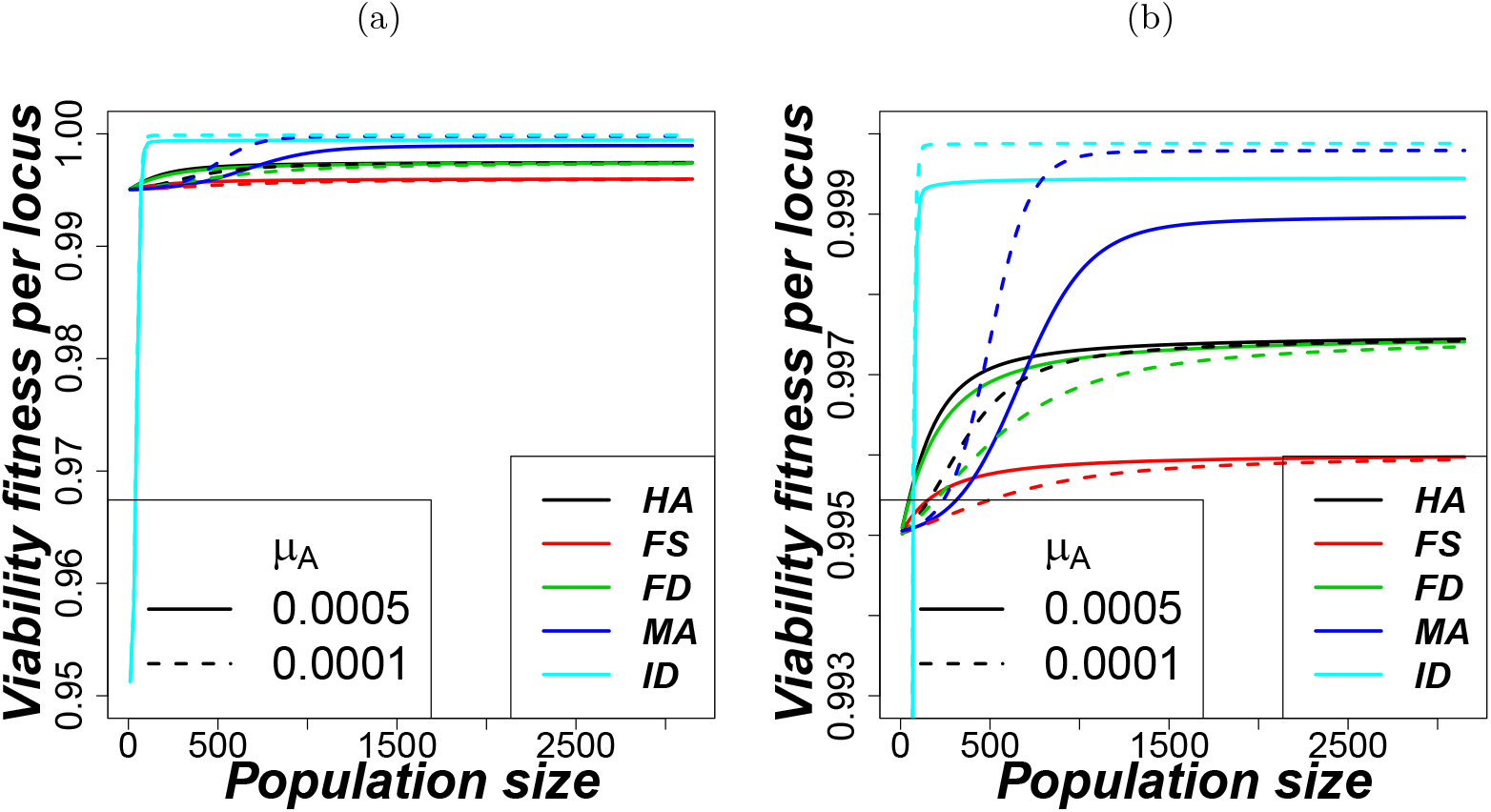
Viability fitness per locus at different population sizes for the different mechanisms from the Markov chain approach at equilibrium (Equation (S2.6)). Parameters: *s_a_* = *s_A_* = 0.005, *μ_A_* = *μ_a_*, *κ* = 50 for balancing selection mechanisms and for MA: *s_A_* = 0, *s_a_* = 0.005, *h* = 0.5, 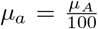, and ID: *s_A_* = 0, *s_a_* = 0.05, *h* = 0.01, 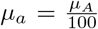. Panel (b) zooms in on the upper viability range in (a).

**Figure S6.2:**
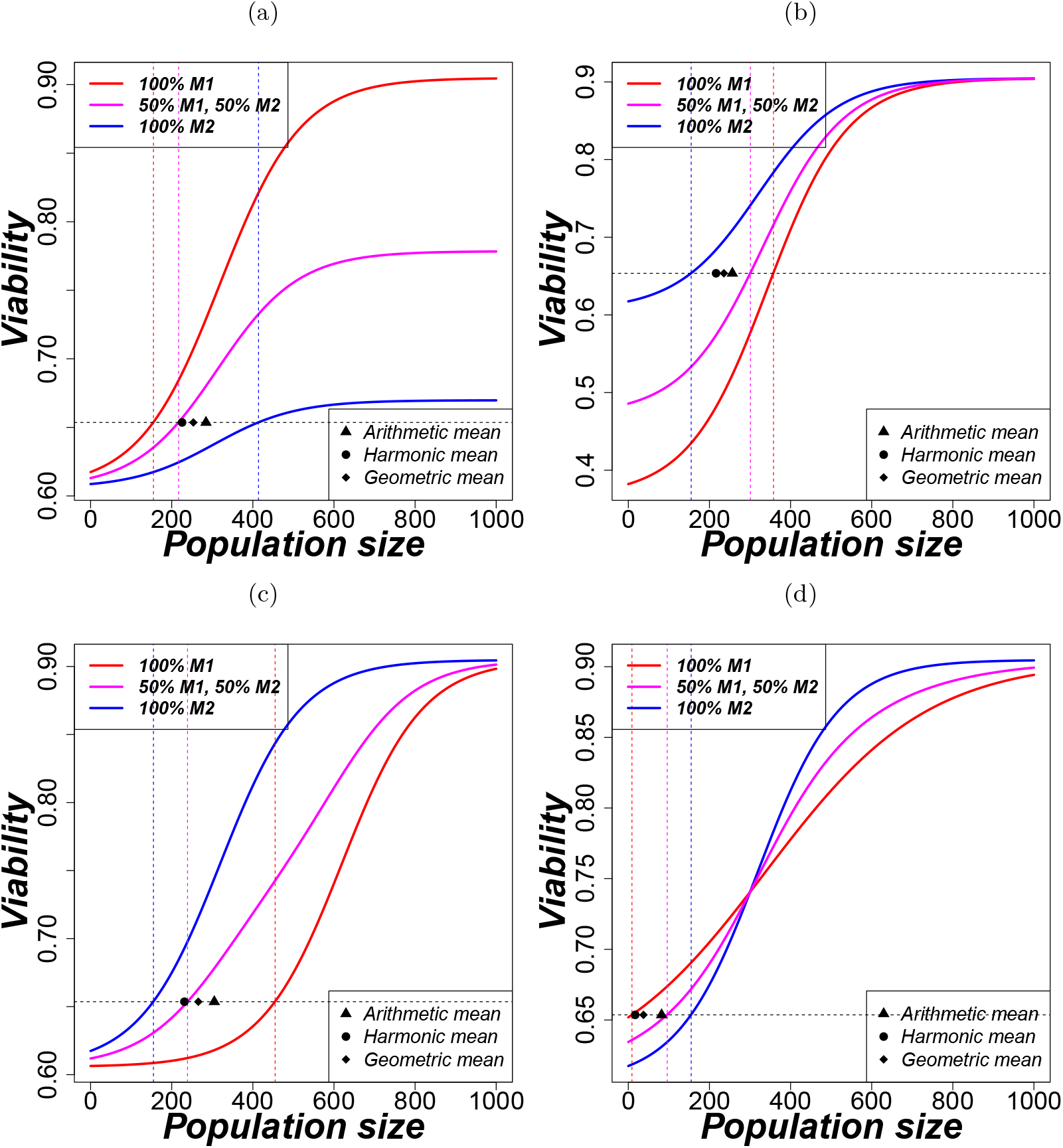
Logistic curve fitting 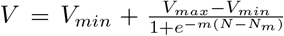, where *V* is the viability per locus, *V_min_* and *V_max_* are the minimum and maximum viability per locus, and *N_m_* is the population size *N* of the viability sigmoid midpoint, and *m* is steepness of the curve. Illustration of the relationship between minimum viable population sizes in a logistic curve setting (as observed from Figure S6.1). To get viability for a single mechanism with *n* = 100 loci, we simple use *V^n^* (blue and red solid lines), and for two mechanisms, we use 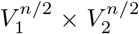 (magenta solid line). The corresponding minimum viable population sizes with a critical viability *v_crit_* = 1/*E* = 1/1.53 are shown as dashed vertical lines. The circle, diamond, and triangle indicate the harmonic, geometric, and arithmetic mean of the two separate minimum viable population sizes, respectively.

**Figure S6.3:**
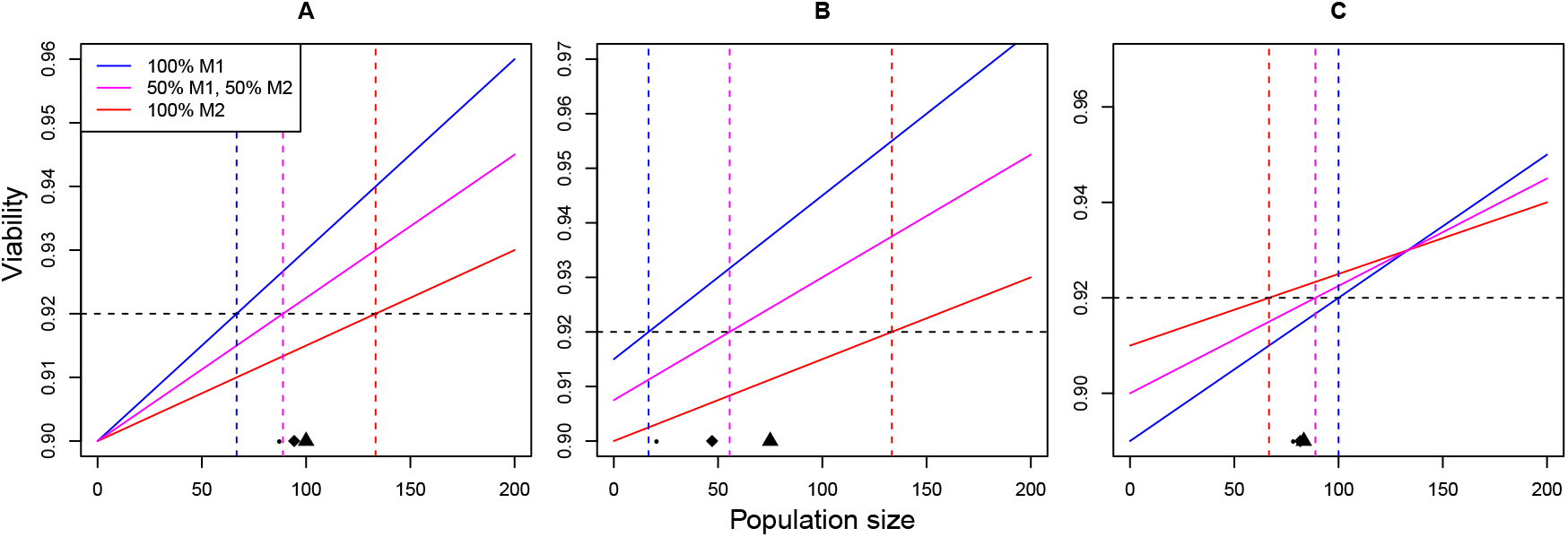
Illustration of the relationship between minimum viable population sizes in a simple linear setting. Assuming approximately additive effects of small fitness reductions, the viability with 50% of loci under mechanism 1 and 50% under mechanism 2 for a given population size (magenta solid line) is the arithmetic mean of viabilities for the two separate mechanisms acting at 100% of loci (blue and red solid lines). The corresponding minimum viable population sizes with a critical viability *v_crit_* = 1/*E* = 0.92 are shown as dashed vertical lines. The circle, diamond, and triangle indicate the harmonic, geometric, and arithmetic mean of the two separate minimum viable population sizes, respectively.

Here we use a simple setting to illustrate possible relationships between MVP sizes under single versus multiple genetic mechanisms. For this section, we assume that the relationship between population size and viability at low population sizes can be well approximated by a linear relationship (Figures S6.1 and S6.2). Assume that we have two genetic mechanisms (*i* = 1, 2) that when acting at *n* loci have viability

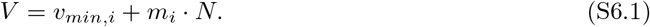

The respective minimum viable population size then is the population size at which viability takes the critical value *v_crit_* = 1/*E* at which absolute fitness is 1. Thus,

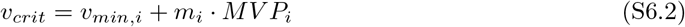

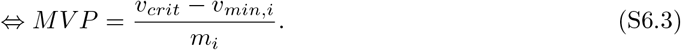

As long as deviations of viability from 1 are small, loci make roughly additive contributions to reducing viability and thus the viability relationship with half of the loci under mechanisms 1 and half of the loci under mechanism 2 is

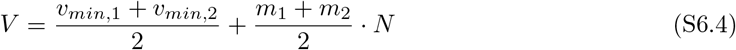

and the corresponding minimum viable population size is

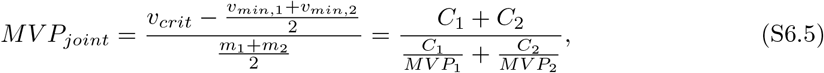

where *C_i_* – *v_crit_* – *v_min,i_*, *i* = 1, 2.

We now consider three different exemplary cases (Fig. S6.3). In the first case (Fig. S6.3 A), the two mechanisms have the same minimum viability *v*_*min*,1_ = *v*_*min*,2_ = *v_min_* (i.e., *C*_1_ = *C*_2_) as was also approximately the case in Fig. S6.1. Then

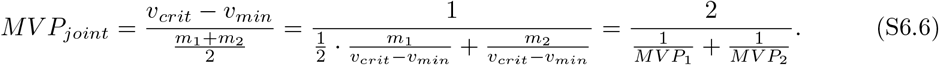

That is, the joint minimum viable population size (magenta vertical line in Fig. S6.3 A) is equal to the harmonic mean (indicated by a black dot in Fig. S6.3 A) of MVP sizes under mechanism 1 and 2. Intuitively, this is the case because minimum viable population size is inversely proportional to the rate at which viability increases with increasing population size. This is approximately true even when the viability functions are nonlinear and thus explains why we often obtain joint MVP sizes close to the harmonic mean of individual MVP sizes.

However, this equality does not hold if the two mechanisms have different *v_min_* (i.e., *C*_1_ ≠ *C*_2_). Depending on the relation of the two slopes and intercepts, we can then get scenarios where the joint MVP is larger than the harmonic mean of MVPs but still smaller than the arithmetic mean (Fig. S6.3 B) or scenarios where the joint MVP is even larger than the arithmetic mean of MVPs (Fig. S6.3 C). Also for general nonlinear viability functions, all cases are possible and the exact relationship between MVPs will depend on the precise shape of the curves.

### S7 Initial allele frequency

Figure S7.1 shows how initial allele frequency depends affects MVP estimates at different mutation rates for different isolated mechanisms. Initial allele frequency has large effects in FS. For loci under mutation-selection-drift balance (ID and MA), MVPs are insensitive to initial allele frequency unless the frequency of the deleterious allele is extreme like above 90%. Similarly, for HA, only very extreme allele frequencies have an effect on MVPs.

**Figure S7.1:**
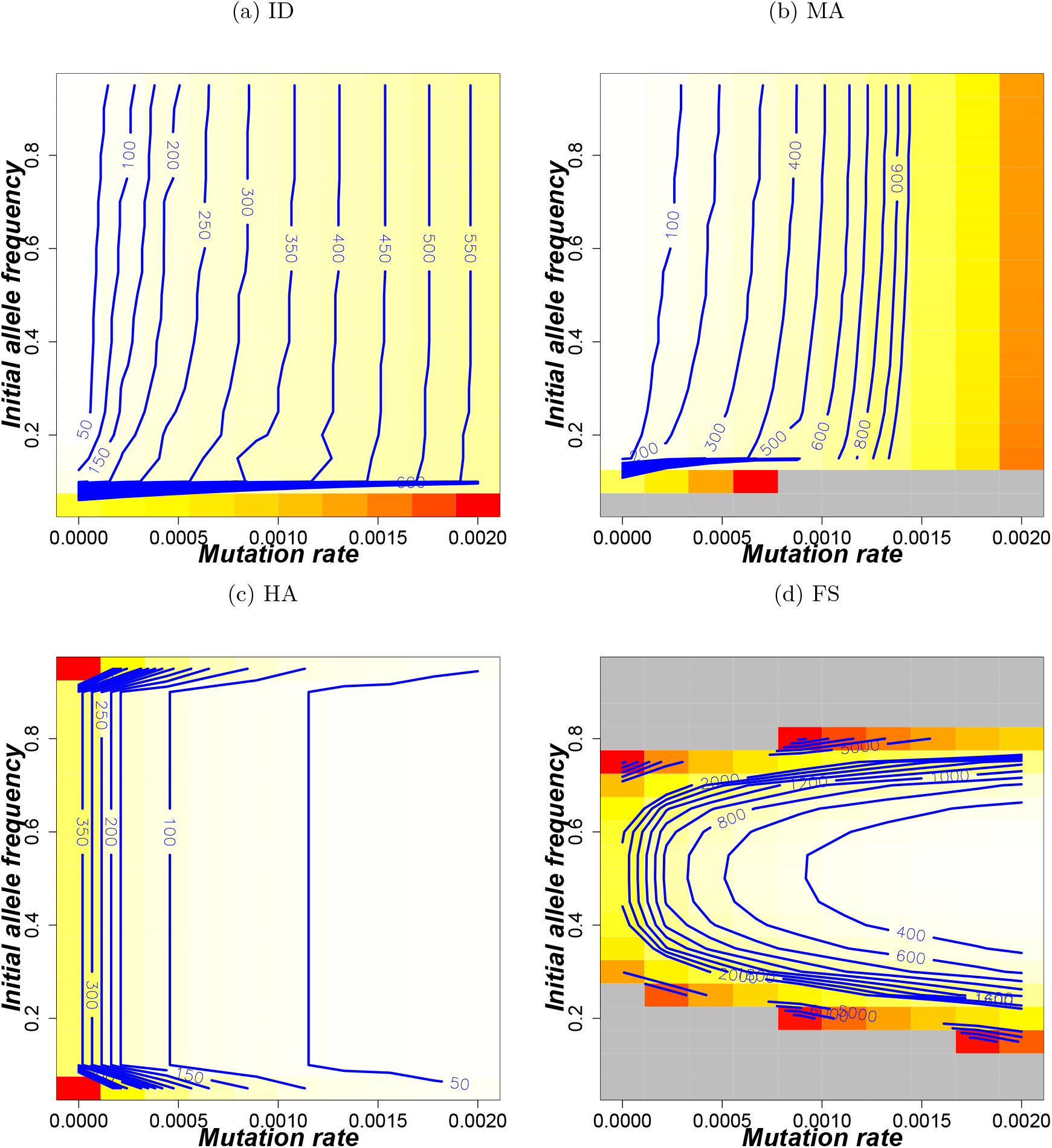
Effect of initial allele frequency on MVP size estimates. The blue lines show MVP trajectories at given initial allele frequency. The parameters for balancing selection were *s_a_* = *s_A_* = 0.005, *μ_A_* = *μ_a_*, *κ* = 50, for ID and MA *s_A_* = 0, *s_a_* = 0.005, 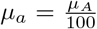 while *h* = 0.5 and *h* = 0 were respectively for MA and ID. Other shared parameters were *K* = *N*_0_, *r* = ln(1.53). The gray shaded parts have their MVP estimates above 10000.

