## Supplementary material for "Modeling minimum viable population size with multiple genetic problems of small populations": R codes used to simulate results: Execute_DiffusionApproximation_FixedLoci.R

```
rm(list=ls())

# Packages mainly for parallelisation
library("doMC")
cores=detectCores()
registerDoMC(cores)
require(reshape2)

setwd("~/MVP_GENETICPROBLEMS")
source("MainFunct_DiffusionApproximation.R")

# List of selection mechanism combintation. Each item is a vector of length 5. Within the vector, the 1st, 2nd, 3rd, 4th and 5th elements are the proportion 
# of loci under HA, FS, FD, ID and MA. e.g c(1,0,0,0,0) implies 100% HA loci, c(1/5,1/5,1/5,1/5,1/5) implies 1/5th loci under each mechanism
# NOTE THAT RUNNING THROUGH THE LIST BELOW CAN TAKE SEVERAL HOURS/DAYS EVEN ON CLUSTER. SO YOU CAN REDUCE ON THE NUMBER OF ELEMENTS IN THE LIST
LOCI = list(c(1,0,0, 0, 0), c(0,1,0,0, 0),c(0,0,1,0, 0),c(0,0,0,1, 0),c(0,0,0,0, 1),
            c(1/2,1/2,0,0, 0),  c(1/2,0,1/2,0, 0), c(1/2,0,0,1/2, 0), c(1/2,0,0,0, 1/2), c(0,1/2,1/2,0, 0), c(0,1/2,0,1/2, 0), c(0,1/2,0,0, 1/2),
            c(0,0,1/2,1/2, 0), c(0,0,1/2,0, 1/2), c(0,0,0,1/2, 1/2),
            c(1/3,1/3,1/3,0, 0), c(1/3,0,1/3,1/3, 0), c(1/3,0,0,1/3, 1/3), c(1/3, 1/3,0, 1/3, 0), c(1/3,1/3,0,0, 1/3),c(0,1/3,1/3,1/3, 0),c(0,0,1/3,1/3, 1/3),
            c(0,1/3,0,1/3,1/3), c(1/3,0,1/3,0,1/3), c(0,1/3,1/3,0,1/3),
            c(1/4,1/4,1/4,1/4, 0),c(1/4,1/4,1/4,0, 1/4), c(1/4, 1/4,0, 1/4, 1/4), c(1/4,0,1/4,1/4, 1/4), c(0,1/4,1/4,1/4, 1/4),
            c(1/5,1/5,1/5,1/5, 1/5))


ts.folder <- file.path(getwd(), paste("/MVP_ResultsFolder", sep = "" ))
if(!dir.exists(ts.folder)){
  dir.create(ts.folder)
}

setwd(ts.folder)


# Inital population size = Carrying capacity. 
Founderpop = round(seq(10, 100, length.out = 10),0)
# Used  round(seq(10, 3500, length.out = 350),0) that take time
capacity = Founderpop

#Initial starting allele frequency. HA, FS and FD is 0.5 and ID and MA is 0.95 
Startfreq = c(0.5,0.5,0.5,0.95,0.95)

# Do not change these set parameters! Not used in this paper!
initStoch = 1
Envmax = 1
Envrate = 1
capaStoch  = 0
growthStoch = 0

rate = log(1.53) # intrinsic growth rate
Tmax = 400 # Maximum simulation time
# Used Tmax = 4000
kk = 20 # Discretization of allele frequency. reciprocal or step length   
Tau = 1 # Discretization of time. Step length between time.
kapa = 50 # Generations in a complete cycle
Percent = 40 # parameter for FS
#------------------------------------------------------------------------
tol = 1e-39 # used during taking of inverse of the matrix 1 + alpha*RR
#Selection coeficients 
Mutaccsa = 0.005
Inbsa = 0.05
Hetsa = 0.005
HetsA = 0.005
Freqsa = 0.005
FreqsA = 0.005
FlsA = 0.005
Flsa = 0.005
# Dominance Coefficents
Inbhh = 0.01
Mutacchh = 0.5

# Mutation rates
mutrate = round(seq(0.00005, 0.0002, length.out = 2),5)
#Used mutrate = round(seq(0.00005, 0.002, length.out = 10),5)
mut_a = mutrate
mut_A = mutrate
inmut_a = mutrate/100
mamut_a = mutrate/100

#------------------------------------------------------
conames = list()
NNames = c("HA","FS","FD","ID","MA")

for (ll in 1:length(LOCI)){
  L=c()
  for(i in 1:length(LOCI[[ll]])){
    L = c(L,LOCI[[ll]][i])
  }
  L[which(L>0)] = which(L>0)
  
  conames[ll] = paste(NNames[L[1]],NNames[L[2]],NNames[L[3]],NNames[L[4]],NNames[L[5]], sep="")
}
#conames

# matt Matrix for storing MVPs. 1st column is the mutation rate 
#and other columns are for the different combinations of selection mechanisms
matt = matrix(NA, nrow = length(mutrate), ncol = length(LOCI)+1)

colnames(matt) = c("mut",unlist(conames))
matt[,1] = mutrate
Mupar = 1:length(mutrate)

#looping through combination of mechanisms
for (mech in 1:length(LOCI)) {
  print(c("mechcomb",conames[mech]))
  
  lociRatioVector = LOCI[[mech]]
  # Total number of loci
  if(length(lociRatioVector>0)==3){
    Numberofloci = 90
  }else{
    Numberofloci=100
  }
  # Parallelization for mutation rates
  RRES =  mclapply(Mupar, FUN = function(mut){
    
    for (pop in 1:length(capacity)){# looping through carrying capacities
      
      RRes = ApproxMixFunc(startfreq = Startfreq, numberofloci = Numberofloci, founderpop = Founderpop[pop], Capacity = capacity[pop], Rate = rate, tmax = Tmax, tau = Tau, KK = kk, 
                           Kapa = kapa, percent = Percent, Mut_a = mut_a[mut], inMut_a = inmut_a[mut], maMut_a = mamut_a[mut],
                           Mut_A = mut_A[mut], envmax = Envmax, envrate = Envrate, InitStoch = initStoch, CapaStoch = capaStoch, GrowthStoch = growthStoch, hetsa = Hetsa, hetsA = HetsA, flsa = Flsa, flsA = FlsA,  
                           freqsa = Freqsa,
                           freqsA = FreqsA, inbsa = Inbsa, inbhh=Inbhh, LociRatioVector = lociRatioVector, Tol = tol, mutaccsa = Mutaccsa, mutacchh=Mutacchh)
      
      
      if(RRes[Tmax,1]>0){ # Checking if there is at least 1 individual
        mattt = c(mutrate[mut],capacity[pop])
        print(c("mechcomb",conames[mech], "Mut", mutrate[mut], "pop", capacity[pop]))
        
        break
      }else{
        mattt = c(mutrate[mut],NA)
        
      }
    }
    return(mattt)
    
  }, mc.preschedule = FALSE, mc.cores = cores)
  
  # Rearranging MVPs from the above parallelization
  # matt Matrix for storing MVPs. 1st column is the mutation rate 
  #and other columns are for the different combinations of selection mechanisms
  LL = unlist(RRES)[seq(1,2*length(mutrate),2)]
  LLorder = LL[order(LL)]
  
  LLmvp = unlist(RRES)[seq(2,2*length(mutrate),2)]
  
  MVPorder = LLmvp[order(LL)]
  matt[1:length(MVPorder),mech+1] = MVPorder
}
#Storing mvp matrix results as csv file for later use.
write.csv(matt, file = paste(ts.folder,"/MVPfixedlociDAA_","fit_r0log_",round(exp(rate),4),"_hsa_", Hetsa, "_hsA_", HetsA, "_flsa_", Flsa, "_flsA_", FlsA, "_frsa_", Freqsa, 
                              "_frsA_", FreqsA, "_insa_",Inbsa, "_inh_", Inbhh, "_masa_", Mutaccsa, "_mah_", Mutacchh, "_kapa_", kapa,"tmax", Tmax,  ".csv", sep = ""))


setwd("~/MVP_GENETICPROBLEMS") # Set the directory again!!
```
