## Supplementary material for "Modeling minimum viable population size with multiple genetic problems of small populations": R codes used to simulate results: Execute_DiffusionApproximation_VariableLoci.R

```
rm(list=ls())
# Packages mainly for parallelisation
library("doMC")
cores=detectCores()
cl <- makeCluster(cores) 
registerDoMC(cores)
require(reshape2)


setwd("~/MVP_GENETICPROBLEMS")
source("MainFunct_DiffusionApproximation.R")

ts.folder <- file.path(paste(getwd(),"/MVP_ResultsFolder", sep = "" ))
if(!dir.exists(ts.folder)){
  dir.create(ts.folder)
}

setwd(ts.folder)

#Initial starting allele frequency. HA, FS and FD is 0.5 and ID and MA is 0.95 
Startfreq = c(0.5,0.5,0.5,0.95,0.95)

# Do not change these set parameters! Not used in this paper!
initStoch = 1
Envmax = 1
Envrate = 1
capaStoch  = 0
growthStoch = 0
#-------------------------------------------

rate = log(1.53) # intrinsic growth rate
Tmax = 400 # Maximum simulation time;
#Used Tmax = 4000
kk = 20 # Discretization of allele frequency. reciprocal or step length   
Tau = 1 # Discretization of time. Step length between time.
kapa = 50 # Generations in a complete cycle
Percent = 40 # parameter for FS
#------------------------------------------------------------------------
tol = 1e-39 # used during taking of inverse of the matrix 1 + alpha*RR
#Selection coeficients 
Mutaccsa = 0.005
Inbsa = 0.05
Hetsa = 0.005
HetsA = 0.005
Freqsa = 0.005
FreqsA = 0.005
FlsA = 0.005
Flsa = 0.005
# Dominance Coefficents
Inbhh = 0.01
Mutacchh = 0.5

# Mutation rates
mutrate = 0.0001
mut_a = mutrate
mut_A = mutrate
inmut_a = mutrate/100
mamut_a = mutrate/100


# Inital population size = Carrying capacity
Founderpop = round(seq(10, 200, by = 20),0) 
# Used Founderpop = round(seq(10, 150000, by = 20),0)#WARNING! THIS CAN TAKE SEVERAL HOURS OR DAYS TO COMPLETE

capacity = Founderpop

   
#------------------------------------------------------------
#Here, we just have different combinations of 2 selection mechanisms in a matrix with 2 columns. The rows also include cases with same mechanism e.g. row with c(1,1) implies only HA
vv = c(1,2,3,4,5)# represents selection mechanisms. 1 - HA, 2 - FS, 3 - FD, 4 - ID and 5 - MA
yy = t(combn(vv,2))
yy1 = matrix(c(yy[,2],yy[,1]), ncol = 2, byrow = FALSE)
yy2 = matrix(rep(c(1:5),each=2),ncol = 2, byrow = TRUE)
YY1 = rbind(yy2,yy,yy1)# All different combinations together.
#YY = YY1

DAA_MVPaddedloci = function(Minloci, DiffPairs, addloci){

XX = 1:length(addloci)

conames = list()
NNames = c("HA","FS","FD","ID","MA")
for (ppair in 1:nrow(DiffPairs)){
conames[ppair] = paste(NNames[DiffPairs[ppair,][1]],NNames[DiffPairs[ppair,][2]], sep="-")
}

# Matrix for storing MVPs. Column 1 is the number of loci added and the remaining columns represent different mechanisms as indicated by column names
matt = matrix(NA, ncol = nrow(DiffPairs)+1, nrow = length(XX))
colnames(matt) = c("addloci",unlist(conames))
matt[,1] = addloci 


for (pair in 1:nrow(DiffPairs)) {
print(c("mechcombination",DiffPairs[pair,][1],DiffPairs[pair,][2]))
RRES =  mclapply(XX, FUN = function(lo){
  Numberofloci = Minloci+addloci[lo]
  ll = c(Minloci/Numberofloci,addloci[lo]/Numberofloci)
  lociRatioVector = rep(0,5) # In the order HA,FS,FD,ID and MA
  cx = DiffPairs[pair,]
if(cx[1]!=cx[2]){
  lociRatioVector[cx] = ll
  }else{
    lociRatioVector[cx[1]] = 1
  }
  
  
  for (pop in 1:length(capacity)){
    
    RRes = ApproxMixFunc(numberofloci = Numberofloci,startfreq=Startfreq, founderpop = Founderpop[pop], Capacity = capacity[pop], Rate = rate, tmax = Tmax, tau = Tau, KK = kk, Kapa = kapa, percent = Percent, Mut_a = mut_a, inMut_a = inmut_a, maMut_a = mamut_a,
                         Mut_A = mut_A, envmax = Envmax, envrate = Envrate, InitStoch = initStoch, CapaStoch = capaStoch, GrowthStoch = growthStoch, hetsa = Hetsa, hetsA = HetsA, flsa = Flsa, flsA = FlsA, freqsa = Freqsa,
                         freqsA = FreqsA, inbsa = Inbsa, inbhh=Inbhh, LociRatioVector = lociRatioVector, Tol = tol, mutaccsa = Mutaccsa, mutacchh = Mutacchh)
    
    
    if(RRes[Tmax,1]>0){# Checking if there is survival
   mattt = c(addloci[lo],capacity[pop])
      print(c("mechcomb",DiffPairs[pair,][1],DiffPairs[pair,][2], "Mut", mut_a, "addloci",addloci[lo], "pop", capacity[pop]))
      
      break
    }else{
     mattt = c(addloci[lo],NA)

      #print(c("addloci",addloci[lo], "notpop", capacity[pop]))
    }
  }
  return(mattt)}, mc.preschedule = FALSE, mc.cores = cores)
#Rearranging MVPs from the above parallelization
LL = unlist(RRES)[seq(1,2*length(addloci),2)]
LLorder = LL[order(LL)]

LLmvp = unlist(RRES)[seq(2,2*length(addloci),2)]

MVPorder = LLmvp[order(LL)]
matt[1:length(MVPorder),pair+1] = MVPorder

}

# Storing MVP matrix for later use.
#write.csv(matt, file = paste(ts.folder,"/MVPaddedloci_", "_mut_", mut_a, "balsel",Hetsa, "inbsa", Inbsa, "inbhh",Inbhh, "mutaccsa", Mutaccsa, "mutacchh", Mutacchh,"tmax",Tmax, ".csv", sep = ""))
return(matt)
}


YYabove100 = YY1 # All possible pairs
YYbelow100 = YY1[1:5,] # Only isolated cases to work for total loci at most 100
addlociAbove100 = seq(0,500,by = 50) # Number of loci for different populations. Should be at least 1. This is added to some MinLoci.
# Used addlociAbove100 = seq(0,1000,by = 5)
addlocibelow100 = seq(1,100,by = 2) # Number of loci for different populations. Should be at least 1. This is added to some MinLoci.
# Used addlocibelow100 = seq(1,100,by = 2)
minlociabove100 = 100 # Minimum number of loci 
minlocibelow100 = 1 # Minimum number of loci for populations with a maximum of 100 loci in total


MVPdaaAbove100 = DAA_MVPaddedloci(Minloci = minlociabove100, DiffPairs = YYabove100, addloci = addlociAbove100)
# One can save the MVPs and use them later. 
write.csv(MVPdaaAbove100, file = paste(ts.folder,"/MVPvariablelociDAAabove100", "_mut_", mut_a, "balsel",Hetsa, "inbsa", Inbsa, "inbhh",Inbhh, "mutaccsa", Mutaccsa, "mutacchh", Mutacchh,"tmax",Tmax, ".csv", sep = ""))

MVPdaabelow100 = DAA_MVPaddedloci(Minloci = minlocibelow100, DiffPairs = YYbelow100, addloci = addlocibelow100)
# One can save the MVPs and use them later.
write.csv(MVPdaabelow100, file = paste(ts.folder,"/MVPvariablelociDAAbelow100", "_mut_", mut_a, "balsel",Hetsa, "inbsa", Inbsa, "inbhh",Inbhh, "mutaccsa", Mutaccsa, "mutacchh", Mutacchh,"tmax",Tmax, ".csv", sep = ""))

setwd("~/MVP_GENETICPROBLEMS")# Set the working directory again
```
