## Supplementary material for "Modeling minimum viable population size with multiple genetic problems of small populations": R codes used to simulate results: Execute_IBM.R

```
rm(list=ls())
# Packages 
library("doMC")
cores=detectCores()
registerDoMC(cores)
require(reshape2)
library(plyr)

setwd("~/MVP_GENETICPROBLEMS")
source("MainFunct_IBM.R")

replicates = 1:4 # Number of replicates
# Used replicates = 1:60 # Number of replicates
lociRatioVector = c(1,0,0,0,0)# HA, FS,FD,ID,MA, must sum to 1.

mutrate = round(seq(0.001, 0.002, length.out = 2), 5)
# Used mutrate = round(seq(0, 0.002, length.out = 10), 5)
Founderpop <- round(seq(10, 100, length.out = 10),0) # starting population size for HA
# Used Founderpop <- round(seq(10, 3500, length.out = 350),0) # starting population size for HA

capacity <- Founderpop # Founder population size denoted as N_0 in the text used as carrying capacity


if(length(which(lociRatioVector>0))==3){
  Numberofloci = 99
}else{
  Numberofloci = 100
}


Tmax = 100 # max. simulation time 
# Used Tmax = 4000 # max. simulation time 
x0 = 0.5# Initail frequency of allele A for all mechanisms

# Selection coefficients
Hetsa = 0.005
HetsA = Hetsa
Fluctsa = 0.005
FluctsA = Fluctsa
Freqsa = 0.005
FreqsA = Freqsa
idsa = 0.005
masa = 0.005

# Dominance coefficients
id_h = 0.0
ma_h = 0.5

#Mutation rates
hetmut_A = mutrate
hetmut_a = mutrate
fluctmut_A = mutrate
fluctmut_a = mutrate
freqmut_A = mutrate
freqmut_a = mutrate
inbmut_A = mutrate
inbmut_a = mutrate/100
mamut_A = mutrate
mamut_a = mutrate/100

rate = log(1.53) # Intrinsic growth rate 
Deathrate = 0
Percent = 40 #Percent is a parameter in heterozygote genotype fitness for FS between 0 and 100 (used 40 throught).
kapa = 50 # Number of generations in a complete cycle

Threshold = 1 # Set threshold to stop running. Here all 100% of replicates must survive to stop increasing capacity.

# Do not change these parameters. They are silent in this study
initStoch = 1
Envrate = 0
Envmax = 1
capaStoch = 0
growthStoch = 0

ts.folder <- file.path( paste(getwd(),"/MVP_ResultsFolder", sep = "" ))
if(!dir.exists(ts.folder)){
  dir.create(ts.folder)
}

setwd(ts.folder)


ts.folderIBM <- file.path(ts.folder,paste("IBM_TotLoci_",Numberofloci, "_hetloci_", round(lociRatioVector[1],2),  "_fluctloci_", round(lociRatioVector[2],2),
                                     "_freqloci_",  round(lociRatioVector[3],2), "_inbloci_", round(lociRatioVector[4],2), "_mutaccloci_", round(lociRatioVector[5],2), sep = ""  ))
if(!dir.exists(ts.folderIBM)){
  dir.create(ts.folderIBM)
}

setwd(ts.folderIBM)

# Here we try to run the IBM for the different replicates and other parameters 
# NOT, THESE LOOPS MAY TAKE A LONG TIME AND MAY NEED ENOUGH COMPUTATIONAL POWER IF NUMBER OF REPLICATES IS HIGH AS WELL AS MANY VALUES OF CARRYING CAPACITY. 
for (mut in 1:length(mutrate)) {
  for (pop in 1:length(Founderpop)){    
    foreach(Rep = replicates, .combine='cbind') %dopar% {
      
      SIMU <- Main_IBM(founderpop =  Founderpop[pop], numberofloci =  Numberofloci, Capacity =  capacity[pop], HAmut_a =  hetmut_a[mut], HAmut_A =  hetmut_A[mut], FSmut_a =  fluctmut_a[mut],
                  FSmut_A =  fluctmut_A[mut], FDmut_a =  freqmut_a[mut], FDmut_A =  freqmut_A[mut], IDmut_a =  inbmut_a[mut], IDmut_A =  inbmut_A[mut], MAmut_a =  mamut_a[mut],
                  MAmut_A =  mamut_A[mut], tmax =  Tmax, x0 =  x0, hetsa =  Hetsa, hetsA =  HetsA, fluctsa =  Fluctsa, fluctsA =  FluctsA,
                  freqsa =  Freqsa, freqsA =  FreqsA, IDsa =  idsa, MAsa =  masa, ID_h =  id_h, MA_h =  ma_h, Rate =  rate,
                  deathrate =  Deathrate, percent =  Percent, Kapa =  kapa, InitStoch =  initStoch, envrate =  Envrate,
                  CapaStoch =  capaStoch, GrowthStoch =  growthStoch, LociRatioVector = lociRatioVector)
      
      
      # We write all the results here as csv file for each replicate as long as the population went extinct in less than the set survival threshold.
      write.csv(SIMU, file = paste(ts.folderIBM, "/Mixloci_r0_log",exp(rate), "_N0_",  Founderpop[pop] , "_K_",  capacity[pop], "_sel_", HetsA, "_insa_",  idsa,  "_masa_",  masa, "_inh_",  id_h, "_mah_",
                                    ma_h, "_muA_",  hetmut_A[mut], "_mua_",  hetmut_a[mut], "_n_",  Numberofloci, "_hlc_", round(lociRatioVector[1],2),  "_fllc_", round(lociRatioVector[2],2),  "_frlc_",
                                   round(lociRatioVector[3],2),  "_inlc_", round(lociRatioVector[4],2),  "_malc_", round(lociRatioVector[5],2), "_kapa_",  kapa,
                                   "_rep", Rep, ".csv", sep = ""
      ))
    }
    
    
    #-------------------------------------------
    Pattern = paste( "Mixloci_r0_log",exp(rate), "_N0_",  Founderpop[pop] , "_K_",  capacity[pop], "_sel_", HetsA, "_insa_",  idsa,  "_masa_",  masa, "_inh_",  id_h, "_mah_", ma_h, "_muA_", 
                         hetmut_A[mut], "_mua_",  hetmut_a[mut], "_n_",  Numberofloci, "_hlc_", round(lociRatioVector[1],2),  "_fllc_", round(lociRatioVector[2],2),  "_frlc_",
                                   round(lociRatioVector[3],2),  "_inlc_", round(lociRatioVector[4],2),  "_malc_", round(lociRatioVector[5],2), "_kapa_",  kapa,
                      "*", sep = "")
    #We check if our persistence threshold is satisfied at the end of simulation time.
    temp = list.files(path = getwd(), pattern=Pattern)
    myfiles = lapply(temp, read.csv)
    X1 = dplyr::bind_rows(myfiles)
    Xpopsize = matrix(X1$POPSIZE, nrow = Tmax, ncol = length(replicates), byrow = FALSE)
    existrep = length(which(Xpopsize[Tmax,]>0))/length(replicates)
    
    if(existrep>=Threshold){ # Checking
      print(c("mut",mutrate[mut],"pop", Founderpop[pop]))
      break
    }else{
      print(c("mut",mutrate[mut],"notpop", Founderpop[pop]))
     }

    #------------------------------------------------
    
  }
  
}


# Now combining the trajectories together


ts.folder1 <- file.path(ts.folderIBM,paste("CapacityMutation", sep = ""  ))
if(!dir.exists(ts.folder1)){
  dir.create(ts.folder1)
}


for (pop in 1:length(Founderpop)){
  for (mut in 1:length(mutrate)){

    Pattern = paste( "Mixloci_r0_log",exp(rate), "_N0_",  Founderpop[pop] , "_K_",  capacity[pop], "_sel_", HetsA, "_insa_",  idsa,  "_masa_",  masa, "_inh_",  id_h, "_mah_", ma_h, "_muA_",
                     hetmut_A[mut], "_mua_",  hetmut_a[mut], "_n_",  Numberofloci, "_hlc_", round(lociRatioVector[1],2),  "_fllc_", round(lociRatioVector[2],2),  "_frlc_",
                     round(lociRatioVector[3],2),  "_inlc_", round(lociRatioVector[4],2),  "_malc_", round(lociRatioVector[5],2), "_kapa_",  kapa,
                      "*", sep = "")
    
    
    temp = list.files(path = getwd(), pattern=Pattern)
    if(length(temp)>0){
      myfiles = lapply(temp, read.csv)
      X1 = dplyr::bind_rows(myfiles)
      
      Xpopsize = matrix(X1$POPSIZE, nrow = Tmax, ncol = length(replicates), byrow = FALSE)
      Xmeanfit = matrix(X1$MEANFIT, nrow = Tmax, ncol = length(replicates), byrow = FALSE)
      
      # Population size and viability fitness trajectories for all mutation rates, replicates and capacities
      write.csv(Xpopsize, file = paste(ts.folder1,"/POPSIZE_r0_log",exp(rate ),   "_Capacity_", capacity[pop], "_hsa_",  Hetsa, "_hsA_",  HetsA, "_flsa_",  Fluctsa,
                                       "_flsA_",  FluctsA, "_frsa_",  Freqsa, "_frsA_",  FreqsA, "_insa_",  idsa,  "_masa_",  masa, "_inh_",  id_h, "_mah_",  ma_h,
                                       "_muA_",  hetmut_A[mut], "_mua_",  hetmut_a[mut], "_n_",  Numberofloci, "_hlc_", round(lociRatioVector[1],2),  "_fllc_", round(lociRatioVector[2],2),  "_frlc_",
                                       round(lociRatioVector[3],2),  "_inlc_", round(lociRatioVector[4],2),  "_malc_", round(lociRatioVector[5],2), "_kapa_",  kapa,  
                                       "_tmax_", Tmax, ".csv", sep = ""))
      
      write.csv(Xmeanfit, file = paste(ts.folder1,"/MEANFIT_r0_log",exp(rate ),   "_Capacity_", capacity[pop], "_hsa_",  Hetsa, "_hsA_",  HetsA, "_flsa_",  Fluctsa,
                                       "_flsA_",  FluctsA, "_frsa_",  Freqsa, "_frsA_",  FreqsA, "_insa_",  idsa,  "_masa_",  masa, "_inh_",  id_h, "_mah_",  ma_h,
                                       "_muA_",  hetmut_A[mut], "_mua_",  hetmut_a[mut], "_n_",  Numberofloci, "_hlc_", round(lociRatioVector[1],2),  "_fllc_", round(lociRatioVector[2],2),  "_frlc_",
                                       round(lociRatioVector[3],2),  "_inlc_", round(lociRatioVector[4],2),  "_malc_", round(lociRatioVector[5],2), "_kapa_",  kapa,
                                       "_tmax_", Tmax, ".csv", sep = ""))
    }
  }
}
```
