## Supplementary material for "Modeling minimum viable population size with multiple genetic problems of small populations": R codes used to simulate results: Execute_MarkovChain_POPequiApproach_FixedLoci.R

```
#numberofloci is number of loci, used mainly 100
#LociRatioVector is a vector of loci ratios in the order HA, FS, FD, ID, MA. The vector should sum to 1.
#N_eq is the carrying capacity.
#Rate is the growth rate. Used log(1.53).
#MAsA = 0 is the selection coeff. against AA for mutation accumulation.
#MAsa is the selection coeff. against aa for mutation accumulation.
#MA_h=0 is the dominance coefficient of allele a in mutation accumulation.
#bitaID is the mutation rate for a to A in inbreeding depression (ID).
#bitaMA is the mutation accumulation for a to A in mutation accumulation.
#IDsa is the selection coeff. against aa for inbreeding depression.
#ID_h is the dominance coeff. of a allele in inbreeding depression.
#FDsA is the selection coeff. oagainst AA for negative frequency-dependent selection (FD).
#FDsa is the selection coeff. against aa for FD.
#FSsa is the selection coeff. against aa for fluctuating selection with reversal of dominance (FS).
#FSsA is the selection coeff. against AA for FS.
#percent is a parameter in heterozygote genotype fitness for FS between 0 and 1 (used 0.4 throught).
#kapa is number of generations in a complete cyle in FS (used 50).
#HAsA is the selection coeff. against AA for heterozygote advantage (HA).
#HAsa is the selection coeff. against aa for HA.
#mu is the mutation rate from A to a, for balancing selection mechanisms.
#bita is the mutation rate from a to A, for balancing selection mechanisms (Used mu = bita).

#--------------------------------------------------------------
rm(list=ls())
# Packages mainly for parallelisation
library("doMC")
cores=detectCores()
registerDoMC(cores)
require(reshape2)

setwd("~/MVP_GENETICPROBLEMS")
source("MainFunct_MarkovChain_POPequiApproach_FixedLoci.R")

# List of selection mechanism combintation. Each item is a vector of length 5. Within the vector, the 1st, 2nd, 3rd, 4th and 5th elements are the proportion 
# of loci under HA, FS, FD, ID and MA. e.g c(1,0,0,0,0) implies 100% HA loci, c(1/5,1/5,1/5,1/5,1/5) implies 1/5th loci under each mechanism
LOCI = list(c(1,0,0, 0, 0), c(0,1,0,0, 0),c(0,0,1,0, 0),c(0,0,0,1, 0),c(0,0,0,0, 1),
           c(1/2,1/2,0,0, 0),  c(1/2,0,1/2,0, 0), c(1/2,0,0,1/2, 0), c(1/2,0,0,0, 1/2), c(0,1/2,1/2,0, 0), c(0,1/2,0,1/2, 0), c(0,1/2,0,0, 1/2),
           c(0,0,1/2,1/2, 0), c(0,0,1/2,0, 1/2), c(0,0,0,1/2, 1/2),
           c(1/3,1/3,1/3,0, 0), c(1/3,0,1/3,1/3, 0), c(1/3,0,0,1/3, 1/3), c(1/3, 1/3,0, 1/3, 0), c(1/3,1/3,0,0, 1/3),c(0,1/3,1/3,1/3, 0),c(0,0,1/3,1/3, 1/3),
          c(0,1/3,0,1/3,1/3), c(1/3,0,1/3,0,1/3), c(0,1/3,1/3,0,1/3),
          c(1/4,1/4,1/4,1/4, 0),c(1/4,1/4,1/4,0, 1/4), c(1/4, 1/4,0, 1/4, 1/4), c(1/4,0,1/4,1/4, 1/4), c(0,1/4,1/4,1/4, 1/4),
          c(1/5,1/5,1/5,1/5, 1/5))


#--------------------------------------------------------------
Mu  <- round(seq(0.00005, 0.0002, length.out = 2),5) #mutation rate
# Used Mu  <- round(seq(0.00005, 0.002, length.out = 10),5) #mutation rate
Bita = Mu
BitaID = Mu/100
BitaMA = Mu/100
n_eq  <- round(seq(10, 100,length.out = 10),0)
# Used n_eq  <- round(seq(10, 3500,length.out = 350),0)
rate = log(1.53)

#selection coefficients
hasA = 0.005
hasa = 0.005
fssa = 0.005
fssA = 0.005
fdsA = 0.005
fdsa = 0.005
idsa = 0.05
masA = 0
masa = 0.005
Percent = 40
Kapa = 50 # Number of generations in a complete cycle

# Dominance coefficients for ID and MA
id_h = 0.01
ma_h = 0.5

ts.folder <- file.path(paste(getwd(),"/MVP_ResultsFolder", sep = "" ))
if(!dir.exists(ts.folder)){
  dir.create(ts.folder)
}

setwd(ts.folder)

#------------------------------------------------------
conames = list()
NNames = c("HA","FS","FD","ID","MA")

for (ll in 1:length(LOCI)){
  L=c()
  for(i in 1:length(LOCI[[ll]])){
    L = c(L,LOCI[[ll]][i])
  }
  L[which(L>0)] = which(L>0)
  
  conames[ll] = paste(NNames[L[1]],NNames[L[2]],NNames[L[3]],NNames[L[4]],NNames[L[5]], sep="")
}

# Matrix for storing MVPs. 1st column is the mutation rate 
#and other columns are for the different combinations of selection mechanisms
matt = matrix(NA, nrow = length(Mu), ncol = length(LOCI)+1) 
colnames(matt) = c("mut",unlist(conames))
matt[,1] = Mu 
Mupar = 1:length(Mu)

for (mech in 1:length(LOCI)) { # loop for the different combinations
  print(c("mechcomb",conames[mech]))
  
  lociRatioVector = LOCI[[mech]]
  # Fixed number of loci. 
  if(length(lociRatioVector>0)==3){
    Numberofloci = 90
  }else{
    Numberofloci=100
  }
  
  RRES =  mclapply(Mupar, FUN = function(mut){
    
    for (pop in 1:length(n_eq)){
      
      FF =  MainFuncPois(numberofloci = Numberofloci, LociRatioVector = lociRatioVector, N_eq = n_eq[pop], Rate = rate, MAsA = masA, MAsa = masa, MA_h=ma_h,
                         IDsa = idsa, ID_h=id_h, FDsA =fdsA, FDsa = fdsa, FSsa = fssa, FSsA = fssA, percent = Percent, kapa = Kapa, HAsA = hasA,
                         HAsa = hasa, mu =Mu[mut], bita = Bita[mut], bitaID = BitaID[mut], bitaMA = BitaMA[mut])
      
      if(FF[1]>=1){
        mattt = c(Mu[mut],n_eq[pop])
        print(c("mechcomb",conames[mech], "Mut", Mu[mut], "pop", n_eq[pop]))
        
        break
      }else{
        mattt = c(Mu[mut],NA)
        #print(c("addloci",addloci[lo], "notpop", capacity[pop]))
      }
    }
    return(mattt)
    
  }, mc.preschedule = FALSE, mc.cores = cores)
  # Rearranging MVPs from parallelization above
  LL = unlist(RRES)[seq(1,2*length(Mu),2)]
  LLorder = LL[order(LL)]
  
  LLmvp = unlist(RRES)[seq(2,2*length(Mu),2)]
  
  MVPorder = LLmvp[order(LL)]
  matt[1:length(MVPorder),mech+1] = MVPorder
}

# Store the MVP matrix in the working folder
write.csv(matt, file = paste(ts.folder,"/MVPfixedlociMCAEqui_","fit_r0log_",round(exp(rate),4),"_hsa_", hasa, "_hsA_", hasA, "_flsa_", fssa,
"_flsA_", fssA, "_frsa_", fdsa, "_frsA_", fdsA, "_insa_",idsa, "_inh_", id_h, "_masa_", masa, "_masA_", masA, "_mah_", ma_h, "_kapa_", Kapa, ".csv", sep = ""))

setwd("~/MVP_GENETICPROBLEMS") # Set the working directory again
```
