## Supplementary material for "Modeling minimum viable population size with multiple genetic problems of small populations": R codes used to simulate results: Execute_MarkovChain_POPequiApproach_VariableLoci.R

```
#numberofloci is number of loci
#LociRatioVector is a vector of loci ratios in the order HA, FS, FD, ID, MA. The vector should sum to 1.
#N_eq is the carrying capacity.
#Rate is the growth rate. Used Rate = log(1.53).
#FDsA is the selection coefficient oagainst AA for negative frequency-dependent selection (FD).
#FDsa is the selection coefficient against aa for FD.
#FSsa is the selection coefficient against aa for fluctuating selection with reversal of dominance (FS).
#FSsA is the selection coefficient against AA for FS.
#percent is a parameter in heterozygote genotype fitness for FS between 0 and 100 (used percent = 40 throught).
#kapa is number of generations in a complete cyle in FS (used 50).
#HAsA is the selection coefficient against AA for heterozygote advantage (HA).
#HAsa is the selection coefficient against aa for HA.
#mu is the mutation rate from A to a, for balancing selection mechanisms and ID and MA.
#bita is the mutation rate from a to A, for balancing selection mechanisms (Used mu = bita).
#MAsA = 0 is the selection coefficient against AA for mutation accumulation and inbreeding depression.
#MAsa is the selection coefficient against aa for mutation accumulation.
#MA_h is the dominance coefficient of allele a in ID (used MA_h = 0.5).
#bitaID is the mutation rate for a to A in inbreeding depression (ID) (used bitaID = mu/100).
#bitaMA is the mutation accumulation for a to A in mutation accumulation (MA) (used bitaMA = mu/100).
#IDsa is the selection coefficient against aa for inbreeding depression.
#ID_h is the dominance coefficient of a allele in inbreeding depression.

rm(list=ls())
# Packages mainly for parallelisation
library("doMC")
cores=detectCores()
registerDoMC(cores)
require(reshape2)

setwd("~/MVP_GENETICPROBLEMS")
source("MainFunct_MarkovChain_POPequiApproach_VariableLoci.R")


ts.folder <- file.path(paste(getwd(), "/MVP_ResultsFolder", sep = "" ))
if(!dir.exists(ts.folder)){
  dir.create(ts.folder)
}

setwd(ts.folder)

# mutation rates
Mu  <- 0.0001 
Bita = Mu
BitaID = Mu/100
BitaMA = Mu/100

# Population sizes/carrying capacity vector 
n_eq  <- round(seq(10, 200, by = 20),0)
# Used n_eq  <- round(seq(10, 3150, by = 20),0)
rate = log(1.53) # intrinsic growth rate

# Selection coefficients
hasA = 0.005
hasa = 0.005
fssa = 0.005
fssA = 0.005
fdsA = 0.005
fdsa = 0.005
idsa = 0.05
masA = 0
masa = 0.005
#--------------
Percent = 40
Kapa = 50 # Number of generations in a complete cycle
# Dominance coeeficients for ID and MA
id_h = 0.01 
ma_h = 0.5
MechVector = rep(1,5)
NNames = c("HA","FS","FD","ID","MA") # Names of different selection mechanisms

# This calls the function that gives out mean per-locus offspring viability fitnes
OffspringViabiltyDifferentK = MainFuncPoisPeter(LociRatioVector = MechVector, N_eq = n_eq, MAsA = masA, MAsa = masa, MA_h=ma_h,
             IDsa = idsa, ID_h=id_h, FDsA =fdsA, FDsa = fdsa, FSsa = fssa, FSsA = fssA, percent = Percent, kapa = Kapa, HAsA = hasA,
             HAsa = hasa, mu =Mu, bita = Bita, bitaID = BitaID, bitaMA = BitaMA)

# One can store this fitness as a csv file and then load any time. If so, uncommend this line.
#write.csv(OffspringViabiltyDifferentK, file = paste(ts.folder,"/MCA_MechDistriPopSizes", "_mut_", Mu, "balsel",hasa, "inbsa", idsa, "inbhh",id_h, "mutaccsa", masa, "mutacchh", ma_h, ".csv", sep = ""))


#Here, we just have different combinations of 2 selection mechanisms in a matrix with 2 columns. The rows also include cases with same mechanism e.g. row with c(1,1) implies only HA
vv = c(1,2,3,4,5)# represents selection mechanisms. 1 - HA, 2 - FS, 3 - FD, 4 - ID and 5 - MA
yy = t(combn(vv,2))
yy1 = matrix(c(yy[,2],yy[,1]), ncol = 2, byrow = FALSE)
yy2 = matrix(rep(c(1:5),each=2),ncol = 2, byrow = TRUE)
YY1 = rbind(yy2,yy,yy1)# All different combinations together.

# Column names for the different combinations of selection mechanisms
MCA_MVPaddedloci = function(Minloci, DiffPairs, addloci){
conames = list()
for (ppair in 1:nrow(DiffPairs)){
  conames[ppair] = paste(NNames[DiffPairs[ppair,][1]],NNames[DiffPairs[ppair,][2]], sep="-")
}
#------------------------------------------------------

XX1 = 1:nrow(DiffPairs)

RRES = mclapply(XX1, FUN = function(pair){
#for (pair in 1:nrow(DiffPairs)) {
mattt = rep(NA,length(addloci))
for (lo in 1:length(addloci)){
  ll = c(Minloci, addloci[lo])# Total number of loci
  # for (pair in 1:nrow(DiffPairs)) {
    addlociRatioVector = rep(0,5)
    cx = DiffPairs[pair,]
    if(cx[1]!=cx[2]){
      addlociRatioVector[cx] = ll
      mechpo = which(addlociRatioVector>0)
      mechnumberloci = addlociRatioVector[mechpo] # Number of loci for the 1st and 2nd mechanism
      mech1 = OffspringViabiltyDifferentK[,mechpo[1]+1]^mechnumberloci[1]# Offspring viability from the Number of loci of the 1st mechanism 
      mech2 = OffspringViabiltyDifferentK[,mechpo[2]+1]^mechnumberloci[2]# Offspring viability from Number of loci of the 2nd mechanism
      absoluteF = exp(rate)*mech1*mech2 # Absolute fitness
     
      ttest = which(absoluteF>=1)
      if (length(ttest)>0){
        # matt[lo, pair+1] = n_eq[min(which(absoluteF>=1))]   
        mattt[lo] = n_eq[min(ttest)]
      }else{
        mattt[lo] = NA
      }
    }else{
      # Here for a case with only one mechanism
      addlociRatioVector[cx[1]] = sum(ll)
      mechpo = which(addlociRatioVector>0)
      mechnumberloci = addlociRatioVector[mechpo]# Number of loci for the only mechanism
      mech1 = OffspringViabiltyDifferentK[,mechpo[1]+1]^mechnumberloci[1]# Offspring viability fitness from the loci for the 1st and only mechanism
      absoluteF = exp(rate)*mech1
      ttest = which(absoluteF>=1)
      if (length(ttest)>0){
      #matt[lo,pair+1] =  n_eq[min(which(absoluteF1>=1))]
      mattt[lo] = n_eq[min(ttest)]
      }else{
        mattt[lo] = NA
      }
    }
    #----------------
    if(!is.na(mattt[lo])){
      print(c(pair,addloci[lo],mattt[lo]))
    }else{
      break
      }
}
  return(c(pair,mattt))
}, mc.preschedule = FALSE, mc.cores = cores)
# Rearranging MVP size from the parallelization above. The columns are for the different pairs and the rows are for the different number of loci added to the minimum number of loci

Mt = matrix(unlist(RRES), ncol = nrow(DiffPairs), nrow = length(addloci)+1)
LL = order(Mt[1,])
LLmvp1 = Mt[-1,LL]
LLmvp = cbind(addloci, LLmvp1)
colnames(LLmvp) = c("addloci",unlist(conames))

# One can save the MVPs and use them later. If so, uncommend the line below.
#write.csv(LLmvp, file = paste(ts.folder,"/MCA_MVPaddedloci", "_mut_", Mu, "balsel",hasa, "inbsa", idsa, "inbhh",id_h, "mutaccsa", masa, "mutacchh", ma_h, ".csv", sep = ""))

return(LLmvp)
}

YYabove100 = YY1 # All possible pairs
YYbelow100 = YY1[1:5,] # Only isolated cases to work for total loci at most 100
addlociAbove100 = seq(1,1000,by = 5) # Number of loci for different populations. Should be at least 1. This is added to some MinLoci.
addlocibelow100 = seq(1,100,by = 2) # Number of loci for different populations. Should be at least 1. This is added to some MinLoci.
minlociabove100 = 100 # Minimum number of loci 
minlocibelow100 = 1 # Minimum number of loci for populations with a maximum of 100 loci in total


MVPmcaAbove100 = MCA_MVPaddedloci(Minloci = minlociabove100, DiffPairs = YYabove100, addloci = addlociAbove100)
# One can save the MVPs and use them later. 
write.csv(MVPmcaAbove100, file = paste(ts.folder,"/MVPvariablelociMCAabove100", "_mut_", Mu, "balsel",hasa, "inbsa", idsa, "inbhh",id_h, "mutaccsa", masa, "mutacchh", ma_h, ".csv", sep = ""))

MVPmcabelow100 = MCA_MVPaddedloci(Minloci = minlocibelow100, DiffPairs = YYbelow100, addloci = addlocibelow100)
# One can save the MVPs and use them later.
write.csv(MVPmcabelow100, file = paste(ts.folder,"/MVPvariablelociMCAbelow100", "_mut_", Mu, "balsel",hasa, "inbsa", idsa, "inbhh",id_h, "mutaccsa", masa, "mutacchh", ma_h, ".csv", sep = ""))

setwd("~/MVP_GENETICPROBLEMS") # Set again the working directory
```
