## Supplementary material for "Modeling minimum viable population size with multiple genetic problems of small populations": R codes used to simulate results: MainFunct_DiffusionApproximation.R

```
# Main function for the Diffusion Approximation DAA

ApproxMixFunc = function(startfreq, numberofloci, founderpop, Capacity, Rate, tmax, tau, KK, Kapa, percent, Mut_a, Mut_A, inMut_a, maMut_a, envmax, envrate, InitStoch = 1, CapaStoch = 0, GrowthStoch = 0, hetsa, hetsA, flsa, flsA, freqsa, freqsA, inbsa, inbhh, LociRatioVector, Tol = 1e-39, mutaccsa, mutacchh){
  
  ENVSTOCHFUNC =  function(gentime, InitStoch=1, envmax=1, envrate=0){
    
    Stoch = (InitStoch*envmax*exp(envrate*(gentime-1)))/(envmax + InitStoch*(exp(envrate*(gentime-1))-1))
    
    return(Stoch)
  }
  #------------------------ Loci ratios 
  LociDistr = numberofloci*LociRatioVector
  HAloci = LociDistr[1]
  FSloci = LociDistr[2]
  FDloci = LociDistr[3]
  IDloci = LociDistr[4]
  MAloci = LociDistr[5]
  
  #--------------------------------------------
  h = 1/KK # spacing over [0,1]
  tt = seq(1,tmax,tau) # time steps
  ii = 0:KK
  xx = h*ii
  Alpha = tau/(2*h^2)
  
  hetHetero = numeric(length(tt)) # stores heterozygosity
  hetHetero[1] = 1#x0 # initial heterozygosity
  
  fluctHetero = numeric(length(tt)) # stores heterozygosity
  fluctHetero[1] = 1#x0 # initial heterozygosity
  
  freqHetero = numeric(length(tt)) # stores heterozygosity
  freqHetero[1] = 1#x0 # initial heterozygosity
  
  inbHetero = numeric(length(tt)) # stores heterozygosity
  inbHetero[1] = 1#x0 # initial heterozygosity
  
  mutaccHetero = numeric(length(tt)) # stores heterozygosity
  mutaccHetero[1] = 1#x0 # initial heterozygosity
  
  Ne = numeric(length(tt) + 1) # Population size
  Ne[1] = founderpop
  
  ffhet = matrix(0, nrow = c(KK+1), ncol = length(tt)) # stores density for each time step
  ffhet[round(startfreq[1]*KK+1,0),1] = 1/h # start with a dirac function represented by a pick at initial frequency
  
  fffluct = matrix(0, nrow = c(KK+1), ncol = length(tt)) # stores density for each time step
  fffluct[round(startfreq[2]*KK+1,0),1] = 1/h # start with a dirac function represented by a pick at initial frequency
  
  fffreq = matrix(0, nrow = c(KK+1), ncol = length(tt)) # stores density for each time step
  fffreq[round(startfreq[3]*KK+1,0),1] = 1/h # start with a dirac function represented by a pick at initial frequency
  
  ffinb = matrix(0, nrow = c(KK+1), ncol = length(tt)) # stores density for each time step
  ffinb[round(startfreq[4]*KK+1,0),1] = 1/h # start with a dirac function represented by a pick at initial frequency
  
  ffmutacc = matrix(0, nrow = c(KK+1), ncol = length(tt)) # stores density for each time step
  ffmutacc[round(startfreq[5]*KK+1,0),1] = 1/h # start with a dirac function represented by a pick at initial frequency

  
  #++++++++++++++++++++
  WWhet = numeric(length(tt))
  WWfluct = numeric(length(tt))
  WWfreq = numeric(length(tt))
  WWinb = numeric(length(tt))
  WWmutacc = numeric(length(tt))
  Fitness = numeric(length(tt))
  
  #------------------------- Selection coef. for fluct. selection --------------------------------
  fluctsA <- flsA*sin(2*pi*tt/Kapa) # selection coeff for genotype AA
  fluctsa <- flsa*sin(2*pi*tt/Kapa + pi)#smaxt*cos(Ti)^2# selection coeff for aa
  flucthh = 0.5-(percent/100)*sin(2*pi*tt/Kapa)#(smaxs - smaxs*percent/100)*cos(Ti)^10
  
  #---------------- Environmental Stochasticity ------------------------------------------------
  ENVSTOCHFUNC =  function(gentime, InitStoch=1, envmax=1, envrate=0){
    
    # Stoch = envmax*InitStoch/(InitStoch + (envmax-InitStoch)*exp(-envrate*(gentime-1))) 
    Stoch = (InitStoch*envmax*exp(envrate*(gentime-1)))/(envmax + InitStoch*(exp(envrate*(gentime-1))-1))
    return(Stoch)
  }
  
  
  #-------------------------Fitness functions -------------------------------------------------
  
  #------------- Heterozygote advantage (HA). ---------------------
  Wwhet = function(x, Sa, SA){
    #(2*fq*se*(1-fq) + (1-se))
    1 - (x^2*SA + Sa*(1-x)^2)
    
  }
  
  #----------------Fluctuating selection with dominance reversal (FS) ----------------------- 
  Wwfluct = function(x, Sa, SA, hh){
    (1-Sa + x*(Sa-SA)*(x+2*hh-2*x*hh))/(1+max(fluctsA,fluctsa))
    # (1-Sa + x*(Sa-SA)*(x+2*hh-2*x*hh))
  }
  
  #----------- Negative frequency-dependent selection ------------------------------------
  Wwfreq = function(x, Sa, SA){
    1 - (x^2*SA + Sa*(1-x)^2)
    
  }
  
  
  # -------------- Inbreeding depression (ID) -----------------------------------------
  Wwinb = function(x, Sa, hh){
    #1 - (1-x)*Sa*( x*(2*hh-1) + 1)
    wAA = 1; wAa = 1-hh*Sa; waa = 1-Sa
    
    Wfit = wAA*x^2 + 2*x*(1-x)*wAa + (1-x)^2*waa
    return(Wfit)
  }
  
  # ------------------ Mutation accumulation (MA)
  Wwmutacc = function(x, Sa, hh){
    #1 - (1-x)*Sa*( x*(2*hh-1) + 1)
    wAA = 1; wAa = 1-hh*Sa; waa = 1-Sa
    
    Wfit = wAA*x^2 + 2*x*(1-x)*wAa + (1-x)^2*waa
    return(Wfit)
  }
  
  ti = 1
  # Offspring viability in the first generation from eac mechanism per locus
  WWhet[ti] = (h/2)*ffhet[1,ti]*(Wwhet(x = xx[1], Sa= hetsa, SA = hetsA)) + (h/2)*ffhet[KK+1,ti]*(Wwhet(x=xx[KK+1], Sa= hetsa, SA = hetsA)) + sum(h*ffhet[-c(1,KK+1),ti]*(Wwhet(x=xx[2:KK], Sa= hetsa, SA = hetsA)))   #fitness from het. adva
  
  WWinb[ti] = (h/2)*ffinb[1,ti]*(Wwinb(x = xx[1], Sa = inbsa, hh = inbhh)) + (h/2)*ffinb[KK+1,ti]*(Wwinb(x=xx[KK+1], Sa = inbsa, hh = inbhh)) + sum(h*ffinb[-c(1,KK+1),ti]*(Wwinb(x=xx[2:KK], Sa = inbsa, hh = inbhh)))   #fitness from inbreeding
  
  WWfluct[ti] = (h/2)*fffluct[1,ti]*(Wwfluct(x=xx[1], Sa = fluctsa[ti], SA = fluctsA[ti], hh = flucthh[ti])) + (h/2)*fffluct[KK+1,ti]*(Wwfluct(x=xx[KK+1], Sa = fluctsa[ti], SA = fluctsA[ti], hh = flucthh[ti])) + sum(h*fffluct[-c(1,KK+1),ti]*(Wwfluct(x=xx[2:KK], Sa = fluctsa[ti], SA = fluctsA[ti], hh = flucthh[ti])))  #fitness from fluct. sel with rev. of dom.
  
  WWfreq[ti] = (h/2)*fffreq[1,ti]*(Wwfreq(x=xx[1], Sa= freqsa, SA= freqsA)) + (h/2)*fffreq[KK+1,ti]*(Wwfreq(x=xx[KK+1], Sa= freqsa, SA= freqsA)) + sum(h*fffreq[-c(1,KK+1),ti]*(Wwfreq(x=xx[2:KK], Sa= freqsa, SA= freqsA)))  #fitness from neg. freq. dep. sel.
  
  WWmutacc[ti] = (h/2)*ffmutacc[1,ti]*(Wwmutacc(x = xx[1], Sa = mutaccsa, hh = mutacchh)) + (h/2)*ffmutacc[KK+1,ti]*(Wwmutacc(x=xx[KK+1], Sa = mutaccsa, hh = mutacchh)) + sum(h*ffmutacc[-c(1,KK+1),ti]*(Wwmutacc(x=xx[2:KK], Sa = mutaccsa, hh = mutacchh)))   #fitness from mutation accumulation
  
  # Offspring viability via Multiplicative fitness across loci
  WWt = WWhet[ti]^HAloci * WWinb[ti]^IDloci * WWfluct[ti]^FSloci * WWfreq[ti]^FDloci * WWmutacc[ti]^MAloci
  WW = round(WWt,5)
  Fitness[ti] = WW
  
  # Hard carrying capacity applied

  Ne[ti+1] = Ne[ti]*min(Fitness[ti]*exp(Rate), Capacity/Ne[ti])
  
  
  #------------------------- Functions for infinitismal mean and variance --------------------
  # Variance for all
  VVall = function(x,N){
    x*(1-x)/(4*N)
  }  
  
  # Mean  for HA 
  MMhet = function(x, Sa, SA, hh, Mut_a, Mut_A){
    x*(1-x) * (Sa*(1-x)-SA*x ) * (1 + SA*x^2 + Sa*(1-x)^2 ) + Mut_a*(1-x) - Mut_A*x
  }
  
  # Mean for FS
  MMfluct = function(x, Sa, SA, hh, Mut_a, Mut_A){
    x*(1-x)*(x*(1-2*hh )+hh)*(1+Sa )*(Sa - SA ) - x^2*(1-x)*(Sa - SA)^2*(x*(1-2*hh)+hh)*(x*(1-2*hh)+2*hh) + Mut_a*(1-x) - Mut_A*x
  }
  
  # Mean for FD
  MMfreq = function(x, Sa, SA, hh, Mut_a, Mut_A){
    x*(1-x) * (Sa*(1-x)-SA*x) * (1+SA*x^2+Sa*(1-x)^2)/2 +  Mut_a*(1-x) - Mut_A*x
  }
  
  # Mean for ID
  MMinb = function(x, Sa, SA, hh, Mut_a = inMut_a, Mut_A){
    wAA = 1; wAa = 1-hh*Sa; waa = 1-Sa
    Wfit = wAA*x^2 + 2*x*(1-x)*wAa + waa*(1-x)^2
    pchange = (wAA*x^2 + x*(1-x)*wAa)/Wfit - x + inMut_a*(1-x) - Mut_A * x
    return(pchange)
  }
  
  # Mean for MA
  MMmutacc = function(x, Sa, SA, hh, Mut_a = maMut_a, Mut_A){
    wAA = 1; wAa = 1-hh*Sa; waa = 1-Sa
    Wfit = wAA*x^2 + 2*x*(1-x)*wAa + waa*(1-x)^2
    pchange = (wAA*x^2 + x*(1-x)*wAa)/Wfit - x + maMut_a*(1-x) - Mut_A * x
    return(pchange)
  }
  
  # The k+1 square matrix
  RR = function(x, Sa, SA, hh, MM, VV, N){
    
    V0 = VV(x = x[1], N = N) + h*MM(x = x[1], Sa = Sa, SA = SA, hh = hh, Mut_a= Mut_a, Mut_A = Mut_A)/2
    Vkminus1 = VV(x = x[length(x)-1], N = N) + h*MM(x = x[length(x)-1], Sa = Sa, SA = SA, hh = hh, Mut_a= Mut_a, Mut_A = Mut_A)/2
    U1 = -VV(x=x[2], N = N) + h*MM(x=x[2], Sa = Sa, SA = SA, hh = hh, Mut_a= Mut_a, Mut_A = Mut_A)/2
    Uk = -VV(x=x[length(x)], N = N) + h*MM(x=x[length(x)], Sa = Sa, SA = SA, hh = hh, Mut_a= Mut_a, Mut_A = Mut_A)/2
    
    Vi = VV(x = x[2:(length(x)-1)], N = N) + h*MM(x = x[2:(length(x)-1)], Sa = Sa, SA = SA, hh = hh, Mut_a= Mut_a, Mut_A = Mut_A)/2
    Viminus1 = VV(x = x[(2:(length(x)-1))-1], N = N) + h*MM(x = x[(2:(length(x)-1))-1], Sa = Sa, SA = SA, hh = hh, Mut_a= Mut_a, Mut_A = Mut_A)/2
    Ui = -VV(x = x[2:(length(x)-1)], N = N) + h*MM(x = x[2:(length(x)-1)], Sa = Sa, SA = SA, hh = hh, Mut_a= Mut_a, Mut_A = Mut_A)/2
    Uiplus1 = -VV(x = x[(2:(length(x)-1))+1], N = N) + h*MM(x = x[(2:(length(x)-1))+1], Sa = Sa, SA = SA, hh = hh, Mut_a= Mut_a, Mut_A = Mut_A)/2
    
    # filling the coefficient matrix M on LHS
    Coemat = matrix(0,nrow = (KK+1), ncol = (KK+1))
    Coemat[1,1] = 2*V0
    Coemat[1,2] = 2*U1
    Coemat[KK+1, KK] = -2*Vkminus1
    Coemat[KK+1, KK+1] = -2*Uk
    
    for (i in 2:KK){
      Coemat[i, i] = Vi[i-1]-Ui[i-1]
      Coemat[i,i-1] = -Viminus1[i-1]
      Coemat[i, i+1] = Uiplus1[i-1]
    }
    
    return(Coemat)
  }
  
  
  #----------------------------------------------------------------------------------------------
  for (ti in 1:(length(tt)-1)){ # generation loop
    
    if ( Ne[ti+1]>=1){
      
      # Allele distribution for HA
      if(HAloci>0){
        ffhet[, ti+1] = solve(diag(KK+1) + Alpha*RR(x=xx, Sa = hetsa, SA = hetsA, hh = 0, MM = MMhet, VV = VVall, N= Ne[ti+1]), tol = Tol) %*% (diag(KK+1) - Alpha*RR(x=xx, Sa = hetsa, SA = hetsA, hh = 0, MM = MMhet, VV = VVall, N= Ne[ti]))%*%ffhet[,ti]
       # ffhet[, ti+1] = ffhet[, ti+1]/sum(ffhet[, ti+1])
      }else{
        ffhet[, ti+1] = 0
      }
      
      # Allele distribution for FS
      if(FSloci>0){
        fffluct[, ti+1] = solve(diag(KK+1) + Alpha*RR(x=xx, Sa = fluctsa[ti+1], SA = fluctsA[ti+1], hh = flucthh[ti+1], MM = MMfluct, VV = VVall, N= Ne[ti+1]), tol = Tol) %*% (diag(KK+1) - Alpha*RR(x=xx, Sa = fluctsa[ti], SA = fluctsA[ti], hh = flucthh[ti], MM = MMfluct, VV = VVall, N= Ne[ti]))%*%fffluct[,ti]
      #  fffluct[, ti+1] = fffluct[, ti+1]/sum(fffluct[, ti+1])
      }else{
        fffluct[, ti+1] = 0
      }
      
      # Allele distribution for FD
      if(FDloci>0){
        fffreq[, ti+1] = solve(diag(KK+1) + Alpha*RR(x=xx, Sa = freqsa, SA = freqsA, hh = 1, MM = MMfreq, VV = VVall, N= Ne[ti+1]), tol = Tol) %*% (diag(KK+1) - Alpha*RR(x=xx, Sa = freqsa, SA = freqsA, hh = 1, MM = MMfreq, VV = VVall, N= Ne[ti]))%*%fffreq[,ti]
     #   fffreq[, ti+1] = fffreq[, ti+1]/sum(fffreq[, ti+1])
      }else{
        fffreq[, ti+1] = 0
      }
      
      # Allele distribution for ID
      if(IDloci>0){
        ffinb[, ti+1] = solve(diag(KK+1) + Alpha*RR(x=xx, Sa = inbsa, SA = 0, hh = inbhh, MM = MMinb, VV = VVall, N= Ne[ti+1]), tol = Tol) %*% (diag(KK+1) - Alpha*RR(x=xx, Sa = inbsa, SA = 0, hh = inbhh, MM = MMinb, VV = VVall, N= Ne[ti]))%*%ffinb[,ti]
    #    ffinb[, ti+1] = ffinb[, ti+1]/sum(ffinb[, ti+1])
      }else{
        ffinb[, ti+1] = 0
      }
      
      # Allele distribution for MA
      if(MAloci>0){
        ffmutacc[, ti+1] = solve(diag(KK+1) + Alpha*RR(x=xx, Sa = mutaccsa, SA = 0, hh = mutacchh, MM = MMmutacc, VV = VVall, N= Ne[ti+1]), tol = Tol) %*% (diag(KK+1) - Alpha*RR(x=xx, Sa = mutaccsa, SA = 0, hh = mutacchh, MM = MMmutacc, VV = VVall, N= Ne[ti]))%*%ffmutacc[,ti]
   #     ffmutacc[,ti+1] = ffmutacc[,ti+1]/sum(ffmutacc[,ti+1])
      }else{
        ffmutacc[, ti+1] = 0
      }    
      
    }else{
      
      break
    }
    
    #---------------------Heterozygosity for HA, FS, FD, ID and MA respectively---------------------------
    
    hetHetero[ti+1] = 1*(KK-1)*mean(h*ffhet[-c(1,KK+1),ti+1])
    fluctHetero[ti+1] = 1*(KK-1)*mean(h*fffluct[-c(1,KK+1),ti+1])
    freqHetero[ti+1] = 1*(KK-1)*mean(h*fffreq[-c(1,KK+1),ti+1])
    inbHetero[ti+1] = 1*(KK-1)*mean(h*ffinb[-c(1,KK+1),ti+1])
    mutaccHetero[ti+1] = 1*(KK-1)*mean(h*ffmutacc[-c(1,KK+1),ti+1])
    
    #-------------- Loci Fitness for each category ---------------------
    if(HAloci>0){
      WWhet[ti+1] = (h/2)*ffhet[1,ti+1]*(Wwhet(x = xx[1], Sa= hetsa, SA = hetsA)) + (h/2)*ffhet[KK+1,ti+1]*(Wwhet(x=xx[KK+1], Sa= hetsa, SA = hetsA)) + sum(h*ffhet[-c(1,KK+1),ti+1]*(Wwhet(x=xx[2:KK], Sa= hetsa, SA = hetsA)))   #fitness from het. adva
    }else{
      WWhet[ti+1] = 1
    }
    
    if(IDloci>0){
      WWinb[ti+1] = (h/2)*ffinb[1,ti+1]*(Wwinb(x = xx[1], Sa = inbsa, hh = inbhh)) + (h/2)*ffinb[KK+1,ti+1]*(Wwinb(x=xx[KK+1], Sa = inbsa, hh = inbhh)) + sum(h*ffinb[-c(1,KK+1),ti+1]*(Wwinb(x=xx[2:KK], Sa = inbsa, hh = inbhh)))   #fitness from inbreeding
    }else{
      WWinb[ti+1] = 1
    }
    
    if(FSloci>0){
      WWfluct[ti+1] = (h/2)*fffluct[1,ti+1]*(Wwfluct(x=xx[1], Sa = fluctsa[ti+1], SA = fluctsA[ti+1], hh = flucthh[ti+1])) + (h/2)*fffluct[KK+1,ti+1]*(Wwfluct(x=xx[KK+1], Sa = fluctsa[ti+1], SA = fluctsA[ti+1], hh = flucthh[ti+1])) + sum(h*fffluct[-c(1,KK+1),ti+1]*(Wwfluct(x=xx[2:KK], Sa = fluctsa[ti+1], SA = fluctsA[ti+1], hh = flucthh[ti+1])))  #fitness from fluct. sel with rev. of dom.
    }else{
      WWfluct[ti+1] = 1
    }
    
    if(FDloci>0){
      WWfreq[ti+1] = (h/2)*fffreq[1,ti+1]*(Wwfreq(x=xx[1], Sa= freqsa, SA= freqsA)) + (h/2)*fffreq[KK+1,ti+1]*(Wwfreq(x=xx[KK+1], Sa= freqsa, SA= freqsA)) + sum(h*fffreq[-c(1,KK+1),ti+1]*(Wwfreq(x=xx[2:KK], Sa= freqsa, SA= freqsA)))  #fitness from neg. freq. dep. sel.
    }else{
      WWfreq[ti+1] = 1
    }
    
    if(MAloci>0){
      WWmutacc[ti+1] = (h/2)*ffmutacc[1,ti+1]*(Wwmutacc(x = xx[1], Sa = mutaccsa, hh = mutacchh)) + (h/2)*ffmutacc[KK+1,ti+1]*(Wwmutacc(x=xx[KK+1], Sa = mutaccsa, hh = mutacchh)) + sum(h*ffmutacc[-c(1,KK+1),ti+1]*(Wwmutacc(x=xx[2:KK], Sa = mutaccsa, hh = mutacchh)))   #fitness from mutation accumulation
    }else{
      WWmutacc[ti+1] = 1
    }
    
    # Offspring viability via multiplicative fitness across loci at time t.
    WWt = WWhet[ti+1]^HAloci * WWinb[ti+1]^IDloci * WWfluct[ti+1]^FSloci * WWfreq[ti+1]^FDloci * WWmutacc[ti+1]^MAloci
    WW = round(WWt,5)
    
    Fitness[ti+1] = WW
    
    #------------------------Environmental stochasticity------------------------------------------
    
    envstoch = ENVSTOCHFUNC(gentime = ti, InitStoch = InitStoch, envmax = envmax, envrate = envrate)
    if (CapaStoch == 1){
      
      Kapacity = Capacity*(1/envstoch) + rnorm(1,0,sd = 10*envstoch)
    }else{
      Kapacity = Capacity
    }
    
    if(GrowthStoch == 1){
      RRat = exp(Rate)*(1/envstoch) + rnorm(1,0,sd = 0.01*envstoch)
      RRate = log(RRat)
    }else{
      RRate = Rate
    }
    
    # Hard carrying capacity
      Ne[ti+2] = Ne[ti+1]*min(Fitness[ti+1]*exp(RRate), Kapacity/Ne[ti+1])
    
    
  }
  #Here, we return Population size ,overall  population viablity, and  respecitive viabilities from different mechanisms.
 OUT = cbind(POPSIZE=Ne[-(length(tt)+1)], POPFIT=Fitness, hetHet = hetHetero, fluctHet = fluctHetero, freqHet = freqHetero, inbHet = inbHetero, mutaccHet = mutaccHetero, hetWW = WWhet^HAloci, fluctWW = WWfluct^FSloci, freqWW = WWfreq^FDloci, inbWW = WWinb^IDloci, mutaccWW = WWmutacc^MAloci)
  
  #OUT = cbind(Ne[-(length(tt)+1)], Fitness)
  return(OUT)
  #return(Ne[length(tt)])
}
```
