## Supplementary material for "Modeling minimum viable population size with multiple genetic problems of small populations": R codes used to simulate results: MainFunct_IBM.R

```
# Main function for the Individual- based model (IBM) 
# Environmental stochasticity. Do not change anything, it is not used in this study 
ENVSTOCHFUNC =  function(gentime, InitStoch=1, envmax=1, envrate=0){
  
  Stoch = envmax*InitStoch/(InitStoch + (envmax-InitStoch)*exp(-envrate*(gentime-1))) 
  
  return(Stoch)
}


FITNESS_ALLOCATION<-function( allalleles_mat, HAfitness, FSfitness, FDfitness, IDfitness, MAfitness, numberofloci, LociRatioVector){
  if(nrow(allalleles_mat)>0){
    
    refma<-cbind(rep(1:nrow(allalleles_mat),each=numberofloci),seq(1,numberofloci*2,2))#do before function so we do not repeat uneccessary!
    refmb<-cbind(rep(1:nrow(allalleles_mat),each=numberofloci),seq(2,numberofloci*2,2))
    #setting up locus descriptions
    ma<-matrix(paste(allalleles_mat[refma],allalleles_mat[refmb],sep=""), ncol=numberofloci, nrow = nrow(allalleles_mat), byrow = T) #pasting together so we have AA, aa, Aa, and aA ||| rows are individualsalleles, collumn are 
    
    #calculating fitness
    ma1<-t(apply(ma,1,function(x){rbind(sum(x=="AA"),sum(x=="aa"),sum(x=="Aa"|x=="aA"))})) # transfering it to the number of aa, AA, and aA/Aa Loci ||| collumns are individuals row1==AA,row2==aa,row3==aA/Aa
    colnames(ma1)<-c("AA","aa","Aa/aA")
    #finding polymorphism at population level (polymorphism== which loci are still polymorphic, ma2==how polymorphic are the loci)
    #####polymorphicloci<-numberofloci-sum(apply(ma1==0,1,function(x){sum(x==c(F,T,T))==3|sum(x==c(T,F,T))==3}))# number of poly loci at population level #TRUE means non polymorphic!!
    # propor_aa = sum(ma1[,"aa"])/sum(ma1)
    propor_AA = sum(ma1[,"AA"])/sum(ma1)
    propor_Aa = sum(ma1[,"Aa/aA"])/sum(ma1)
    #pop_freq_a<-(sum(ma1[,2])*2+sum(ma1[,3])) /(sum(ma1)*2) #proportion of loci being a
    pop_freq_A<-(sum(ma1[,1])*2+sum(ma1[,3])) /(sum(ma1)*2) #proportion of loci being A
    
    
    ma2 <- apply(ma,2,function(x){rbind(sum(x=="AA"),sum(x=="aa"),sum(x=="Aa"|x=="aA"))})
    rownames(ma2)<-c("AA","aa","Aa/aA")
    degreeloci_freq_a = (ma2[2,]*2+ma2[3,]) /(colSums(ma2)*2)
    degreeloci_freq_A = (ma2[1,]*2+ma2[3,]) /(colSums(ma2)*2)
    popsize = nrow(allalleles_mat)
    
    
    wa = which(degreeloci_freq_a>0); wA = which(degreeloci_freq_A>0); WW = c(wa,wA)
    polymorphicloci = length(WW)-length(unique(WW))#
    
    #--------------Fitness per selection mechanism-----------------------------------
    #------------------------ Loci ratios ------------
    LociDistr = numberofloci*LociRatioVector
    HAloci = LociDistr[1]
    FSloci = LociDistr[2]
    FDloci = LociDistr[3]
    IDloci = LociDistr[4]
    MAloci = LociDistr[5]
    #-------------------------------
    #For HA
    if(HAloci > 0){
      HAlocimat = allalleles_mat[,1:(2*HAloci), drop = FALSE]
      
      refma<-cbind(rep(1:nrow(HAlocimat),each=HAloci),seq(1,HAloci*2,2))#do before function so we do not repeat uneccessary!
      refmb<-cbind(rep(1:nrow(HAlocimat),each=HAloci),seq(2,HAloci*2,2))
      #setting up locus descriptions
      ma<-matrix(paste(HAlocimat[refma],HAlocimat[refmb],sep=""), ncol=HAloci, nrow = nrow(HAlocimat), byrow = TRUE) #pasting together so we have AA, aa, Aa, and aA ||| rows are individualsalleles, collumn are 
      
      #calculating fitness
      ma1<-t(apply(ma,1,function(x){rbind(sum(x=="AA"),sum(x=="aa"),sum(x=="Aa"|x=="aA"))})) # transfering it to the number of aa, AA, and aA/Aa Loci ||| collumns are individuals row1==AA,row2==aa,row3==aA/Aa
      colnames(ma1)<-c("AA","aa","Aa/aA")
      
      HAindifit<-apply(t(HAfitness^t(ma1)),1,prod) #transfering to fitness (multiplicative) per inidiviudal
    }else{
      HAindifit = 1
    }
    
    # For FS
    if(FSloci > 0){ 
      FSlocimat = allalleles_mat[,(2*HAloci+1):(2*HAloci+2*FSloci), drop = FALSE]
      refma<-cbind(rep(1:nrow(FSlocimat),each=FSloci),seq(1,FSloci*2,2))#do before function so we do not repeat uneccessary!
      refmb<-cbind(rep(1:nrow(FSlocimat),each=FSloci),seq(2,FSloci*2,2))
      #setting up locus descriptions
      ma<-matrix(paste(FSlocimat[refma],FSlocimat[refmb],sep=""), ncol=FSloci, nrow = nrow(FSlocimat), byrow = TRUE) #pasting together so we have AA, aa, Aa, and aA ||| rows are individualsalleles, collumn are 
      
      #calculating fitness
      ma1<-t(apply(ma,1,function(x){rbind(sum(x=="AA"),sum(x=="aa"),sum(x=="Aa"|x=="aA"))})) # transfering it to the number of aa, AA, and aA/Aa Loci ||| collumns are individuals row1==AA,row2==aa,row3==aA/Aa
      colnames(ma1)<-c("AA","aa","Aa/aA")
      
      FSindifit <- apply(t(FSfitness^t(ma1)),1,prod) #transfering to fitness (multiplicative) per inidiviudal
    }else{
      FSindifit = 1
    }
    
    # For FD
    if(FDloci > 0){
      FDlocimat = allalleles_mat[,(2*HAloci+2*FSloci+1):(2*HAloci+2*FSloci+2*FDloci), drop = FALSE]
      refma<-cbind(rep(1:nrow(FDlocimat),each=FDloci),seq(1,FDloci*2,2))#do before function so we do not repeat uneccessary!
      refmb<-cbind(rep(1:nrow(FDlocimat),each=FDloci),seq(2,FDloci*2,2))
      #setting up locus descriptions
      ma<-matrix(paste(FDlocimat[refma],FDlocimat[refmb],sep=""), ncol=FDloci, nrow = nrow(FDlocimat), byrow = TRUE) #pasting together so we have AA, aa, Aa, and aA ||| rows are individualsalleles, collumn are 
      
      #calculating fitness
      #ma1<-t(apply(ma,1,function(x){rbind(sum(x=="AA"),sum(x=="aa"),sum(x=="Aa"|x=="aA"))})) # transfering it to the number of aa, AA, and aA/Aa Loci ||| collumns are individuals row1==AA,row2==aa,row3==aA/Aa
      #colnames(ma1)<-c("AA","aa","Aa/aA")
      
      FDindifit <-  apply(ma, 1, FUN = function(x){
        
        y_AA = which(x == "AA")
        if (length(y_AA)>0){
          yA = FDfitness[1,y_AA] 
        }else{
          yA = NA
        }
        
        y_aa = which(x == "aa")
        if (length(y_aa)>0){
          ya = FDfitness[2,y_aa]
        }else{
          ya = NA
        }
        
        y_Aa = which(x == "Aa" | x == "aA")
        
        if (length(y_Aa)>0){
          yh = FDfitness[3,y_Aa] 
        }else{
          yh = NA
        }
        
        personfit = prod(yA,ya,yh, na.rm = TRUE) 
        return(personfit)
      }) #transfering to fitness (multiplicative) per inidiviudal
    }else{
      FDindifit = 1
    }
    
    # For ID
    if(IDloci > 0){
      IDlocimat = allalleles_mat[,(2*HAloci+2*FSloci+2*FDloci+1):(2*HAloci+2*FSloci+2*FDloci+2*IDloci), drop = FALSE]
      refma<-cbind(rep(1:nrow(IDlocimat),each=IDloci),seq(1,IDloci*2,2))#do before function so we do not repeat uneccessary!
      refmb<-cbind(rep(1:nrow(IDlocimat),each=IDloci),seq(2,IDloci*2,2))
      #setting up locus descriptions
      ma<-matrix(paste(IDlocimat[refma],IDlocimat[refmb],sep=""), ncol=IDloci, nrow = nrow(IDlocimat), byrow = TRUE) #pasting together so we have AA, aa, Aa, and aA ||| rows are individualsalleles, collumn are 
      
      #calculating fitness
      ma1<-t(apply(ma,1,function(x){rbind(sum(x=="AA"),sum(x=="aa"),sum(x=="Aa"|x=="aA"))})) # transfering it to the number of aa, AA, and aA/Aa Loci ||| collumns are individuals row1==AA,row2==aa,row3==aA/Aa
      colnames(ma1)<-c("AA","aa","Aa/aA")
      
      IDindifit<-apply(t(IDfitness^t(ma1)),1,prod) #transfering to fitness (multiplicative) per inidiviudal
    }else{
      IDindifit = 1
    }
    
    # For MA
    if(MAloci > 0){
      MAlocimat = allalleles_mat[,(2*HAloci+2*FSloci+2*FDloci+2*IDloci+1):(2*HAloci+2*FSloci+2*FDloci+2*IDloci+2*MAloci), drop = FALSE]
      refma<-cbind(rep(1:nrow(MAlocimat),each=MAloci),seq(1,MAloci*2,2))#do before function so we do not repeat uneccessary!
      refmb<-cbind(rep(1:nrow(MAlocimat),each=MAloci),seq(2,MAloci*2,2))
      #setting up locus descriptions
      ma<-matrix(paste(MAlocimat[refma],MAlocimat[refmb],sep=""), ncol=MAloci, nrow = nrow(MAlocimat), byrow = TRUE) #pasting together so we have AA, aa, Aa, and aA ||| rows are individualsalleles, collumn are 
      
      #calculating fitness
      ma1<-t(apply(ma,1,function(x){rbind(sum(x=="AA"),sum(x=="aa"),sum(x=="Aa"|x=="aA"))})) # transfering it to the number of aa, AA, and aA/Aa Loci ||| collumns are individuals row1==AA,row2==aa,row3==aA/Aa
      colnames(ma1)<-c("AA","aa","Aa/aA")
      
      MAindifit<-apply(t(MAfitness^t(ma1)),1,prod) #transfering to fitness (multiplicative) per inidiviudal
    }else{
      MAindifit = 1
    }
    
    # Offspring viability via multiplicative fitness across loci
    indifit = HAindifit * FSindifit * FDindifit * IDindifit * MAindifit
    
    popfit <- mean(indifit)
    
    #returns pop. size, pop. mean viability fitness, proportions of Aa and AA,allele frequency of A at population and locus level respectively.
    return(list(popsize, popfit, polymorphicloci, propor_Aa, propor_AA, pop_freq_A, degreeloci_freq_A))
  }else{
    popfit = 0
    polymorphicloci = 0
    propor_Aa = 0
    propor_AA = 0
    pop_freq_A = 0
    degreeloci_freq_A = rep(0, numberofloci)
    popsize = 0
    return(list(popsize, popfit, polymorphicloci, propor_Aa, propor_AA, pop_freq_A, degreeloci_freq_A))
    
  }}


# This function gives us the genes for each child at each locus
BABYMAT = function(allalleles_mat, numberofloci, LociRatioVector, HAmut_a, HAmut_A, FSmut_a, FSmut_A, FDmut_a, FDmut_A, IDmut_a, IDmut_A, MAmut_a, MAmut_A, deathrate, Rate, Capacity, envstoch=1, CapaStoch=0,GrowthStoch=0){
  
  #------------------------ Loci ratios ------------------------
  LociDistr = numberofloci*LociRatioVector
  HAloci = LociDistr[1]
  FSloci = LociDistr[2]
  FDloci = LociDistr[3]
  IDloci = LociDistr[4]
  MAloci = LociDistr[5]
  #-------------------------------------------------------------
  
  N = nrow(allalleles_mat)
  
  if (CapaStoch == 1){
    Capacity = rnorm(1,Capacity, sd = envstoch)
  }
  
  if(GrowthStoch == 1){
    Rate = exp(Rate)
    Rate = rnorm(1,Rate, sd = envstoch/100)
    Rate = log(Rate)
  }
  
  reproducers = 1-rbinom(N,1,deathrate)
  tot_reproducers = sum(reproducers)
  
  
    tot_offspring = rpois(1,tot_reproducers*exp(Rate))
  
  if(tot_offspring>0){
    mother_index = sample(tot_reproducers,tot_offspring,replace = TRUE)#1:tot_reproducers
    father_index = sample(tot_reproducers,tot_offspring,replace = TRUE)
    
    #-----------------------------------HA-------------------------------------------
    if(HAloci > 0){
    HAallalleles_reproducer = allalleles_mat[(reproducers==1), 1:(2*HAloci), drop = FALSE]
    fathergenes = HAallalleles_reproducer[father_index,,drop = FALSE]
    mothergenes = HAallalleles_reproducer[mother_index,,drop = FALSE]
    
    genesmother<-matrix(c(mothergenes[,seq(1,ncol(mothergenes),2)],mothergenes[,seq(2,ncol(mothergenes),2)]),ncol = 2, byrow = FALSE)
    genesfather<-matrix(c(fathergenes[,seq(1,ncol(fathergenes),2)],fathergenes[,seq(2,ncol(fathergenes),2)]), ncol = 2, byrow = FALSE)
    
    
    frompa = apply(genesfather, 1, function(x) {sample(x,1,replace = TRUE)})
    fromma = apply(genesmother, 1, function(x) {sample(x,1,replace = TRUE)})
    HAgenesbaby = matrix(0, nrow = tot_offspring, ncol = 2*HAloci)
    xx = seq(1,ncol(HAgenesbaby), 2)
    yy = seq(2, ncol(HAgenesbaby),2)
    HAgenesbaby[,xx] = fromma
    HAgenesbaby[,yy] = frompa 
    index_a = which(HAgenesbaby=="a")
    index_A = which(HAgenesbaby=="A")
    
    HAMUTS_a = which(runif(length(which(HAgenesbaby=="a")))<=HAmut_a) #selecting places where mutation happens for a
    HAMUTS_A = which(runif(length(which(HAgenesbaby=="A")))<=HAmut_A) #selecting places where mutation happens for A
    if (length(HAMUTS_a)>0){
      HAgenesbaby[index_a[HAMUTS_a]] = "A"
    }
    if (length(HAMUTS_A)>0){
      HAgenesbaby[index_A[HAMUTS_A]] = "a"
    }
    
    }else{
      HAgenesbaby = matrix(0, nrow = tot_offspring, ncol = 0)
    }
    
    
    #-------------------------------------------------------------------------------------
    
    #----------------------------------FS--------------------------------------------
    if(FSloci > 0){
      FSallalleles_reproducer = allalleles_mat[(reproducers==1), (2*HAloci+1):(2*HAloci+2*FSloci), drop = FALSE]
      fathergenes = FSallalleles_reproducer[father_index,,drop = FALSE]
      mothergenes = FSallalleles_reproducer[mother_index,,drop = FALSE]
      
      genesmother<-matrix(c(mothergenes[,seq(1,ncol(mothergenes),2)],mothergenes[,seq(2,ncol(mothergenes),2)]),ncol = 2, byrow = FALSE)
      genesfather<-matrix(c(fathergenes[,seq(1,ncol(fathergenes),2)],fathergenes[,seq(2,ncol(fathergenes),2)]), ncol = 2, byrow = FALSE)
      
      
      frompa = apply(genesfather, 1, function(x) {sample(x,1,replace = TRUE)})
      fromma = apply(genesmother, 1, function(x) {sample(x,1,replace = TRUE)})
      FSgenesbaby = matrix(0, nrow = tot_offspring, ncol = 2*FSloci)
      xx = seq(1,ncol(FSgenesbaby), 2)
      yy = seq(2, ncol(FSgenesbaby),2)
      FSgenesbaby[,xx] = fromma
      FSgenesbaby[,yy] = frompa 
      index_a = which(FSgenesbaby=="a")
      index_A = which(FSgenesbaby=="A")
      
      FSMUTS_a = which(runif(length(which(FSgenesbaby=="a")))<=FSmut_a) #selecting places where mutation happens for a
      FSMUTS_A = which(runif(length(which(FSgenesbaby=="A")))<=FSmut_A) #selecting places where mutation happens for A
      if (length(FSMUTS_a)>0){
        FSgenesbaby[index_a[FSMUTS_a]] = "A"
      }
      if (length(FSMUTS_A)>0){
        FSgenesbaby[index_A[FSMUTS_A]] = "a"
      }
      
    }else{
      FSgenesbaby = matrix(0, nrow = tot_offspring, ncol = 0)
    }
    
    
    #-------------------------------------------------------------------------------------
    
    #----------------------------------FD--------------------------------------------
    if(FDloci > 0){
      FDallalleles_reproducer = allalleles_mat[(reproducers==1), (2*HAloci+2*FSloci+1):(2*HAloci+2*FSloci+2*FDloci), drop = FALSE]
     # print(1*FDloci)
      fathergenes = FDallalleles_reproducer[father_index,,drop = FALSE]
      mothergenes = FDallalleles_reproducer[mother_index,,drop = FALSE]
      
      genesmother<-matrix(c(mothergenes[,seq(1,ncol(mothergenes),2)],mothergenes[,seq(2,ncol(mothergenes),2)]),ncol = 2, byrow = FALSE)
      genesfather<-matrix(c(fathergenes[,seq(1,ncol(fathergenes),2)],fathergenes[,seq(2,ncol(fathergenes),2)]), ncol = 2, byrow = FALSE)
      
      
      frompa = apply(genesfather, 1, function(x) {sample(x,1,replace = TRUE)})
      fromma = apply(genesmother, 1, function(x) {sample(x,1,replace = TRUE)})
      FDgenesbaby = matrix(0, nrow = tot_offspring, ncol = (2*FDloci))
      xx = seq(1,ncol(FDgenesbaby), 2)
      yy = seq(2, ncol(FDgenesbaby),2)
      #print(length(xx))
      #print(ncol(FDgenesbaby))
      FDgenesbaby[,xx] = fromma
      FDgenesbaby[,yy] = frompa 
      index_a = which(FDgenesbaby=="a")
      index_A = which(FDgenesbaby=="A")
      
      FDMUTS_a = which(runif(length(which(FDgenesbaby=="a")))<=FDmut_a) #selecting places where mutation happens for a
      FDMUTS_A = which(runif(length(which(FDgenesbaby=="A")))<=FDmut_A) #selecting places where mutation happens for A
      if (length(FDMUTS_a)>0){
        FDgenesbaby[index_a[FDMUTS_a]] = "A"
      }
      if (length(FDMUTS_A)>0){
        FDgenesbaby[index_A[FDMUTS_A]] = "a"
      }
      
    }else{
      FDgenesbaby = matrix(0, nrow = tot_offspring, ncol = 0)
    }
    
    
    #-------------------------------------------------------------------------------------
    #---------------------------------- ID --------------------------------------------
    if(IDloci > 0){
      IDallalleles_reproducer = allalleles_mat[(reproducers==1), (2*HAloci+2*FSloci+2*FDloci+1):(2*HAloci+2*FSloci+2*FDloci+2*IDloci), drop = FALSE]
      fathergenes = IDallalleles_reproducer[father_index,,drop = FALSE]
      mothergenes = IDallalleles_reproducer[mother_index,,drop = FALSE]
      
      genesmother<-matrix(c(mothergenes[,seq(1,ncol(mothergenes),2)],mothergenes[,seq(2,ncol(mothergenes),2)]),ncol = 2, byrow = FALSE)
      genesfather<-matrix(c(fathergenes[,seq(1,ncol(fathergenes),2)],fathergenes[,seq(2,ncol(fathergenes),2)]), ncol = 2, byrow = FALSE)
      
      
      frompa = apply(genesfather, 1, function(x) {sample(x,1,replace = TRUE)})
      fromma = apply(genesmother, 1, function(x) {sample(x,1,replace = TRUE)})
      IDgenesbaby = matrix(0, nrow = tot_offspring, ncol = 2*IDloci)
      xx = seq(1,ncol(IDgenesbaby), 2)
      yy = seq(2, ncol(IDgenesbaby),2)
      IDgenesbaby[,xx] = fromma
      IDgenesbaby[,yy] = frompa 
      index_a = which(IDgenesbaby=="a")
      index_A = which(IDgenesbaby=="A")
      
      IDMUTS_a = which(runif(length(which(IDgenesbaby=="a")))<=IDmut_a) #selecting places where mutation happens for a
      IDMUTS_A = which(runif(length(which(IDgenesbaby=="A")))<=IDmut_A) #selecting places where mutation happens for A
      if (length(IDMUTS_a)>0){
        IDgenesbaby[index_a[IDMUTS_a]] = "A"
      }
      if (length(IDMUTS_A)>0){
        IDgenesbaby[index_A[IDMUTS_A]] = "a"
      }
      
    }else{
      IDgenesbaby = matrix(0, nrow = tot_offspring, ncol = 0)
    }
    
    
    #-------------------------------------------------------------------------------------
    #---------------------------------- MA --------------------------------------------
    if(MAloci > 0){
      MAallalleles_reproducer = allalleles_mat[(reproducers==1), (2*HAloci+2*FSloci+2*FDloci+2*IDloci+1):(2*HAloci+2*FSloci+2*FDloci+2*IDloci+2*MAloci), drop = FALSE]
      fathergenes = MAallalleles_reproducer[father_index,,drop = FALSE]
      mothergenes = MAallalleles_reproducer[mother_index,,drop = FALSE]
      
      genesmother<-matrix(c(mothergenes[,seq(1,ncol(mothergenes),2)],mothergenes[,seq(2,ncol(mothergenes),2)]),ncol = 2, byrow = FALSE)
      genesfather<-matrix(c(fathergenes[,seq(1,ncol(fathergenes),2)],fathergenes[,seq(2,ncol(fathergenes),2)]), ncol = 2, byrow = FALSE)
      
      
      frompa = apply(genesfather, 1, function(x) {sample(x,1,replace = TRUE)})
      fromma = apply(genesmother, 1, function(x) {sample(x,1,replace = TRUE)})
      MAgenesbaby = matrix(0, nrow = tot_offspring, ncol = 2*MAloci)
      xx = seq(1,ncol(MAgenesbaby), 2)
      yy = seq(2, ncol(MAgenesbaby),2)
      MAgenesbaby[,xx] = fromma
      MAgenesbaby[,yy] = frompa 
      index_a = which(MAgenesbaby=="a")
      index_A = which(MAgenesbaby=="A")
      
      MAMUTS_a = which(runif(length(which(MAgenesbaby=="a")))<=MAmut_a) #selecting places where mutation happens for a
      MAMUTS_A = which(runif(length(which(MAgenesbaby=="A")))<=MAmut_A) #selecting places where mutation happens for A
      if (length(MAMUTS_a)>0){
        MAgenesbaby[index_a[MAMUTS_a]] = "A"
      }
      if (length(MAMUTS_A)>0){
        MAgenesbaby[index_A[MAMUTS_A]] = "a"
      }
      
    }else{
      MAgenesbaby = matrix(0, nrow = tot_offspring, ncol = 0)
    }
    
    
    #-------------------------------------------------------------------------------------
    genesbaby = cbind(HAgenesbaby,FSgenesbaby,FDgenesbaby,IDgenesbaby,MAgenesbaby)
    
    
    return(genesbaby)
  }
}


#This function gives offspring viability for each offspring
OFFSPRING_FITNESS <- function( allalleles_mat, HAfitness, FSfitness, FDfitness, IDfitness, MAfitness, numberofloci, LociRatioVector){
  if(nrow(allalleles_mat)>0){
    
    #------------------------ Loci ratios
    LociDistr = numberofloci*LociRatioVector
    HAloci = LociDistr[1]
    FSloci = LociDistr[2]
    FDloci = LociDistr[3]
    IDloci = LociDistr[4]
    MAloci = LociDistr[5]
    #-------------------------------
    
    if(HAloci > 0){
      HAlocimat = allalleles_mat[,1:(2*HAloci), drop = FALSE]
      refma<-cbind(rep(1:nrow(HAlocimat),each=HAloci),seq(1,HAloci*2,2))#do before function so we do not repeat uneccessary!
      refmb<-cbind(rep(1:nrow(HAlocimat),each=HAloci),seq(2,HAloci*2,2))
      #setting up locus descriptions
      ma<-matrix(paste(HAlocimat[refma],HAlocimat[refmb],sep=""), ncol=HAloci, nrow = nrow(HAlocimat), byrow = TRUE) #pasting together so we have AA, aa, Aa, and aA ||| rows are individualsalleles, collumn are 
      
      #calculating fitness
      ma1<-t(apply(ma,1,function(x){rbind(sum(x=="AA"),sum(x=="aa"),sum(x=="Aa"|x=="aA"))})) # transfering it to the number of aa, AA, and aA/Aa Loci ||| collumns are individuals row1==AA,row2==aa,row3==aA/Aa
      colnames(ma1)<-c("AA","aa","Aa/aA")
      
      HAindifit<-apply(t(HAfitness^t(ma1)),1,prod) #transfering to fitness (multiplicative) per inidiviudal
    }else{
      HAindifit = 1
    }
    
    
    if(FSloci > 0){ 
      FSlocimat = allalleles_mat[,(2*HAloci+1):(2*HAloci+2*FSloci), drop = FALSE]
      refma<-cbind(rep(1:nrow(FSlocimat),each=FSloci),seq(1,FSloci*2,2))#do before function so we do not repeat uneccessary!
      refmb<-cbind(rep(1:nrow(FSlocimat),each=FSloci),seq(2,FSloci*2,2))
      #setting up locus descriptions
      ma<-matrix(paste(FSlocimat[refma],FSlocimat[refmb],sep=""), ncol=FSloci, nrow = nrow(FSlocimat), byrow = TRUE) #pasting together so we have AA, aa, Aa, and aA ||| rows are individualsalleles, collumn are 
      
      #calculating fitness
      ma1<-t(apply(ma,1,function(x){rbind(sum(x=="AA"),sum(x=="aa"),sum(x=="Aa"|x=="aA"))})) # transfering it to the number of aa, AA, and aA/Aa Loci ||| collumns are individuals row1==AA,row2==aa,row3==aA/Aa
      colnames(ma1)<-c("AA","aa","Aa/aA")
      
      FSindifit <- apply(t(FSfitness^t(ma1)),1,prod) #transfering to fitness (multiplicative) per inidiviudal
    }else{
      FSindifit = 1
    }
    
    
    if(FDloci > 0){
      FDlocimat = allalleles_mat[,(2*HAloci+2*FSloci+1):(2*HAloci+2*FSloci+2*FDloci), drop = FALSE]
      refma<-cbind(rep(1:nrow(FDlocimat),each=FDloci),seq(1,FDloci*2,2))#do before function so we do not repeat uneccessary!
      refmb<-cbind(rep(1:nrow(FDlocimat),each=FDloci),seq(2,FDloci*2,2))
      #setting up locus descriptions
      ma<-matrix(paste(FDlocimat[refma],FDlocimat[refmb],sep=""), ncol=FDloci, nrow = nrow(FDlocimat), byrow = TRUE) #pasting together so we have AA, aa, Aa, and aA ||| rows are individualsalleles, collumn are 
      
      #calculating fitness
      #ma1<-t(apply(ma,1,function(x){rbind(sum(x=="AA"),sum(x=="aa"),sum(x=="Aa"|x=="aA"))})) # transfering it to the number of aa, AA, and aA/Aa Loci ||| collumns are individuals row1==AA,row2==aa,row3==aA/Aa
      #colnames(ma1)<-c("AA","aa","Aa/aA")
      
      FDindifit <-  apply(ma, 1, FUN = function(x){
        
        y_AA = which(x == "AA")
        if (length(y_AA)>0){
          yA = FDfitness[1,y_AA] 
        }else{
          yA = NA
        }
        
        y_aa = which(x == "aa")
        if (length(y_aa)>0){
          ya = FDfitness[2,y_aa]
        }else{
          ya = NA
        }
        
        y_Aa = which(x == "Aa" | x == "aA")
        
        if (length(y_Aa)>0){
          yh = FDfitness[3,y_Aa] 
        }else{
          yh = NA
        }
        
        personfit = prod(yA,ya,yh, na.rm = TRUE) 
        return(personfit)
      }) #transfering to fitness (multiplicative) per inidiviudal
    }else{
      FDindifit = 1
    }
    
    
    if(IDloci > 0){
      IDlocimat = allalleles_mat[,(2*HAloci+2*FSloci+2*FDloci+1):(2*HAloci+2*FSloci+2*FDloci+2*IDloci), drop = FALSE]
      refma<-cbind(rep(1:nrow(IDlocimat),each=IDloci),seq(1,IDloci*2,2))#do before function so we do not repeat uneccessary!
      refmb<-cbind(rep(1:nrow(IDlocimat),each=IDloci),seq(2,IDloci*2,2))
      #setting up locus descriptions
      ma<-matrix(paste(IDlocimat[refma],IDlocimat[refmb],sep=""), ncol=IDloci, nrow = nrow(IDlocimat), byrow = TRUE) #pasting together so we have AA, aa, Aa, and aA ||| rows are individualsalleles, collumn are 
      
      #calculating fitness
      ma1<-t(apply(ma,1,function(x){rbind(sum(x=="AA"),sum(x=="aa"),sum(x=="Aa"|x=="aA"))})) # transfering it to the number of aa, AA, and aA/Aa Loci ||| collumns are individuals row1==AA,row2==aa,row3==aA/Aa
      colnames(ma1)<-c("AA","aa","Aa/aA")
      
      IDindifit<-apply(t(IDfitness^t(ma1)),1,prod) #transfering to fitness (multiplicative) per inidiviudal
    }else{
      IDindifit = 1
    }
    
    
    if(MAloci > 0){
      MAlocimat = allalleles_mat[,(2*HAloci+2*FSloci+2*FDloci+2*IDloci+1):(2*HAloci+2*FSloci+2*FDloci+2*IDloci+2*MAloci), drop = FALSE]
      refma<-cbind(rep(1:nrow(MAlocimat),each=MAloci),seq(1,MAloci*2,2))#do before function so we do not repeat uneccessary!
      refmb<-cbind(rep(1:nrow(MAlocimat),each=MAloci),seq(2,MAloci*2,2))
      #setting up locus descriptions
      ma<-matrix(paste(MAlocimat[refma],MAlocimat[refmb],sep=""), ncol=MAloci, nrow = nrow(MAlocimat), byrow = TRUE) #pasting together so we have AA, aa, Aa, and aA ||| rows are individualsalleles, collumn are 
      
      #calculating fitness
      ma1<-t(apply(ma,1,function(x){rbind(sum(x=="AA"),sum(x=="aa"),sum(x=="Aa"|x=="aA"))})) # transfering it to the number of aa, AA, and aA/Aa Loci ||| collumns are individuals row1==AA,row2==aa,row3==aA/Aa
      colnames(ma1)<-c("AA","aa","Aa/aA")
      
      MAindifit<-apply(t(MAfitness^t(ma1)),1,prod) #transfering to fitness (multiplicative) per inidiviudal
    }else{
      MAindifit = 1
    }
    
    
    indifit = HAindifit * FSindifit * FDindifit* IDindifit* MAindifit
    #print(mean(indifit))
    return(indifit)
  }else{
    return(-1)
  }
  
}


#------------ Main function that combines the above functions ------------------
Main_IBM <- function( founderpop, numberofloci = 100, Capacity, LociRatioVector, HAmut_a, HAmut_A, FSmut_a, FSmut_A, FDmut_a, FDmut_A, IDmut_a, IDmut_A, MAmut_a , MAmut_A, tmax, x0, hetsa, hetsA, fluctsa, fluctsA, freqsa, freqsA, IDsa, MAsa, ID_h, MA_h = 0,Rate,deathrate = 0, percent, Kapa = 50,InitStoch = 1, envrate = 0, envmax = 0, CapaStoch = 0, GrowthStoch = 0) {
  
  #------------------------ Loci ratios
  LociDistr = numberofloci*LociRatioVector
  HAloci = LociDistr[1]
  FSloci = LociDistr[2]
  FDloci = LociDistr[3]
  IDloci = LociDistr[4]
  MAloci = LociDistr[5]
  #-------------------------------
  
  populationsize <- rep(0,tmax)
  NumberLociPoly = rep(0, tmax)# Number of polymorphic loci
  PopFreq_A = rep(0, tmax)# frequency of "A" in the population
  POPfit = rep(0, tmax)# population fitness
  Proportion_Aa = rep(0, tmax)#proportion of Aa at population level 
  #Proportion_AA = rep(0, tmax)# Proportion of AA at population level
  LociFreq_A = list(matrix(-1, nrow = tmax, ncol = numberofloci))# frequency of allele "A" at locus level
  
  
  noMate = 0
  
  alleles <- c("a", "A") # alleles present at each locus
  
  # Initial fitness of genotypes
  Fit_AA = 1
  Fit_Aa = 1
  Fit_aa = 1
  
  
    if (FDloci>0){
      FDfitness_init = matrix(1,nrow = 3, ncol = FDloci)}else{
        FDfitness = matrix(1,nrow = 3, ncol = 0)
      }
  
    HAfitness_init = c(Fit_AA, Fit_aa, Fit_Aa)
    FSfitness_init = c(Fit_AA, Fit_aa, Fit_Aa)
    IDfitness_init = c(Fit_AA, Fit_aa, Fit_Aa)
    MAfitness_init = c(Fit_AA, Fit_aa, Fit_Aa)
  
  
  inpop = rpois(1,founderpop)# Starting pop
  
  allalleles <- matrix(sample(alleles, inpop * 2* numberofloci,replace=TRUE, prob = c(x0, 1-x0)),ncol=2*numberofloci)#allels to be distributed among individuals
  
  
  # Initial parameters from the initial population 
  Output1 = FITNESS_ALLOCATION(allalleles_mat = allalleles, numberofloci = numberofloci, HAfitness = HAfitness_init, FSfitness = FSfitness_init, FDfitness = FDfitness_init, IDfitness = IDfitness_init, MAfitness = MAfitness_init, LociRatioVector = LociRatioVector)
  populationsize[1] <- Output1[[1]]
  POPfit[1] = Output1[[2]]
  NumberLociPoly[1] =  Output1[[3]]
  Proportion_Aa[1] = Output1[[4]]
  #Proportion_AA[1] = Output1[[5]]
  PopFreq_A[1] = Output1[[6]]
  LociFreq_A[[1]] = Output1[[7]]
  
  
  Ti = 1:tmax
  fluct_s <- fluctsa*sin(2*pi*Ti/Kapa + pi) # selection coeff for genotype AA
  fluct_t <- fluctsA*sin(2*pi*Ti/Kapa)  
  
  fluct_h = 0.5-(percent/100)*sin(2*pi*Ti/Kapa) 
  
  
  FS_AA = (1-fluct_t)/(1+max(fluctsa,fluctsA))
  FS_aa = (1-fluct_s)/(1+max(fluctsa,fluctsA))
  FS_Aa =  (1-fluct_h*fluct_t-(1-fluct_h)*fluct_s)/(1+max(fluctsa,fluctsA)) 
  
  t = 2
  
  # looping through generations as long as population size is above 0
  while (t <= tmax & populationsize[t-1] > 0){
    #fitness of genotypes
    #-----------Het. Adv ---------------------
    
      HFit_AA = 1-hetsA
      HFit_aa = 1-hetsa
      HFit_Aa = 1
      HAfitness = c(HFit_AA, HFit_aa, HFit_Aa)
      
   #---------- Fluct. Selection ---------------
      
      FSfitness = c(FS_AA[t-1], FS_aa[t-1], FS_Aa[t-1])
    
      #--------------Freq. Dep ---------------
      if (FDloci>0){
      Fitmat = matrix(0, nrow = 3, ncol = FDloci)#c(FitAA, Fitaa, FitAa)  
      freqA =  LociFreq_A[[t-1]][(HAloci+FSloci+1):(HAloci+FSloci+FDloci)]#PopFreq_A[t-1]
      #print(freqA)
      FitAA = 1-freqsA*freqA
      Fitaa = 1-freqsa*(1-freqA)
      FitAa = 1-(freqsA*freqA + freqsa*(1-freqA) )/2
      FDfitness = matrix(c(FitAA, Fitaa, FitAa), nrow = 3, ncol = FDloci, byrow = TRUE)#c(FitAA, Fitaa, FitAa)
      }else{
        FDfitness = matrix(1,nrow = 3, ncol = 0)
      }
    #------------Inbreeding Dep.--------------
    
    IFit_AA = 1
    IFit_aa = 1-IDsa
    IFit_Aa = 1 - ID_h*IDsa
    IDfitness = c(IFit_AA, IFit_aa, IFit_Aa)
    
    #------------Mut. Accumulation --------------
    
    MFit_AA = 1
    MFit_aa = 1-MAsa
    MFit_Aa = 1 - MA_h * MAsa
    MAfitness = c(MFit_AA, MFit_aa, MFit_Aa)
    
    #-------------------------------------
    
    
    N = nrow(allalleles)
    
    
    
    if (N > 0){
      #let all the individuals reproduce
      envstoch = ENVSTOCHFUNC(gentime = t-1, InitStoch = InitStoch, envmax = envmax, envrate = envrate)
      offspringmat = BABYMAT(allalleles_mat = allalleles, numberofloci = numberofloci, LociRatioVector = LociRatioVector, HAmut_a = HAmut_a, HAmut_A = HAmut_A, FSmut_a = FSmut_a, FSmut_A = FSmut_A, FDmut_a = FDmut_a, FDmut_A = FDmut_A, IDmut_a = IDmut_a, IDmut_A = IDmut_A, MAmut_a = MAmut_a, MAmut_A = MAmut_A, deathrate = deathrate, Rate = Rate, Capacity = Capacity, envstoch = envstoch, CapaStoch = CapaStoch, GrowthStoch = GrowthStoch)
      
      
      if (is.numeric(nrow(offspringmat))){
        offspringfit = OFFSPRING_FITNESS(allalleles_mat = offspringmat, numberofloci = numberofloci, HAfitness = HAfitness, FSfitness = FSfitness, FDfitness = FDfitness, IDfitness = IDfitness, MAfitness = MAfitness, LociRatioVector = LociRatioVector)
        
        Viable1 = which(runif(nrow(offspringmat))<=offspringfit)
        offspringmat2 = offspringmat[Viable1,,drop = FALSE]
        Accepted =  min(length(Viable1), Capacity)
       
        ss = sample(nrow(offspringmat2), Accepted, replace = FALSE)
        Viable = Viable1[ss]
       # Viable
        
        if (length(ss)>0){
          allalleles = offspringmat2[ss, ,drop =FALSE]

          # updating the measurements where viable offspring replace parents
          
          Output2 = FITNESS_ALLOCATION(allalleles_mat = allalleles, numberofloci = numberofloci, HAfitness = HAfitness, FSfitness = FSfitness, FDfitness = FDfitness, IDfitness = IDfitness, MAfitness = MAfitness, LociRatioVector = LociRatioVector)
          populationsize[t] <- Output2[[1]]
          POPfit[t] = Output2[[2]]
          NumberLociPoly[t] =  Output2[[3]]
          Proportion_Aa[t] = Output2[[4]]
          #Proportion_AA[t] = Output2[[5]]
          PopFreq_A[t] = Output2[[6]]
          LociFreq_A[[t]] = Output2[[7]]
          #print(populationsize[t])
        }else{
          populationsize[t] <- 0
          POPfit[t] = 0
          NumberLociPoly[t] =  0
          Proportion_Aa[t] = 0
          #Proportion_AA[t] = 0
          PopFreq_A[t] = 0
          LociFreq_A[[t]] = rep(0,numberofloci)
        }
      }
      
      t = t + 1
    }
    
  }
  # 
  #OUT = cbind(POPSIZE = populationsize, POLYLOCI = NumberLociPoly, MEANFIT = POPfit, PROPOHET = Proportion_Aa, FREQA = PopFreq_A)
  # Output Population size and mean population viability fitness. 
 OUT = cbind(POPSIZE = populationsize, MEANFIT = POPfit)
  return(OUT)
}
```
