## Supplementary material for "Modeling minimum viable population size with multiple genetic problems of small populations": R codes used to simulate results: MainFunct_MarkovChain_POPequiApproach_FixedLoci.R

```
# Main function for the Markov chain with fixed number of loci
Allefreq = function(p, wAA, wAa, waa, mu, bita){
  wbar  <- p^2 * wAA + 2*p*(1-p)*wAa + (1-p)^2 * waa #assuming Hardy-Weinberg equilibrium
  pselect  <- (p^2 * wAA + p*(1-p)*wAa)/wbar #expected allele frequency after selection
  pmut  <- pselect * (1-mu) + (1-pselect)*bita #expected allele frequency after selection and mutation
  return(pmut)
}

PoisThreshold = 35

Fitness = function(N, wAA, wAa, waa, mu, bita, loci){
  ##Transition matrix, state is number of copies of the A allele and takes values 0,1,2,...,2N.
  TransMat  <- matrix(0,nrow=2*N+1,ncol=2*N+1)
  ##The entry in row j and column k is the probability that the population goes from state j to state k in one generation.
  for (j in 0:(2*N)){
    p  <- j/(2*N)#frequency of allele A before selection
    pmut = Allefreq(p=p, wAA=wAA, wAa=wAa, waa=waa, mu = mu, bita=bita)
    
    ## binomial sampling probabilities
     for (k in 0:(2*N)){
      if (N<=500){
        TransMat[j+1,k+1] <- choose(2*N,k) * pmut^k * (1-pmut)^(2*N-k) # binomial sampling
      }else if(2*N*pmut <PoisThreshold){
          TransMat[j+1,k+1] <- dpois(x=k, lambda = 2*N*pmut) # Poisson distribution approximation
      }else if(2*N*(1-pmut)<PoisThreshold){
        TransMat[j+1,k+1] <- dpois(x=(2*N-k), lambda = 2*N*(1-pmut)) # Poisson distribution approximation
      }else{
          TransMat[j+1,k+1] <- dnorm(x=k, mean = 2*N*pmut, sd = sqrt(2*N*pmut*(1-pmut))) # Normal distribution approximation 
        }
        
      
    }
  }

  
  ## get eigenvalues and eigenvectors
  eigen  <- eigen(t(TransMat))
  #get stationary distribution as eigenvector belonging to the eigenvalue with largest real part (should actually always be 1)
  unnormalizedeq <- Re(eigen$vectors[,which(Re(eigen$values)==max(Re(eigen$values)))])
  normalizedeq  <- unnormalizedeq/(sum(unnormalizedeq))
  
  MeanFitness = function(x){x^2*wAA + 2*x*(1-x)*wAa + (1-x)^2*waa} # Mean fitness assuming infinite population size
  
  FFit = sum(normalizedeq * MeanFitness(x=(0:(2*N))/(2*N)))^loci # Offspring survival fitness using the distribution
 
  
  return(FFit)
}


MainFuncPois = function(numberofloci, LociRatioVector, N_eq, Rate, MAsA = 0, MAsa, MA_h=0, bitaID, bitaMA, IDsa, ID_h=0, FDsA, FDsa, FSsa, FSsA, percent, kapa, HAsA, HAsa, mu, bita){
  
  #------------------------ Loci ratios
  LociDistr = numberofloci*LociRatioVector # Actual number of loci under each selection mechanism
  HAloci = LociDistr[1] # Number of loci under HA
  FSloci = LociDistr[2]# Number of loci under FS
  FDloci = LociDistr[3]# Number of loci under FS
  IDloci = LociDistr[4]# Number of loci under ID
  MAloci = LociDistr[5]# Number of loci under MA
  #-------------------------------
  
  #--------- Heterozygote Advantage (HA)---------------------------------
  if (HAloci > 0){
    # Fitness of homozygote AA, heterozygote Aa and homozygote aa
    HAwAA  <- 1-HAsA # Fitness of homozygote AA
    HAwAa  <- 1 # Fitness of heterozygote Aa
    HAwaa  <- 1-HAsa # Fitness of homozygote aa
    
    # Offspring viability fitness for loci under HA
    HAFit = Fitness(N=N_eq, wAA = HAwAA, wAa = HAwAa, waa = HAwaa, mu = mu, bita = bita, loci = HAloci)
  }else{
    HAFit = 1 
  }
  
  #-------------- Fluctuating Selection with reversal of dominance (FS) ------------------
  if(FSloci > 0){
    Ti = 1:kapa # generations in a complete cycle
    fluct_s <- FSsa*sin(2*pi*Ti/kapa + pi) # selection coefficient for genotype AA
    fluct_t <- FSsA*sin(2*pi*Ti/kapa)  # selection coefficient for aa
    fluct_h = 0.5-(percent/100)*sin(2*pi*Ti/kapa) # dominance coefficients 
    
    #Relative fitness of homozygote AA, heterozygote Aa and homozygote aa
    FS_AA = (1-fluct_t)/(1+max(FSsa,FSsA))
    FS_aa = (1-fluct_s)/(1+max(FSsa,FSsA))
    FS_Aa =  (1-fluct_h*fluct_t-(1-fluct_h)*fluct_s)/(1+max(FSsa,FSsA)) 
    
    # geometric mean relative fitness of homozygote AA, heterozygote Aa and homozygote aa
    FSwAA = exp(mean(log(FS_AA)))
    FSwAa = exp(mean(log(FS_Aa)))
    FSwaa = exp(mean(log(FS_aa)))
    
    # Offspring viability from loci under FS
    FSFit = Fitness(N=N_eq, wAA = FSwAA, wAa = FSwAa, waa = FSwaa, mu = mu, bita = bita, loci = FSloci)
  }else{
    FSFit = 1 
  }
  
  
  #------------ Inbreeding depression (ID)----------------------
  if (IDloci > 0){
    # Relative fitness of homozygote AA, heterozygote Aa and homozygote aa
    IDwAA  <- 1
    IDwAa  <- 1 - ID_h*IDsa
    IDwaa  <- 1 - IDsa
    
    #Offspring viability from loci under ID
    IDFit = Fitness(N=N_eq, wAA = IDwAA, wAa = IDwAa, waa = IDwaa, mu = mu, bita = bitaID, loci = IDloci)
  }else{
    IDFit = 1 
  }
  
  #------------- Mutation Accumulation (MA)--------------------------
  
  if (MAloci > 0){
    # Relative fitness of homozygote AA, heterozygote Aa and homozygote aa
    MAwAA  <- 1 - MAsA
    MAwAa  <- 1 - MA_h*MAsa
    MAwaa  <- 1 - MAsa
    #Offspring viability from loci under MA
    MAFit = Fitness(N=N_eq, wAA = MAwAA, wAa = MAwAa, waa = MAwaa, mu = mu, bita = bitaMA, loci = MAloci)
  }else{
    MAFit = 1 
  }
  
  #----------------------------------------------------------
  
  #--------------- Negative Frequency-Dependent Selection (FD) -------------------------
  if (FDloci > 0){
    fdfitfunc = function(freqA){
      # Relative fitness of homozygote AA, heterozygote Aa and homozygote aa
      FDwAA = 1-FDsA*freqA
      FDwaa = 1-FDsa*(1-freqA)
      FDwAa = 1-(FDsA*freqA + FDsa*(1-freqA))/2
      OUT = c(FDwAA, FDwAa, FDwaa)
      return(OUT)
    }
    ##Transition matrix, state is number of copies of the A allele and takes values 0,1,2,...,2N.
    TransMat  <- matrix(0,nrow=2*N_eq+1,ncol=2*N_eq+1)
    ##The entry in row j and column k is the probability that the population goes from state j to state k in one generation.
    for (j in 0:(2*N_eq)){
      p  <- j/(2*N_eq)#frequency of allele A before selection
      
      pmut = Allefreq(p=p, wAA=fdfitfunc(freqA = p)[1], wAa=fdfitfunc(freqA = p)[2], waa=fdfitfunc(freqA = p)[3], mu = mu, bita=bita)
      
      ## binomial sampling probabilities
     for (k in 0:(2*N_eq)){
        if (N_eq<=500){
          TransMat[j+1,k+1] <- choose(2*N_eq,k) * pmut^k * (1-pmut)^(2*N_eq-k) # Binomial sampling
        }else if(2*N_eq*pmut <PoisThreshold){
          TransMat[j+1,k+1] <- dpois(x=k, lambda = 2*N_eq*pmut)
        }else if(2*N_eq*(1-pmut) <PoisThreshold){
          TransMat[j+1,k+1] <- dpois(x=(2*N_eq-k), lambda = 2*N_eq*(1-pmut))
        }else{
            TransMat[j+1,k+1] <- dnorm(x=k, mean = 2*N_eq*pmut, sd = sqrt(2*N_eq*pmut*(1-pmut))) # Normal distribution approximation  
          }
        
      }
    }

    
    ## get eigenvalues and eigenvectors
    eigen  <- eigen(t(TransMat))
    #get stationary distribution as eigenvector belonging to the eigenvalue with largest real part (should actually always be 1)
    unnormalizedeq <- Re(eigen$vectors[,which(Re(eigen$values)==max(Re(eigen$values)))])
    normalizedeq  <- unnormalizedeq/(sum(unnormalizedeq))
    
    FreqA = (0:(2*N_eq))/(2*N_eq)
    fdwAA = 1-FDsA*FreqA
    fdwaa = 1-FDsa*(1-FreqA)
    fdwAa = 1-(FDsA*FreqA + FDsa*(1-FreqA))/2
    
    fdMeanFitness = function(x){x^2*fdwAA + 2*x*(1-x)*fdwAa + (1-x)^2*fdwaa} # Mean fitness in an infinite population
    
    #Offspring viability from loci under FD
    FDFit = sum(normalizedeq * fdMeanFitness(x=(0:(2*N_eq))/(2*N_eq)))^FDloci
    
    
  }else{
    FDFit = 1 
  }
  
  #----------------------------------------
  
  #------------ Population Absolute Fitness -----------------------
  PopFitness = HAFit * FSFit * FDFit * IDFit * MAFit * exp(Rate) 
  return(PopFitness)
}
```
