## Supplementary material for "Modeling minimum viable population size with multiple genetic problems of small populations": R codes used to simulate results: MainFunct_MarkovChain_POPequiApproach_VariableLoci.R

```
# Main function for the Markov Chain with Variable number of loci
# Function that returns allele frequency after selection and mutation
Allefreq <- function(p, wAA, wAa, waa, mu, bita){
  wbar  <- p^2 * wAA + 2*p*(1-p)*wAa + (1-p)^2 * waa #assuming Hardy-Weinberg equilibrium
  pselect  <- (p^2 * wAA + p*(1-p)*wAa)/wbar #expected allele frequency after selection
  pmut  <- pselect * (1-mu) + (1-pselect)*bita #expected allele frequency after selection and mutation
  return(pmut)
}

PoisThreshold = 35
# Function that calculates the transition matrix, the distribution and then returns the offspring viability.
LocusMeanFitness <- function(N, wAA, wAa, waa, mu, bita){
  ##Transition matrix, state is number of copies of the A allele and takes values 0,1,2,...,2N.
  TransMat  <- matrix(0,nrow=2*N+1,ncol=2*N+1)
  ##The entry in row j and column k is the probability that the population goes from state j to state k in one generation.
  for (j in 0:(2*N)){
    p  <- j/(2*N)#frequency of allele A before selection
    pmut <- Allefreq(p=p, wAA=wAA, wAa=wAa, waa=waa, mu = mu, bita=bita)
    
    ## binomial sampling probabilities
    for (k in 0:(2*N)){
      if (N<=500){
        TransMat[j+1,k+1] <- choose(2*N,k) * pmut^k * (1-pmut)^(2*N-k)
      }else if(2*N*pmut <PoisThreshold){
        TransMat[j+1,k+1] <- dpois(x=k, lambda = 2*N*pmut)
      }else if(2*N*(1-pmut) <PoisThreshold){
        TransMat[j+1,k+1] <- dpois(x=(2*N-k), lambda = 2*N*(1-pmut))
      }else{
        TransMat[j+1,k+1] <- dnorm(x=k, mean = 2*N*pmut, sd = sqrt(2*N*pmut*(1-pmut)))
      }
      
      
    }
  }
  
  
  ## get eigenvalues and eigenvectors
  eigen  <- eigen(t(TransMat))
  #get stationary distribution as eigenvector belonging to the eigenvalue with largest real part (should actually always be 1)
  unnormalizedeq <- Re(eigen$vectors[,which(Re(eigen$values)==max(Re(eigen$values)))])
  normalizedeq  <- unnormalizedeq/(sum(unnormalizedeq))
  
  MeanFitness <- function(x){x^2*wAA + 2*x*(1-x)*wAa + (1-x)^2*waa} # Mean fitness in an infinite population
  
  #FFit <- sum(normalizedeq * MeanFitness(x=(0:(2*N))/(2*N)))^loci
  FFit <- sum(normalizedeq * MeanFitness(x=(0:(2*N))/(2*N))) # Calculating offspring viability at one locus
  #return(FFit)
  return(FFit)
}

#Main Function that considers all the selection mechanisms and uses the above functions. 
MainFuncPoisPeter <- function(LociRatioVector, N_eq, MAsA = 0, MAsa, MA_h=0, bitaID, bitaMA, IDsa, ID_h=0, FDsA, FDsa, FSsa, FSsA, percent, kapa, HAsA, HAsa, mu, bita){
  
  #parallelization of this part such that the transition matrix and so is the distribution and finally offspring viability at one locus is calculated for all possible population sizes from different cores at ago.
  LocusMeanFitness = mclapply(XX, FUN = function(ne){
  #------------------------ Loci ratios ------------
    # Here we mainly check which selection mechanisms we calculate the offspring viability at one locus
  HAloci <- LociRatioVector[1] # Ratio under HA
  XX = 1:length(N_eq)
  FDloci <- LociRatioVector[3] # Ratio under FD
  IDloci <- LociRatioVector[4] # Ratio under ID
  MAloci <- LociRatioVector[5] # Ratio under MA
  #-------------------------------
  FSloci <- LociRatioVector[2] # Ratio under FS
  
  #--------- Heterozygote Advantage (HA)---------------------------------
  if (HAloci > 0){
    # Relative fitness for genotypes AA, Aa and aa
    HAwAA  <- 1-HAsA
    HAwAa  <- 1
    HAwaa  <- 1-HAsa
    
    # Offspring viabilty per locus
    HAFit <- LocusMeanFitness(N=N_eq[ne], wAA = HAwAA, wAa = HAwAa, waa = HAwaa, mu = mu, bita = bita)
  }else{
    HAFit <- 1
  }
  
  #-------------- Fluctuating selection with reversal of dominance (FS)------------------
  if(FSloci > 0){
    Ti <- 1:kapa #Generations in a complete cycle 
    fluct_s <- FSsa*sin(2*pi*Ti/kapa + pi) # selection coefficient for AA
    fluct_t <- FSsA*sin(2*pi*Ti/kapa)
    fluct_h <- 0.5-(percent/100)*sin(2*pi*Ti/kapa)
    
    # Relative fitness for genotypes AA, Aa and aa
    FS_AA <- (1-fluct_t)/(1+max(FSsa,FSsA))
    FS_aa <- (1-fluct_s)/(1+max(FSsa,FSsA))
    FS_Aa <-  (1-fluct_h*fluct_t-(1-fluct_h)*fluct_s)/(1+max(FSsa,FSsA))
    
    FSwAA <- exp(mean(log(FS_AA)))
    FSwAa <- exp(mean(log(FS_Aa)))
    FSwaa <- exp(mean(log(FS_aa)))
    
    # Offspring viabilty per locus
    FSFit <- LocusMeanFitness(N=N_eq[ne], wAA = FSwAA, wAa = FSwAa, waa = FSwaa, mu = mu, bita = bita)
  }else{
    FSFit <- 1
  }
  
  
  #------------ Inbreeding depression (ID) ----------------------
  if (IDloci > 0){
    # Relative fitness for genotypes AA, Aa and aa
    IDwAA  <- 1
    IDwAa  <- 1 - ID_h*IDsa
    IDwaa  <- 1 - IDsa
    
    # Offspring viabilty per locus
    IDFit <- LocusMeanFitness(N=N_eq[ne], wAA = IDwAA, wAa = IDwAa, waa = IDwaa, mu = mu, bita = bitaID)
  }else{
    IDFit <- 1
  }
  
  #------------- Mutation Accumulation (MA)--------------------------
  
  if (MAloci > 0){
    # Relative fitness for genotypes AA, Aa and aa
    MAwAA  <- 1 - MAsA
    MAwAa  <- 1 - MA_h*MAsa
    MAwaa  <- 1 - MAsa
    
    # Offspring viabilty per locus
    MAFit <- LocusMeanFitness(N=N_eq[ne], wAA = MAwAA, wAa = MAwAa, waa = MAwaa, mu = mu, bita = bitaMA)
  }else{
    MAFit <- 1
  }
  
  #--------------- Negative Frequency-Dependent Selection (FD)-------------------------
  if (FDloci > 0){
    fdfitfunc <- function(freqA){
      # Relative fitness for genotypes AA, Aa and aa
      FDwAA <- 1-FDsA*freqA
      FDwaa <- 1-FDsa*(1-freqA)
      FDwAa <- 1-(FDsA*freqA + FDsa*(1-freqA))/2
      OUT <- c(FDwAA, FDwAa, FDwaa)
      return(OUT)
    }
    ##Transition matrix, state is number of copies of the A allele and takes values 0,1,2,...,2N.
    TransMat  <- matrix(0,nrow=2*N_eq[ne]+1,ncol=2*N_eq[ne]+1)
    ##The entry in row j and column k is the probability that the population goes from state j to state k in one generation.
    for (j in 0:(2*N_eq[ne])){
      p  <- j/(2*N_eq[ne])#frequency of allele A before selection
      
      pmut <- Allefreq(p=p, wAA=fdfitfunc(freqA = p)[1], wAa=fdfitfunc(freqA = p)[2], waa=fdfitfunc(freqA = p)[3], mu = mu, bita=bita)
      
      ## binomial sampling probabilities with their approximations
      for (k in 0:(2*N_eq[ne])){
        if (N_eq[ne]<=500){
          TransMat[j+1,k+1] <- choose(2*N_eq[ne],k) * pmut^k * (1-pmut)^(2*N_eq[ne]-k)
        }else if(2*N_eq[ne]*pmut <PoisThreshold){
          TransMat[j+1,k+1] <- dpois(x=k, lambda = 2*N_eq[ne]*pmut)
        }else if(2*N_eq[ne]*(1-pmut) <PoisThreshold){
          TransMat[j+1,k+1] <- dpois(x=(2*N_eq[ne])-k, lambda = 2*N_eq[ne]*(1-pmut))
        }else{
          TransMat[j+1,k+1] <- dnorm(x=k, mean = 2*N_eq[ne]*pmut, sd = sqrt(2*N_eq[ne]*pmut*(1-pmut)))
        }
        
      }
    }
    ## get eigenvalues and eigenvectors
    eigen  <- eigen(t(TransMat))
    #get stationary distribution as eigenvector belonging to the eigenvalue with largest real part (should actually always be 1)
    unnormalizedeq <- Re(eigen$vectors[,which(Re(eigen$values)==max(Re(eigen$values)))])
    normalizedeq  <- unnormalizedeq/(sum(unnormalizedeq))
    
    FreqA <- (0:(2*N_eq[ne]))/(2*N_eq[ne]) # Frequency of A allele 
    #Relative fitness of genotypes AA, aa and Aa
    fdwAA <- 1-FDsA*FreqA
    fdwaa <- 1-FDsa*(1-FreqA)
    fdwAa <- 1-(FDsA*FreqA + FDsa*(1-FreqA))/2
    
    fdMeanFitness <- function(x){x^2*fdwAA + 2*x*(1-x)*fdwAa + (1-x)^2*fdwaa} # Mean fitness in an infinite population size 
    
    # Offspring viabilty per locus
    FDFit <- sum(normalizedeq * fdMeanFitness(x=(0:(2*N_eq[ne]))/(2*N_eq[ne])))
    
  }else{
    FDFit <- 1
  }
  
  print(c("Ne:-", N_eq[ne], HAFit, FSFit, FDFit, IDFit, MAFit))
 
 # Return Population size and offspring viability per locus for HA, FS, FD, ID and MA
  return(c(N_eq[ne], HAFit, FSFit, FDFit, IDFit, MAFit))
  }, mc.preschedule = FALSE, mc.cores = cores)
  
  # Section for assembling locus fitness at each population size for different mechanisms in a matrix. 
  # Here, the first column is the different population sizes, columns 2 to 6 correspond to per locus offspring viability for HA, FS, FD, ID and MA respectively.
  LocusFitness = matrix(0, nrow = length(N_eq), ncol = 6)
  LocusFitness[,1] = N_eq
  
  NeLL = unlist(LocusMeanFitness)[seq(1,6*length(N_eq),6)]
  NeLLorder = NeLL[order(NeLL)]
  
  # HA
  HetLLfit = unlist(LocusMeanFitness)[seq(2,6*length(N_eq),6)]
  Hetfitorder = HetLLfit[order(NeLL)]
  LocusFitness[,2] = Hetfitorder
  
  #FS
  FluctLLfit = unlist(LocusMeanFitness)[seq(3,6*length(N_eq),6)]
  Fluctfitorder = FluctLLfit[order(NeLL)]
  LocusFitness[,3] = Fluctfitorder
  
  #FD
  FreqLLfit = unlist(LocusMeanFitness)[seq(4,6*length(N_eq),6)]
  Freqfitorder = FreqLLfit[order(NeLL)]
  LocusFitness[,4] = Freqfitorder
  
  # ID
  InbLLfit = unlist(LocusMeanFitness)[seq(5,6*length(N_eq),6)]
  Inbfitorder = InbLLfit[order(NeLL)]
  LocusFitness[,5] = Inbfitorder
  
  # MA
  MutaccLLfit = unlist(LocusMeanFitness)[seq(6,6*length(N_eq),6)]
  Mutaccfitorder = MutaccLLfit[order(NeLL)]
  LocusFitness[,6] = Mutaccfitorder

  #Finally, this function returns the matrix.
return(LocusFitness)
}
```
