## Supplementary material for "Modeling minimum viable population size with multiple genetic problems of small populations": R codes used to simulate results: MVP_GeneticProblems_PLOTS.R

```
# Here, we do plotting of all the results both in the Results section and the supplement
# Note that they are not exactly combined like in the main text but you will be able to plot different scenarios separately.
# In this script, we load our csv saved results from the R-scripts that start with "Execute---"
#PLEASE MAKE SURE YOU HAVE RUN THE NECESSARY Execute... scripts and that the results are stored in the 
#named folder.
# This and all other scripts should be in the same folder!!
rm(list=ls())
# Package mainly for plots. 
library("viridis")
#---------------------------------------- Loading MCA and DAA Analysis ----------------------
#MAKE SURE YOU HAVE RUN THE SCRIPTS "Execute_DiffusionApproximation_FixedLoci.R", "Execute_DiffusionApproximation_VariableLoci.R",  "Execute_MarkovChain_POPequiApproach_VariableLoci.R", and "Execute_MarkovChain_POPequiApproach_FixedLoci.R". And THAT THE GENERATED RESULTS ARE STORED ACCORDINGLY (i.e., in "MVP_ResultsFolder" located in your working directory)

#------------------- Variable loci under DAA ----------------------

setwd("~/MVP_GENETICPROBLEMS")
ts.folder <- file.path(paste(getwd(), "/MVP_ResultsFolder", sep = "" ))

setwd(ts.folder)

#Initial starting allele frequency. HA, FS and FD is 0.5 and ID and MA is 0.95 
Startfreq = c(0.5,0.5,0.5,0.95,0.95)

# Do not change these set parameters! Not used in this paper!
initStoch = 1
Envmax = 1
Envrate = 1
capaStoch  = 0
growthStoch = 0
#-------------------------------------

rate = log(1.53) # intrinsic growth rate
Tmax = 400 # Maximum simulation time
# Used Tmax = 4000 # Maximum simulation time
kk = 20 # Discretization of allele frequency. reciprocal or step length   
Tau = 1 # Discretization of time. Step length between time.
kapa = 50 # Generations in a complete cycle
Percent = 40 # parameter for FS
#------------------------------------------------------------------------
tol = 1e-39 # used during taking of inverse of the matrix 1 + alpha*RR
#Selection coeficients 
Mutaccsa = 0.005
Inbsa = 0.05
Hetsa = 0.005
HetsA = 0.005
Freqsa = 0.005
FreqsA = 0.005
FlsA = 0.005
Flsa = 0.005
# Dominance Coefficents
Inbhh = 0.01
Mutacchh = 0.5

# Mutation rates
mutrate = 0.0001
mut_a = mutrate
mut_A = mutrate
inmut_a = mutrate/100
mamut_a = mutrate/100

# Inital population size = Carrying capacity
#Founderpop = round(seq(10, 100, by = 10),0)
#Used Founderpop = round(seq(10, 150000, by = 20),0)
#capacity = Founderpop

MVPvariablelociDAAbelow100 <- read.csv(paste(ts.folder,"/MVPvariablelociDAAbelow100", "_mut_", mut_a, "balsel",Hetsa, "inbsa", Inbsa, "inbhh",Inbhh, "mutaccsa", Mutaccsa, "mutacchh", Mutacchh,"tmax",Tmax, ".csv", sep = ""))

MVPvariablelociDAAabove100 <- read.csv(paste(ts.folder,"/MVPvariablelociDAAabove100", "_mut_", mut_a, "balsel",Hetsa, "inbsa", Inbsa, "inbhh",Inbhh, "mutaccsa", Mutaccsa, "mutacchh", Mutacchh,"tmax",Tmax, ".csv", sep = ""))

cala =c("HA", "FS","ID", "MA")
##----------- Choose which combination is to be plotted.------------------------ 
nn = c( "HA.FS","HA.MA","HA.ID","HA.HA")
#nn= c("FS.HA","FS.MA","FS.ID","FS.FS" )
#nn = c( "MA.HA","MA.FS","MA.ID","MA.MA")
#nn = c("ID.HA","ID.FS","ID.MA","ID.ID")
Calas = order(nn)


llom = MVPvariablelociDAAabove100[,2][max(which(MVPvariablelociDAAabove100[,nn[1:4]]>0, arr.ind = TRUE))]
llom = llom + 5
par(mgp=c(2.9,1,0),mar=c(5,5,1,1) + 0.1)
cca = unlist(lapply(nn, function(x) which(cala==substring(x, 4, 5)) ))#
matplot(log = 'y',100+MVPvariablelociDAAabove100[,2], MVPvariablelociDAAabove100[,nn[1:4]], xlim = c(1,llom+100), ylim = c(min(MVPvariablelociDAAabove100[,nn],MVPvariablelociDAAbelow100[,nn[4]], na.rm = TRUE), max(MVPvariablelociDAAabove100[,nn], MVPvariablelociDAAbelow100[,nn[4]], na.rm = TRUE)), 
        cex.axis = 2.0, col = cca, lty = 1, lwd = 3, ty = 'l', xlab = list("Number of loci acted on", cex=2.8, font =4), ylab = list("Estimated MVP (log-scale)", cex=2.8, font =4))

matlines(MVPvariablelociDAAbelow100[,2], MVPvariablelociDAAbelow100[,nn[4]],col = cca[4], lty = 1, lwd = 3,)

abline(v=100, col="gray", lty=2, lwd=3)

for(i in 1:length(nn)){
  cau = which(cala==substring(nn[i], 4, 5))
  cal = which(cala==substring(nn[i], 1, 2))
  text(100+MVPvariablelociDAAabove100[,2][max(which(MVPvariablelociDAAabove100[,nn[i]]>0))],max(MVPvariablelociDAAabove100[,nn[i]],na.rm = TRUE), substring(nn[i],4,5), col = cau, cex = 2, font = 2)
  
}
text(1,min(MVPvariablelociDAAbelow100[,nn[4]][-1],na.rm = TRUE), cala[cca[4]], col = cca[4], cex = 2, font = 4 )


#------------------- Variable loci under MCA ----------------------
setwd("~/MVP_GENETICPROBLEMS")
ts.folder <- file.path(paste(getwd(),"/MVP_ResultsFolder", sep = "" ))

setwd(ts.folder)

# mutation rates
Mu  <- 0.0001 
Bita = Mu
BitaID = Mu/100
BitaMA = Mu/100

# Population sizes/carrying capacity vector 
#n_eq  <- round(seq(10, 3150, by = 20),0)
rate = log(1.53) # intrinsic growth rate

# Selection coefficients
hasA = 0.005
hasa = 0.005
fssa = 0.005
fssA = 0.005
fdsA = 0.005
fdsa = 0.005
idsa = 0.05
masA = 0
masa = 0.005
#--------------
Percent = 40
Kapa = 50 # Number of generations in a complete cycle
# Dominance coeeficients for ID and MA
id_h = 0.01 
ma_h = 0.5

MVPvariablelociMCAbelow100 <- read.csv(paste(ts.folder,"/MVPvariablelociMCAbelow100", "_mut_", Mu, "balsel",hasa, "inbsa", idsa, "inbhh",id_h, "mutaccsa", masa, "mutacchh", ma_h, ".csv", sep = ""))

MVPvariablelociMCAabove100 <- read.csv(paste(ts.folder,"/MVPvariablelociMCAabove100", "_mut_", Mu, "balsel",hasa, "inbsa", idsa, "inbhh",id_h, "mutaccsa", masa, "mutacchh", ma_h, ".csv", sep = ""))

cala =c("HA", "FS","ID", "MA") # Names of different selection mechanisms

##----------- Choose which combination is to be plotted.------------------------ 
#nn = c( "HA.FS","HA.MA","HA.ID","HA.HA")
#nn= c("FS.HA","FS.MA","FS.ID","FS.FS" )
nn = c( "MA.HA","MA.FS","MA.ID","MA.MA")
#nn = c("ID.HA","ID.FS","ID.MA","ID.ID")
Calas = order(nn)


llom = MVPvariablelociMCAabove100[,2][max(which(MVPvariablelociMCAabove100[,nn[1:4]]>0, arr.ind = TRUE))]
llom = llom + 5
par(mgp=c(2.9,1,0),mar=c(5,5,1,1) + 0.1)
cca = unlist(lapply(nn, function(x) which(cala==substring(x, 4, 5)) ))#
matplot(log = 'y',100+MVPvariablelociMCAabove100[,2], MVPvariablelociMCAabove100[,nn[1:4]], xlim = c(1,llom+100), ylim = c(min(MVPvariablelociMCAabove100[,nn],MVPvariablelociMCAbelow100[,nn[4]], na.rm = TRUE), max(MVPvariablelociMCAabove100[,nn], MVPvariablelociMCAbelow100[,nn[4]], na.rm = TRUE)), 
        cex.axis = 2.0, col = cca, lty = 1, lwd = 3, ty = 'l', xlab = list("Number of loci acted on", cex=2.8, font =4), ylab = list("Estimated MVP (log-scale)", cex=2.8, font =4))

matlines(MVPvariablelociMCAbelow100[,2], MVPvariablelociMCAbelow100[,nn[4]],col = cca[4], lty = 1, lwd = 3,)

abline(v=100, col="gray", lty=2, lwd=3)

for(i in 1:length(nn)){
  cau = which(cala==substring(nn[i], 4, 5))
  cal = which(cala==substring(nn[i], 1, 2))
  text(100+MVPvariablelociMCAabove100[,2][max(which(MVPvariablelociMCAabove100[,nn[i]]>0))],max(MVPvariablelociMCAabove100[,nn[i]],na.rm = TRUE), substring(nn[i],4,5), col = cau, cex = 2, font = 2)
  
}
text(1,min(MVPvariablelociMCAbelow100[,nn[4]][-1],na.rm = TRUE), cala[cca[4]], col = cca[4], cex = 2, font = 4 )


#------------------- Fixed loci under MCA ----------------------
setwd("~/MVP_GENETICPROBLEMS")
ts.folder <- file.path(getwd(), paste("/MVP_ResultsFolder", sep = "" ))


setwd(ts.folder)


#--------------------------------------------------------------
Mu  <- round(seq(0.00005, 0.0002, length.out = 2),5) #mutation rate
#Used Mu  <- round(seq(0.00005, 0.002, length.out = 10),5) #mutation rate
Bita = Mu
BitaID = Mu/100
BitaMA = Mu/100
#n_eq  <- round(seq(10, 3500,length.out = 350),0)
rate = log(1.53)

#selection coefficients
hasA = 0.005
hasa = 0.005
fssa = 0.005
fssA = 0.005
fdsA = 0.005
fdsa = 0.005
idsa = 0.05
masA = 0
masa = 0.005

Percent = 40
Kapa = 50 # Number of generations in a complete cycle

# Dominance coefficients for ID and MA
id_h = 0.01
ma_h = 0.5

MVPfixedMCA <- read.csv(paste(ts.folder,"/MVPfixedlociMCAEqui_","fit_r0log_",round(exp(rate),4),"_hsa_", hasa, "_hsA_", hasA, "_flsa_", fssa,
                              "_flsA_", fssA, "_frsa_", fdsa, "_frsA_", fdsA, "_insa_",idsa, "_inh_", id_h, "_masa_", masa, "_masA_", masA, "_mah_", ma_h, "_kapa_", Kapa, ".csv", sep = ""))


FDlist = c("FD","HAFD","FSFD","FDID","FDMA","HAFSFD","HAFDID","FSFDID", "FDIDMA","HAFDMA","FSFDMA","HAFSFDID","HAFSFDMA","HAFDIDMA","HAFSFDIDMA","FSFDIDMA", "FDMAID", "HAFDMAID", "FSFDMAID", "HAFSFDMAID")
MVPfixedlociMCA = MVPfixedMCA[,colnames(MVPfixedMCA)[-which(colnames(MVPfixedMCA) %in% FDlist)]]

par(mgp=c(2.9,1,0),mar=c(5,5,2,2) + 0.1)
# Choose which combination of selection mechanisms is to be plotted.
cc = 1:4
#cc = 5:10
#cc = 11:15
cc = cc+1
matplot(Mu, MVPfixedlociMCA[,cc], ty = 'o', log = "y", pch = 16, lty = 1, lwd=3,
        cex.axis = 2.0, ylim = c(min(MVPfixedlociMCA[,cc], na.rm = TRUE), max(MVPfixedlociMCA[,cc], na.rm = TRUE)),
        ylab = list("MVP (log-scale)", font = 4, cex = 2.8), xlab = list("Mutation rate", font = 4, cex = 2.8))
legend("bottom", legend = c(colnames(MVPfixedlociMCA)[cc]), text.font = 4, cex=1.5, col = c(1:length(cc)), lty=c(1), pch = 16, lwd=3)


#------------------- Fixed loci under DAA ----------------------
setwd("~/MVP_GENETICPROBLEMS")
ts.folder <- file.path(getwd(), paste("/MVP_ResultsFolder", sep = "" ))


setwd(ts.folder)


# Inital population size = Carrying capacity
#Founderpop = round(seq(10, 3500, length.out = 350),0)
#capacity = Founderpop

#Initial starting allele frequency. HA, FS and FD is 0.5 and ID and MA is 0.95 
Startfreq = c(0.5,0.5,0.5,0.95,0.95)

# Do not change these set parameters! Not used in this paper!
initStoch = 1
Envmax = 1
Envrate = 1
capaStoch  = 0
growthStoch = 0

rate = log(1.53) # intrinsic growth rate
Tmax = 400 # Maximum simulation time
#Used Tmax = 4000 # Maximum simulation time
kk = 20 # Discretization of allele frequency. reciprocal or step length   
Tau = 1 # Discretization of time. Step length between time.
kapa = 50 # Generations in a complete cycle
Percent = 40 # parameter for FS
#------------------------------------------------------------------------
tol = 1e-39 # used during taking of inverse of the matrix 1 + alpha*RR
#Selection coeficients 
Mutaccsa = 0.005
Inbsa = 0.05
Hetsa = 0.005
HetsA = 0.005
Freqsa = 0.005
FreqsA = 0.005
FlsA = 0.005
Flsa = 0.005
# Dominance Coefficents
Inbhh = 0.01
Mutacchh = 0.5

# Mutation rates
mutrate = round(seq(0.00005, 0.0002, length.out = 2),5)
# Used mutrate = round(seq(0.00005, 0.002, length.out = 10),5)
mut_a = mutrate
mut_A = mutrate
inmut_a = mutrate/100
mamut_a = mutrate/100

MVPfixedDAA <- read.csv(paste(ts.folder,"/MVPfixedlociDAA_","fit_r0log_",round(exp(rate),4),"_hsa_", Hetsa, "_hsA_", HetsA, "_flsa_", Flsa, "_flsA_", FlsA, "_frsa_", Freqsa, 
                              "_frsA_", FreqsA, "_insa_",Inbsa, "_inh_", Inbhh, "_masa_", Mutaccsa, "_mah_", Mutacchh, "_kapa_", kapa,"tmax", Tmax,  ".csv", sep = ""))


FDlist = c("FD","HAFD","FSFD","FDID","FDMA","HAFSFD","HAFDID","FSFDID", "FDIDMA","HAFDMA","FSFDMA","HAFSFDID","HAFSFDMA","HAFDIDMA","HAFSFDIDMA","FSFDIDMA", "FDMAID", "HAFDMAID", "FSFDMAID", "HAFSFDMAID")
MVPfixedlociDAA = MVPfixedDAA[,colnames(MVPfixedDAA)[-which(colnames(MVPfixedDAA) %in% FDlist)]]

par(mgp=c(2.9,1,0),mar=c(5,5,2,2) + 0.1)
# Choose which combination of selection mechanisms is to be plotted.
cc = 1:4
#cc = 5:10
#cc = 11:15
cc = cc+1
matplot(mutrate, MVPfixedlociDAA[,cc], ty = 'o', log = "y", pch = 16, lty = 1, lwd=3,
        cex.axis = 2.0, ylim = c(min(MVPfixedlociDAA[,cc], na.rm = TRUE), max(MVPfixedlociDAA[,cc], na.rm = TRUE)),
        ylab = list("MVP (log-scale)", font = 4, cex = 2.8), xlab = list("Mutation rate", font = 4, cex = 2.8))
legend("bottom", legend = c(colnames(MVPfixedlociDAA)[cc]), text.font = 4, cex=1.5, col = c(1:length(cc)), lty=c(1), pch = 16, lwd=3)


#-------------------------------- Population time trajectories under fixed loci, DAA ----------------------
#------------- Section for Population and fitness trajectories ------------------------------
rm(list=ls())
# Packages needed
library("doMC")
cores=detectCores()
registerDoMC(cores)
require(reshape2)

setwd("~/MVP_GENETICPROBLEMS")
source("MainFunct_DiffusionApproximation.R")

# Do not change these set parameters! Not used in this paper!
initStoch = 1
Envmax = 1
Envrate = 1
capaStoch  = 0
growthStoch = 0
#-------------------------------------------

rate = log(1.53) # intrinsic growth rate
Tmax = 4000 # Maximum simulation time
kk = 20 # Discretization of allele frequency. reciprocal or step length   
Tau = 1 # Discretization of time. Step length between time.
kapa = 50 # Generations in a complete cycle
Percent = 40 # parameter for FS
#------------------------------------------------------------------------
tol = 1e-39 # used during taking of inverse of the matrix 1 + alpha*RR
#Selection coeficients 
Mutaccsa = 0.005
Inbsa = 0.05
Hetsa = 0.005
HetsA = 0.005
Freqsa = 0.005
FreqsA = 0.005
FlsA = 0.005
Flsa = 0.005
# Dominance Coefficents
Inbhh = 0.01
Mutacchh = 0.5

# Mutation rates
mutrateTrajectories = 0.0005
mut_a = mutrateTrajectories
mut_A = mutrateTrajectories
inmut_a = mutrateTrajectories/100
mamut_a = mutrateTrajectories/100
Numberofloci = 100
Startfreq = c(0.5,0.5,0.5,0.95,0.95)
lociRatioVectorTrajectories = c(0.5, 0.0, 0.0, 0.5, 0.0) # ratio vector in the order HA, FS, FD, ID and MA number of loci. Shoulöd sum to 1.

FounderpopTrajectories = round(seq(10, 70, by = 10),0)
#FounderpopTrajectories = c(100,150,170,200,250)
(capacityTrajectories = FounderpopTrajectories)
TrajectoriesDAA = foreach(pop = 1:length(capacityTrajectories), .combine='cbind') %dopar% {
  RRes = ApproxMixFunc(startfreq = Startfreq, numberofloci = Numberofloci, founderpop = FounderpopTrajectories[pop], Capacity = capacityTrajectories[pop], Rate = rate, tmax = Tmax, tau = Tau, KK = kk, Kapa = kapa, percent = Percent, Mut_a = mut_a, inMut_a = inmut_a, maMut_a = mamut_a,
                       Mut_A = mut_A, envmax = Envmax, envrate = Envrate, InitStoch = initStoch, CapaStoch = capaStoch, GrowthStoch = growthStoch, hetsa = Hetsa, hetsA = HetsA, flsa = Flsa, flsA = FlsA, freqsa = Freqsa,
                       freqsA = FreqsA, inbsa = Inbsa, inbhh=Inbhh, LociRatioVector = lociRatioVectorTrajectories, Tol = tol, mutaccsa = Mutaccsa, mutacchh=Mutacchh)
  
}


ttb = 4000 # x-axis time upper limit to be displayed
co= min(c(6, length(capacityTrajectories))) # Choose how many values of carrying capacity to be plotted 

# Population size trajectories - Main text
par(mgp=c(2.9,1,0),mar=c(5,5,1,1) + 0.1)
matplot(TrajectoriesDAA[,rev(which(colnames(TrajectoriesDAA)=="POPSIZE"))[round(seq(1,length(capacityTrajectories),length.out = co),0)]], cex.axis=2.0, ty='l', lwd=3, xlim = c(0,ttb), lty=1,col = rev(viridis(length(capacityTrajectories))),
        ylab = list("Population size", font = 4, cex=2.8), xlab = list("Time", font = 4, cex=2.8))
legend("bottomright", legend = c("K", capacityTrajectories[round(seq(1,length(capacityTrajectories),length.out = co),0)]), lwd=3, col = c(NA,viridis(length(capacityTrajectories))[round(seq(1,length(capacityTrajectories),length.out = co),co)]), text.font = 4, cex = 2.0, lty=1 )

# Fitness trajectories - Main text 
par(mgp=c(2.9,1,0),mar=c(5,5,1,1) + 0.1)
matplot(1.53*TrajectoriesDAA[,rev(which(colnames(TrajectoriesDAA)=="POPFIT"))[round(seq(1,length(capacityTrajectories),length.out = co),0)]], cex.axis=2.0, lwd=3, ty='l', ylim = c(0.5,max(1.53*TrajectoriesDAA[,which(colnames(TrajectoriesDAA)=="POPFIT")], na.rm = TRUE)), xlim = c(0,ttb), lty=1,col = rev(viridis(length(which(colnames(TrajectoriesDAA)=="POPSIZE")))),
        ylab = list("Absolute fitness", font = 4, cex=2.8), xlab = list("Time", font = 4, cex=2.8))
abline(h=1, col="gray", lwd=2, lty=1)
legend("bottomright", legend = c("K", capacityTrajectories[round(seq(1,length(capacityTrajectories),length.out = co),0)]), lwd=3, col = c(NA,viridis(length(capacityTrajectories))[round(seq(1,length(which(colnames(TrajectoriesDAA)=="POPSIZE")),length.out = co),co)]), text.font = 4, cex=2.0, lty=1 )

# Viability of the population for cases only with at least 2 selection mechanisms
# The columns for different mechanisms viability are "hetWW", "fluctWW", "freqWW", "inbWW", "mutaccWW" for HA, FS, FD, ID, and MA respectively
#Here, the example is for cases with HA and ID

par(mgp=c(2.9,1,0),mar=c(5,5,1,1) + 0.1)
matplot(1.*TrajectoriesDAA[,rev(which(colnames(TrajectoriesDAA)=="inbWW"))[round(seq(1,length(capacityTrajectories),length.out = co),0)]], cex.axis = 2.0, lwd = 3, ty='l', ylim = c(0.75,max(1.*TrajectoriesDAA[,which(colnames(TrajectoriesDAA)=="inbWW")], na.rm = TRUE)), xlim = c(0,ttb),
        lty=1,col = rev(viridis(length(which(colnames(TrajectoriesDAA)=="POPSIZE")))),
        ylab = list("Offspring survival fitness ", font = 4, cex=2.8), xlab = list("Time", font = 4, cex=2.8))
matlines(1.*TrajectoriesDAA[,rev(which(colnames(TrajectoriesDAA)=="hetWW"))[round(seq(1,length(capacityTrajectories),length.out = co),0)]], lwd = 3, lty = 2, col = rev(viridis(length(which(colnames(TrajectoriesDAA)=="POPSIZE")))))

legend("topright", legend = c("K", capacityTrajectories[round(seq(1,length(capacityTrajectories),length.out = co),0)]), lwd=3, col = c(NA,viridis(length(capacityTrajectories))[round(seq(1,length(which(colnames(TrajectoriesDAA)=="POPSIZE")),length.out = co),co)]), text.font = 4, cex=2, lty=1 )
legend("top", legend = c("ID","HA"), lwd=3, col = 4, lty=c(1,2), text.font = 4, cex = 2 )


#---------------------------------------- Loading IBM Analysis ----------------------
#MAKE SURE YOU HAVE RUN SCRIPT "Execute_IBM.R" and THAT THE GENERATED RESULTS ARE STORED ACCORDINGLY

replicates = 1:4 # Number of replicates
# Used replicates = 1:60 # Number of replicates

lociRatioVector = c(1,0,0,0,0)# HA, FS,FD,ID,MA, must sum to 1.

mutrate = round(seq(0.001, 0.002, length.out = 2), 5)
# Used mutrate = round(seq(0, 0.002, length.out = 10), 5)
Founderpop <- round(seq(10, 100, length.out = 10),0) # starting population size for HA
# Used Founderpop <- round(seq(10, 3500, length.out = 350),0) # starting population size for HA

capacity <- Founderpop # Founder population size denoted as N_0 in the text used as carrying capacity


if(length(which(lociRatioVector>0))==3){
  Numberofloci = 99
}else{
  Numberofloci = 100
}


Tmax = 100 # max. simulation time 
# Used Tmax = 4000 # max. simulation time 
x0 = 0.5# Initail frequency of allele A for all mechanisms

# Selection coefficients
Hetsa = 0.005
HetsA = Hetsa
Fluctsa = 0.005
FluctsA = Fluctsa
Freqsa = 0.005
FreqsA = Freqsa
idsa = 0.005
masa = 0.005

# Dominance coefficients
id_h = 0.0
ma_h = 0.5

#Mutation rates
hetmut_A = mutrate
hetmut_a = mutrate
fluctmut_A = mutrate
fluctmut_a = mutrate
freqmut_A = mutrate
freqmut_a = mutrate
inbmut_A = mutrate
inbmut_a = mutrate/100
mamut_A = mutrate
mamut_a = mutrate/100

rate = log(1.53) # Intrinsic growth rate 
Deathrate = 0
Percent = 40 #Percent is a parameter in heterozygote genotype fitness for FS between 0 and 100 (used 40 throught).
kapa = 50 # Number of generations in a complete cycle

Threshold = 1 # Set threshold to stop running. Here all 100% of replicates must survive to stop increasing capacity.

# Do not change these parameters. They are silent in this study
initStoch = 1
Envrate = 0
Envmax = 1
capaStoch = 0
growthStoch = 0

setwd("~/MVP_GENETICPROBLEMS")
ts.folder <- file.path(paste(getwd(),"/MVP_ResultsFolder", sep = "" ))
if(!dir.exists(ts.folder)){
  dir.create(ts.folder)
}

setwd(ts.folder)


ts.folderIBM <- file.path(ts.folder,paste("IBM_TotLoci_",Numberofloci, "_hetloci_", round(lociRatioVector[1],2),  "_fluctloci_", round(lociRatioVector[2],2),
                                        "_freqloci_",  round(lociRatioVector[3],2), "_inbloci_", round(lociRatioVector[4],2), "_mutaccloci_", round(lociRatioVector[5],2), sep = ""  ))
if(!dir.exists(ts.folderIBM)){
  dir.create(ts.folderIBM)
}

setwd(ts.folderIBM)

ts.folder1 <- file.path(getwd(),paste("CapacityMutation", sep = ""  ))
if(!dir.exists(ts.folder1)){
  dir.create(ts.folder1)
}

PSIZE = array(0, dim = c(Tmax, length(replicates), length(Founderpop), length(mutrate)))
MFIT = array(0, dim = c(Tmax, length(replicates),   length(Founderpop), length(mutrate)))


for (pop in 1:length(Founderpop)) {
  for (mut in 1:length(mutrate)){
    
    if(file.exists( paste(ts.folder1,"/POPSIZE_r0_log",exp(rate ),   "_Capacity_", capacity[pop], "_hsa_",  Hetsa, "_hsA_",  HetsA, "_flsa_",  Fluctsa,
                          "_flsA_",  FluctsA, "_frsa_",  Freqsa, "_frsA_",  FreqsA, "_insa_",  idsa,  "_masa_",  masa, "_inh_",  id_h, "_mah_",  ma_h,
                          "_muA_",  hetmut_A[mut], "_mua_",  hetmut_a[mut], "_n_",  Numberofloci, "_hlc_", round(lociRatioVector[1],2),  "_fllc_", round(lociRatioVector[2],2),  "_frlc_",
                          round(lociRatioVector[3],2),  "_inlc_", round(lociRatioVector[4],2),  "_malc_", round(lociRatioVector[5],2), "_kapa_",  kapa,  
                          "_tmax_", Tmax, ".csv", sep = "") )){
      
      PSIZE[,, pop,mut] = as.matrix(read.csv(paste(ts.folder1,"/POPSIZE_r0_log",exp(rate ),   "_Capacity_", capacity[pop], "_hsa_",  Hetsa, "_hsA_",  HetsA, "_flsa_",  Fluctsa,
                                                   "_flsA_",  FluctsA, "_frsa_",  Freqsa, "_frsA_",  FreqsA, "_insa_",  idsa,  "_masa_",  masa, "_inh_",  id_h, "_mah_",  ma_h,
                                                   "_muA_",  hetmut_A[mut], "_mua_",  hetmut_a[mut], "_n_",  Numberofloci, "_hlc_", round(lociRatioVector[1],2),  "_fllc_", round(lociRatioVector[2],2),  "_frlc_",
                                                   round(lociRatioVector[3],2),  "_inlc_", round(lociRatioVector[4],2),  "_malc_", round(lociRatioVector[5],2), "_kapa_",  kapa, 
                                                   "_tmax_", Tmax, ".csv", sep = ""))[,-1])
      
      MFIT[,, pop, mut] = as.matrix(read.csv(paste(ts.folder1,"/MEANFIT_r0_log",exp(rate ),   "_Capacity_", capacity[pop], "_hsa_",  Hetsa, "_hsA_",  HetsA, "_flsa_",  Fluctsa,
                                                   "_flsA_",  FluctsA, "_frsa_",  Freqsa, "_frsA_",  FreqsA, "_insa_",  idsa,  "_masa_",  masa, "_inh_",  id_h, "_mah_",  ma_h,
                                                   "_muA_",  hetmut_A[mut], "_mua_",  hetmut_a[mut], "_n_",  Numberofloci, "_hlc_", round(lociRatioVector[1],2),  "_fllc_", round(lociRatioVector[2],2),  "_frlc_",
                                                   round(lociRatioVector[3],2),  "_inlc_", round(lociRatioVector[4],2),  "_malc_", round(lociRatioVector[5],2), "_kapa_",  kapa, 
                                                   "_tmax_", Tmax, ".csv", sep = ""))[,-1])
      
    }
  }
}

#----------------------- Plots for the IBM ---------------------------------------------
ttb = Tmax # x-axis upper limit
mu = 2
po = 2
mutrate[mu]
capacity[po]
# Population trajectories in the supplement (S5)--------------------
par(mgp=c(2.9,1,0),mar=c(5,5,2,2) + 0.1)
matplot(1:ttb, PSIZE[1:ttb,,po,mu], lty = 1, cex.axis=2, xlim = c(0,ttb), col = adjustcolor("gray", alpha.f = 0.5), ty = 'l',
        ylab = list("Population size", font = 4, cex=2.8), xlab = list("Time", font = 4, cex=2.8))


par(mgp=c(2.9,1,0),mar=c(5,5,2,2) + 0.1)
matplot(1:ttb, 1.53*MFIT[1:ttb,,po,mu], lty = 1, cex.axis=2, ylim = 1.53*c(0.5,1), xlim = c(0,ttb), col = adjustcolor("gray", alpha.f = 0.5), ty = 'l',
        ylab = list("Absolute fitness", font = 4, cex=2.8), xlab = list("Time", font = 4, cex=2.8))

# ----------- Other analysis in the supplement (S6)---------------
YY = apply(PSIZE[Tmax,,, ], c(2,3), FUN = function(y){
  x = length(which(y>0))/length(replicates)
  
})

THRESHOLD = round(seq(0,1, length.out = 20),2)# survival probability levels
MVPth = matrix(NA, nrow = length(mutrate), ncol = length(THRESHOLD))
for(th in 1:length(THRESHOLD)){
  MVPth[,th] = apply(YY, 2, FUN = function(y){Founderpop[min(which(y>=THRESHOLD[th]))]})
}

# --------- MVP vs Threshold ---------------
par(mgp=c(2.9,1,0),mar=c(5,5,2,2) + 0.1)
matplot(THRESHOLD, t(MVPth), cex.axis=2, ty = 'o', col = viridis(n=length(mutrate)), pch = 16, lty = 1, lwd = 2,
        xlab = list("Survival threshold", font = 4, cex = 2.8), ylab = list("MVP", font = 4, cex = 2.8))
legend("bottomright", legend = c("Mut",mutrate[seq(0,length(mutrate), 2)]), cex = 2, text.font = 4, lwd = 2, pch = 16, col = c(NA,viridis(n=length(mutrate[seq(0,length(mutrate), 2)]))))
#legend("bottom", legend = c("Mut",mutrate[seq(0,length(mutrate), 2)]), cex = 2, text.font = 4, lwd = 2, pch = 16, col = c(NA,viridis(n=length(mutrate[seq(0,length(mutrate), 2)]))))


ZZ1 = apply(PSIZE[Tmax,,, ], c(2,3), FUN = function(y){
  x = mean(y>0)
  
})

# Extinction times, mean and coefficient of variation ---------------------
TTraw = apply(PSIZE, c(2,3,4), FUN = function(y){
  x = max(which(y>0), na.rm = TRUE)
  if(is.infinite(x)){x = Tmax}
  return(x)
})

TTmean = apply(TTraw, c(2,3), FUN = function(y){
  x = mean(y, na.rm = TRUE)
  
})


TTcoevar = apply(TTraw, c(2,3), FUN = function(y){
  x = sd(y, na.rm = TRUE)/mean(y, na.rm = TRUE)
  
})

# Mean survival time vs capacity K
par(mgp=c(2.9,1,0),mar=c(5,5,2,2) + 0.1)
matplot(capacity, TTmean, ty ='o', cex.axis=2, col = viridis(n=length(mutrate)), xlim = c(0,700), lty=1, lwd = 2, pch = 16,xlab = list("K", font = 4, cex = 2.8),
        ylab = list("Mean survival time", font = 4, cex = 2.8))
legend("bottomright",legend = c("Mut",mutrate), text.font = 4, cex = 2, lwd = 2, pch = 16, col = c(NA,viridis(n=length(mutrate))))


#------------ Coef. of Variation deviation vs K-------------
par(mgp=c(2.9,1,0),mar=c(5,5,2,2) + 0.1)
matplot(capacity, TTcoevar, ty ='o', cex.axis=2, xlim = c(0,600), col = viridis(n=length(mutrate)), lty=1, lwd = 2, pch = 16,xlab = list("K", font = 4, cex = 2.8),
        ylab = list("Coef. of var. in survival time", font = 4, cex = 2.8))
legend("topright",legend = c("Mut",mutrate), cex = 2, text.font = 4, lwd = 2, pch = 16, col = c(NA,viridis(n=length(mutrate))))


#-----------------------------------------------------------------------------------------------------
#----------------------------- Other supplementary figures (S4)-----------------------
#-----------------------------------------------------------------------------------------------------
#------------------ Illustration of mvp vs fitness in infinite population -------------------------
mutrate = round(seq(0.00005,0.002,length.out = 10),5)
# dom. coef.
mah1 = 0.01
mah2 = 0.5
#sel. coef.
sela1 = 0.005
sela2 = 0.05
nloci=100# Number of loci
#Mfit_MA_h0p01_sela0p05 =(1-2*mutrate)^nloci
#Mfit_MA_h0p5_sela0p005 = (1-2*mutrate)^nloci

Mfit_MA_h0p5_sela0p005 = numeric(length(mutrate))
Mfit_MA_h0p01_sela0p05 = numeric(length(mutrate))
Mfit_MA_h0_sela0p005 = numeric(length(mutrate))
for (mut in 1:length(mutrate)){
  if(mutrate[mut]<(mah2*sela1)){
    Mfit_MA_h0p5_sela0p005[mut] = (1-2*mutrate[mut] + 2*mutrate[mut]^2/(mah2*sela1) - mutrate[mut]^2/(mah2^2*sela1))^nloci
  }else{
    Mfit_MA_h0p5_sela0p005[mut] = (1-sela1)^nloci
  }
  if(mutrate[mut]<(mah1*sela2)){
    Mfit_MA_h0p01_sela0p05[mut] = (1-2*mutrate[mut] + 2*mutrate[mut]^2/(mah2*sela1) - mutrate[mut]^2/(mah2^2*sela1))^nloci
  }else{
    Mfit_MA_h0p01_sela0p05[mut] = (1-sela2)^nloci
  }
  
  if(mutrate[mut]<(sela2)){
    Mfit_MA_h0_sela0p005[mut] = (1-mutrate[mut])^nloci
  }else{
    Mfit_MA_h0_sela0p005[mut] = (1-sela2)^nloci
  }
  
}

mvpMA_h0_s0p005 = c(300, 310, 320, 340, 360, 390, 420, 450, 490, 510) # Extracted from MA or ID with sa=0.005, h = 0.0
mvpMA_h0p5_s0p005 = c(300,  350,  420,  580,  570,  660,  830, 1090, 1550, 3060) #  Extracted from MA or ID with sa=0.005, h = 0.5

# Plotting Estimated MVP vs equilibrium fitness
par(mgp=c(2.9,1,0),mar=c(5,5,2,2) + 0.1)
plot(Mfit_MA_h0p5_sela0p005, mvpMA_h0p5_s0p005, cex.axis=2, ty='o', log="y", xlim = c(0.66,1), ylim = c(300,3060), lwd=3, pch=16, cex=1.3,
     ylab = list("MVP", cex=2.8, font=4), xlab = list("Equilibrium Fitness", cex=2.8, font=4))
lines(Mfit_MA_h0_sela0p005, mvpMA_h0_s0p005, ty='o', col= 2, lwd=3, pch=16, cex=1.3)
legend("topright", legend = c("ID", "h=0.5,s=0.005","h=0.0,s=0.005"), col = c(NA,1:2), lwd = 3, pch = 16, cex = 2, text.font = 2)


#--------------------- Critical number of loci above which no any arbitrary size can survive --------------------
# Mean fitness at infinite populaion size
WFunc = function(sa, sA, h, p){
  wAA = 1-sA
  wAa = 1-h*sa
  waa = 1-sa
  W = p^2*wAA + 2*p*(1-p)*wAa + (1-p)^2*waa
  
  return(W)
}

# Function takes in selection coefficients sa,sA, equilibrium allele frequency p, dominance coefficients h, 
#fertility E and number of loci of original selection mechanism loci0
loci_crit = function(sa0, sA0, h0, p0, sa1, sA1, h1, p1, E, loci0){
  if(loci0>0){
    V0 = WFunc(sa = sa0, sA = sA0, h = h0, p = p0)  
  }else{
    V0 = 1
  }
  
  V1 = WFunc(sa = sa1, sA = sA1, h = h1, p = p1)
  S_c = log(1/(V0^loci0*E))/log(V1)
  return(S_c)
}

#------------ Hetero Advantage (HA)--------------------------
e = 1.53
Sa0 = 0.005
SA0 = 0.005
H0 = 0
P0 = Sa0/(Sa0+SA0)
Loci0 = 0
H1 = 0
Sel1 = seq(0.0001, 0.01, by = 0.0001)
SS_c = matrix(NA, nrow = length(Sel1), ncol = length(Sel1)) 
for(sea in 1:length(Sel1)){
  for(seA in 1:length(Sel1)){
    P1 = Sel1[sea]/(Sel1[sea]+Sel1[seA])
    SS_c[sea,seA] = loci_crit(sa0=Sa0, sA0=SA0, h0=H0, p0=P0, sa1=Sel1[sea], sA1=Sel1[seA], h1=H1, p1=P1, E=e, loci0=Loci0)
  }
}


# Image showing critical number of loci under HA as selection mechanisms in alleles a and A varies
par(mgp=c(2.9,1,0),mar=c(5,5,2,2) + 0.1)
image(Sel1,Sel1, SS_c, zlim = c(0,max(SS_c, na.rm = TRUE)), cex.axis=2, add = F, col = rev(adjustcolor(heat.colors(100), alpha.f = 0)), 
      ylab = list(expression(s[AA]), font = 4, cex=2.8, col = 1), xlab = list(expression(s[aa]), font = 4, cex=2.8, col = 1))

contour(Sel1,Sel1, SS_c, drawlabels = T, zlim = c(10,500), labcex = 1.3, lwd = 3, col = 4, nlevels = 15, add = T, method = "flattest", vfont = c("sans serif", "plain"))

#------------ Inbreeding depression (ID)--------------------------
e = 1.53 # Fertility
muA = 0.0001 # Forward mutation rate
Sa0 = 0.005 # Sllection against recessive deleterious allele a
SA0 = 0.00 # Selection against wild-type allele A
H0 = 0
P0 = sqrt(muA/Sa0) # Equilibrium allele frequency
Loci0 = 00# Number of loci 
H1 = 0.0
Sel1 = seq(0.001, 0.05, by = 0.001) # selection coefficients
MutA = seq(0.00001,0.001, by=0.00001) # Mutation rate forward
SS_c = matrix(NA, nrow = length(MutA), ncol = length(Sel1))
#SS_c = numeric(length(Sel1))
for(sea in 1:length(Sel1)){
  for(MuA in 1:length(MutA)){
    P1 = 1-sqrt(MutA[MuA]/Sel1[sea])
    SS_c[MuA, sea] = loci_crit(sa0=Sa0, sA0=SA0, h0=H0, p0=P0, sa1=Sel1[sea], sA1=0, h1=H1, p1=P1, E=e, loci0=Loci0)
  }
}

# Ploting critical number of loci above which no finite population size survives vs mutation rate
par(mgp=c(2.9,1,0),mar=c(5,5,2,2) + 0.1)
matplot(MutA,SS_c, ty='l', cex.axis = 2, col = c(viridis(length(Sel1))[round(seq(1,length(Sel1),length.out = 10),10)]), lwd = 3,
        ylab = list("Critical number of loci", font = 4, cex = 2.8, col = 1), xlab = list("Forward mutation rate", cex = 2.8, font = 4, col = 1))
legend("topright", legend = c("Sel. Coef",Sel1[round(seq(1,length(Sel1),length.out = 10),10)]), col = c(NA,viridis(length(Sel1))[round(seq(1,length(Sel1),length.out = 10),10)]),
       cex = 2, text.font = 4, lty=1, lwd=1)

n = 500
nloci = 1:n
hetSa=0.005
hetSA = 0.005
hetH = 0
hetP=hetSa/(hetSa+hetSA)
Whet = 1.53*WFunc(sa = hetSa, sA = hetSA, h=hetH, p=hetP)^nloci

#----------------------------------------- Fitness in infinite population size vs MVP ---------------------
# FS
kapa = 50
percent = 40
FSsa = FSsA=0.005
Ti = 1:kapa
fluct_s <- FSsa*sin(2*pi*Ti/kapa + pi) # selection coeff for genotype AA
fluct_t <- FSsA*sin(2*pi*Ti/kapa)  
fluct_h = 0.5-(percent/100)*sin(2*pi*Ti/kapa) 


FS_AA = (1-fluct_t)/(1+max(FSsa,FSsA))
FS_aa = (1-fluct_s)/(1+max(FSsa,FSsA))
FS_Aa =  (1-fluct_h*fluct_t-(1-fluct_h)*fluct_s)/(1+max(FSsa,FSsA)) 

peq = 0.5 # Equilibrium allele frequency under balancing selection
FSwAA = exp(mean(log(FS_AA)))
FSwAa = exp(mean(log(FS_Aa)))
FSwaa = exp(mean(log(FS_aa)))
WfitFS = FSwAA*peq^2+FSwAa*2*peq*peq+FSwaa*peq^2 # Equilibrium fitness under FS

# HA
hetsa= hetsA=0.005
WfitHA = 1-0.5*hetsa # Equilibrium fitness under HA

# ID or MA
masa = idsa=0.05 # Sel. coeff.
mamut = idmut= 0.001 # Mut. rates
mah=idh=0.0 # Dom. coeff.
if(mah==0){
  mape=min(c(1,1-sqrt(mamut/masa)))
}else{
  mape = min(c(1,1-mamut/(mah*masa)))
}

WfitMA = 1*mape^2+2*mape*(1-mape)*(1-mah*masa)+(1-mape)^2*(1-masa) # Equilibrium fitness under MA or ID


absfit = 1.53*cbind(WfitHA^nloci,WfitFS^nloci, WfitMA^nloci)

loci_c=apply(absfit, 2, FUN = function(x){
  X = which(x<=1)
  if(length(X>0)){return(min(X, na.rm = TRUE))}else{return(length(nloci))}
})

#Plotting absolute fitness at equilibrium with number of loci
par(mgp=c(2.9,1,0),mar=c(5,5,2,2) + 0.1)
matplot(absfit, ty='l', cex.axis = 2, lwd=3, ylab = list("Absolute fitness", cex=2.8, font=4), xlab = list("Number of loci", cex=2.8, font=4))
#abline(h=c(1,1,1), v = nloci[loci_c], col=1:3 )
points(nloci[loci_c],c(1,1,1), col="gray", pch=16, cex=2)
abline(h=1, col="gray",lty=2, lwd=2)
arrows(x0 = nloci[loci_c],y0 = c(1,1,1),x1 = nloci[loci_c], y1 = c(0.15,0.15,0.15), lty = 2, lwd = 2, col = "gray")
legend("topright", legend = c("HA", "FS", "ID"), col=1:3, cex = 2, text.font = 4, lty = 1:3, lwd = 3)
```
