## Supplementary material for "Modeling minimum viable population size with multiple genetic problems of small populations": R codes used to simulate results: ReadMeFirst_MVP_GeneticProblems.pdf

This folder contains the R scripts for simulating the individual-based model and the and analytic approximations for the manuscript "**Modeling minimum viable population size with multiple genetic problems of small population size**". To run any of the files, FIRST CREATE A FOLDER NAMED "MVPGENETICPROBLEMS" in your home folder and place all the *R* files into this folder. The *R* files include:
"MainFunct_IBM.R",
"MainFunct_DiffusionApproximation.R",
"MainFunct_MarkovChain_POPequiApproach_FixedLoci.R",
"MainFunct_MarkovChain_POPequiApproach_VariableLoci.R",
"MVP_GeneticProblems_PLOTS.R", "Execute_DiffusionApproximation_FixedLoci.R",
"Execute_DiffusionApproximation_VariableLoci.R", "Execute_IBM.R",
"Execute_MarkovChain_POPequiApproach_FixedLoci.R", and
"Execute_MarkovChain_POPequiApproach_VariableLoci.R"

The main function for the individual-based model is given in the *R* script ""MainFunct_IBM.R". This script is executed using "Execute_IBM.R" script.

The main function for the Diffusion approximation approach (DAA) is given in the *R* script "MainFunct_DiffusionApproximation.R". This script is executed using two R scripts. The script "Execute_DiffusionApproximation_FixedLoci.R" executes the case with fixed number of loci while the script "Execute_DiffusionApproximation_VariableLoci.R" executes the case with variable number of loci.

The main function for the Markov chain approach (MCA) is coded in two *R* scripts. The first *R* script ""MainFunct_MarkovChain_POPequiApproach_FixedLoci.R" is for the case of fixed number of loci and is executed by "Execute_MarkovChain_POPequiApproach_FixedLoci.R". The second *R* script "MainFunct_MarkovChain_POPequiApproach_VariableLoci.R" is coded for the case of variable number of loci and is executed by "Execute_MarkovChain_POPequiApproach_VariableLoci.R" script.

This script is executed using two *R* scripts. The script "Execute_DiffusionApproximation_FixedLoci.R" executes the case with fixed number of loci while the script "Execute_DiffusionApproximation_VariableLoci.R" executes the case with variable number of loci.

The script "MVP_GeneticProblems_PLOTS.R" is used to visualisze the results. So, all the plots from the Results section and supplementary material can be generated in this script. Note however that in plots were there is need to combine different parameters or different mechanisms, the plots can only be for one given scenario but not combined as in the manuscript.

Within the scripts, all the parameters used are described in the comments. The denotation of some parameters in the scripts differ from those used in the manuscript partly because they are generic functions within R. There are also cases where the definition of parameters differ between scripts, but in such cases, the definition is given in the comment.

Therefore, we strongly advise to read through the description provided within the script so as to have the right parameter combinations. Also, note that there are other parameters not used in our manuscript and are clearly identified via comments. DO NOT CHANGE THE PARAMETER VALUES GIVEN TO SUCH PARAMETERS!

Note that the provided parameter values in the execution scripts may only represent one scenario of parameter combinations used in one of the cases in the manuscript. Also, in the plot script, the axes limits and legends may have to be manually changed to suit the specific parameter combination(s).

WARNING! The actual parameters used to produce the figures in the manuscript may need a lot more time, ranging from hours to weeks even when using a node on cluster with 20 cores! All the execution scripts are parallelized to run on multiple cores. So, it is more suitable to use computer or cluster with more cores.

In the execution scripts provided, we have reduced on parameter values that require much time for purposes of being able to run the scripts on a personal computer even with 2 cores. For example, reduced simulation time "Tmax" from say 4000 to 400 or even less, reduced the length of the mutation rate vector over which the models have to run from say 10 values to 2, reduced the length and values in the carrying capacity vector from say 350 values to only 10 and using lower values say carrying capacity of 50000 to 100, number of replicates from say 60 to only 4, e.t.c. In all cases where this reduction is done, the actual values used in the manuscript are stated in the comment on the immediate line below.

The scripts were individual-based model data was produced on a Linux computing cluster (Ubuntu 16.04.6). The produced data was analysed in Linux operating system (Release: 20.04 ), with *R* version 3.6.3 (2020-02-29) and has not been tested on any other versions.
